# Beyond the Shared Inflammatory Axis: Differentiating Molecular Signatures in Psoriatic Arthritis and Ankylosing Spondylitis through Integrated Omics

**DOI:** 10.1101/2025.08.20.671331

**Authors:** Laís de Carvalho Gonçalves, João Firmino Rodrigues-Neto, Gustavo Antônio de Souza, João Paulo Matos Santos Lima

## Abstract

Spondyloarthropathies (SpA), which include ankylosing spondylitis (AS) and psoriatic arthritis (PsA), are inflammatory rheumatic diseases with overlapping clinical and immunological characteristics but distinct molecular mechanisms. This study aimed to compare transcriptomic signatures of AS and PsA by investigating differentially expressed genes and associated regulatory pathways. Four public RNA-Seq datasets from GEO (GSE186063, GSE117769, GSE205748, and GSE221786) were used. Using an integrative strategy involving scientific text mining, functional enrichment, and biological curation, 433 genes with potential relevance to both conditions were selected. Based on this set, differential gene expression (DEG) analyses were performed, protein-protein interaction (PPI) networks and gene regulatory networks (GRN) were constructed, and hubs and master regulators (MRs) were identified. The integration of these approaches allowed the identification of shared inflammatory pathways, especially related to the IL-23/IL-17 axis, as well as disease-specific signatures. Functional analysis revealed enrichment in Th17 cell differentiation processes, lipopolysaccharide response, and activation of classical inflammatory pathways. This study provides a comprehensive comparative view of the transcriptomic signatures of PsA and AS that point to potential biomarkers and distinct therapeutic targets.

## INTRODUCTION

Psoriatic Arthritis (PsA) and Ankylosing Spondylitis (AS) are chronic inflammatory spondyloarthritides that exhibit clinical, immunological, and genetic overlap; however, they diverge in their phenotypic presentation and underlying pathogenic mechanisms (Feld et al., 2018). These autoimmune conditions have several commonalities, including the involvement of entheses and dactylitis, the presence of various extra-articular manifestations such as uveitis, colitis, inflammatory bowel disease, and cutaneous manifestations, as well as a favorable response to anti-TNF-α and IL-17 inhibiting therapies (Mease et al., 2019). Furthermore, the IL-23/IL-17 pathway is a participant in the inflammatory processes of both diseases (Chimenti et al., 2015). Despite these shared inflammatory pathways, there are distinct clinical and molecular targets observed in PsA and AS, which demand different diagnostic and therapeutic approaches for each condition.

A crucial aspect that differentiates PsA and AS lies in the topography of inflammation. In PsA, peripheral inflammation is characteristic and frequently associated with psoriasis. Its pathogenesis involves a complex interplay of genetic predisposition, including HLA-B27, HLA-Cw6, IL-23R, and ERAP1, alongside environmental factors such as infections, trauma, and smoking. The IL-23/IL-17 pathway plays a particularly prominent role (Veale and Fearon, 2018), where the activation of dendritic cells promotes the release of IL-23, leading to the expansion of Th17 cells and the subsequent production of IL-17A, IL-22, and TNF-α (McGonagle et al., 2015), which are implicated in cutaneous manifestations (Colbert et al., 2010). In contrast, AS is predominantly characterized by axial inflammation, primarily affecting the spine and sacroiliac joints, with a potential for progression to ankylosis (Sieper et al., 2015). Unlike PsA, AS demonstrates a strong association with HLA-B27, present in up to 90% of cases, where altered expression of this gene can lead to dimer formation (Ranganathan et al., 2017). The interaction between HLA-B27, the gut microbiota, and the epithelial barrier is considered a crucial element in the pathogenesis of AS (Rosenbaum & Davey, 2011).

The advent of high-throughput transcriptomics data, particularly RNA-sequencing (RNA-seq), has revolutionized the investigation of the molecular basis of complex diseases (Zhang et al., 2018; Zhang et al., 2021; Wang et al., 2019). The adoption of a systems biology approach, which integrates gene expression analysis with interaction networks and functional pathways, facilitates the construction of more holistic models of disease pathogenesis (Zhang et al., 2023; Zhang et al., 2018; Vidal, Cusick & Barabási, 2011; Marbach et al., 2012). By analyzing and comparing the transcriptomic profiles of patients with AS and PsA, it becomes possible to identify not only genes with altered expression but also key regulatory “hubs” and the specific biological pathways that govern inflammation and tissue damage in these conditions (Zhang et al., 2022; Dey-Rao and Sinha, 2017; Guimarães et al., 2021).

The present study aimed to identify differentially expressed genes in EA and APs, and to analyze the differences and similarities in gene regulatory networks (GRNs) and specific master regulators (MRs) that control the expression of multiple genes involved in each disease. To this end, a set of 433 genes, selected through scientific text mining and functional enrichment, was used as the basis for further analysis. In addition, this work presents a discussion on the functional relevance of these molecular agents in pathogenesis, offering a deeper understanding of common and specific molecular pathways in EA and APs, thus identifying potential therapeutic and predictive targets for these autoimmune diseases.

## METHODOLOGY

The methodological framework for this comparative genomic analysis between Psoriatic Arthritis (PsA) and Ankylosing Spondylitis (AS) involved a systematic, multi-step approach, as summarized in Figure 1.

**Figure 1.**
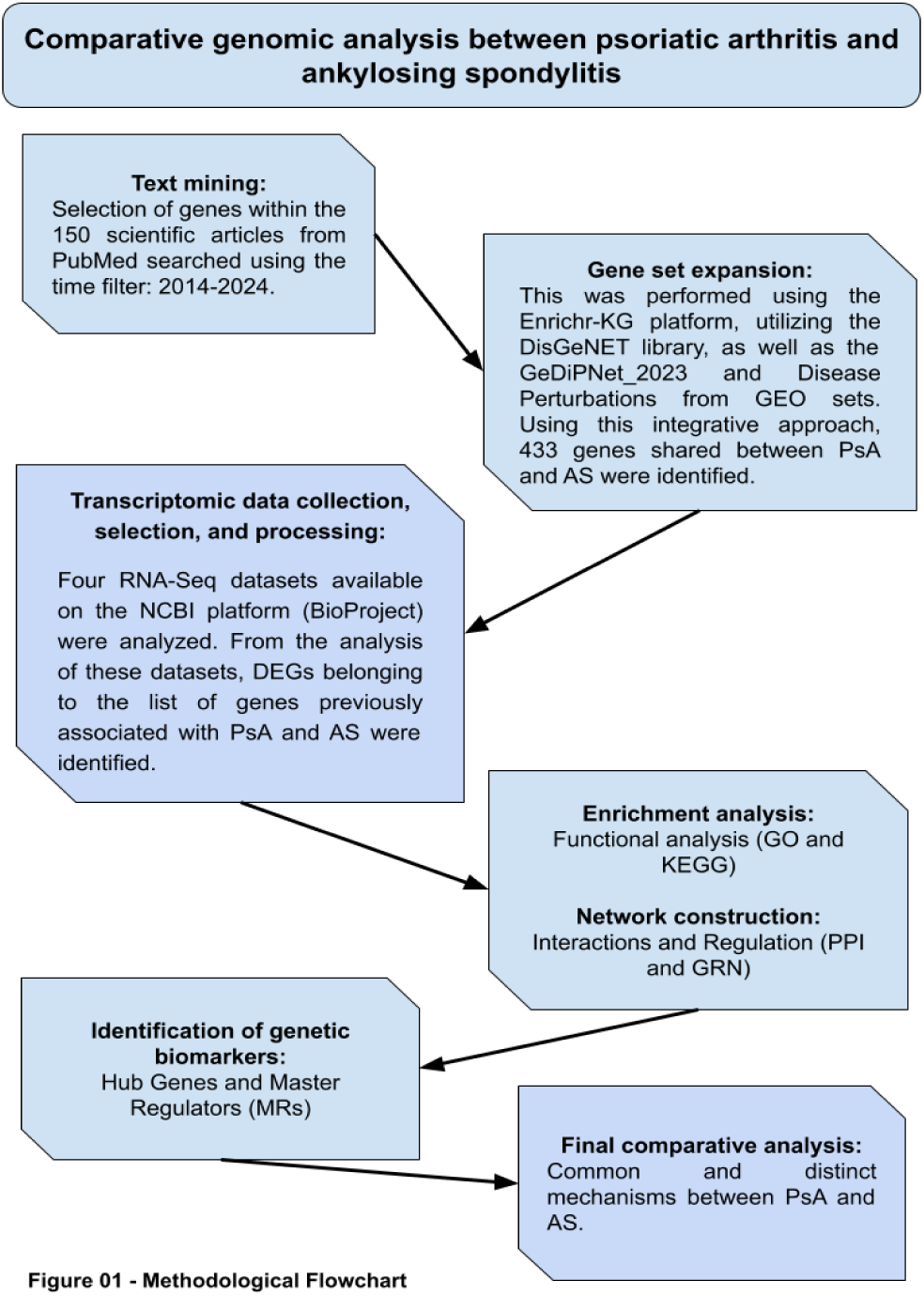
Methodological Flowchart of Comparative Genomic Analysis between Psoriatic Arthritis and Ankylosing Spondylitis. The flowchart illustrates the methodological steps employed for the comparative genomic analysis between psoriatic arthritis (PsA) and ankylosing spondylitis (AS). The study began with text mining of scientific articles from PubMed (2014-2024) to select relevant genes. Subsequently, transcriptomic data were collected from four RNA-Seq datasets from the NCBI BioProject, focusing on genes previously associated with PsA and AS. The gene list was expanded using the Enrichr-EG platform and libraries such as DisGeNET, as well as the GeDiPNet_2023 and Disease Perturbations from GEO datasets, identifying 433 genes shared between the two conditions. Subsequent analyses included functional enrichment (GO and KEGG), the construction of protein-protein interaction (PPI) networks, and gene regulatory networks (GRN). The ultimate goal was the identification of genetic biomarkers (hub genes and master regulators) and the performance of a comprehensive comparative analysis of common and distinct mechanisms between PsA and AS.

### Literature Text Mining

A systematic literature search using the PubMed database (https://pubmed.ncbi.nlm.nih.gov/) was first conducted. Separate queries were performed using the descriptors “Psoriatic Arthritis” (PA) and “Ankylosing Spondylitis” (AS). A temporal filter was applied to restrict results to the period between 2014 and 2024, which yielded 150 scientific articles deemed relevant for the investigation. From these selected studies, all genes mentioned in relevant contexts were meticulously cataloged. Concurrently, complementary filtering was performed to expand and refine the set of genes common to both pathologies, resulting in a total of 433 selected genes. This step was performed with the aid of the Enrichr-KG platform (Evangelista et al., 2023), using the DisGeNET, GeDiPNet_2023, and Disease Perturbations from GEO libraries as a basis, including positively and negatively regulated genes. This multifaceted approach to gene selection, performed before transcriptomic data analysis, ensured an initial set based on scientific literature, increasing the biological relevance and interpretive robustness of subsequent findings.

### Transcriptomic Data Selection and Processing

The datasets used in this study were carefully selected from the NCBI BioProject platform, applying specific filters: “transcriptome,” “GEO Datasets,” “Expression profiling by high throughput sequencing,” and “*Homo sapiens*.” Initial searches were conducted individually for each pathology, followed by combined searches across all pathologies (Table 1). All analyses were performed within the R 4.4.1 environment, using the raw count matrices available for each series in GEO, in conjunction with the Annotation Release 109 (2019-09-05) of the GRCh38.p13 assembly, provided by NCBI RefSeq. All analyzed datasets corresponded to RNA-Seq transcriptomics experiments and underwent processing, normalization, and analysis using the R packages GEOquery (Davis et al., 2007), DESeq2 (Love et al., 2014), edgeR (Robinson et al., 2010), and dplyr (Wickham et al., 2023).

**Table 1:** Characteristics of Publicly Available Datasets Used in the Study.

| **GEO Dataset** | **Reference** | **Sequencing Platform** | **Short description** | **Associated Metadata** | **Samples used and analyzed in the present study** |
| --- | --- | --- | --- | --- | --- |
| GSE186063 | Deng et al., 2022 | Illumina NovaSeq 6000 | The study comprises skin samples categorized into five distinct groups: 13 patients with psoriatic lesions (Pso), 13 patients with non-lesional Pso skin, 13 patients with lesional PsA skin, 15 patients with non-lesional PsA skin, and 12 patients with AS non-lesional skin, the latter serving as a control group for specific analyses. | Diagnosis, age, sex, tissue type, and clinical status of participants | Only the PsA (lesional and non-lesional) and AS groups were used, allowing for an understanding of cutaneous inflammatory processes associated with both pathologies. |
| GSE117769 | Zhang L. and Li Z., 2019 | Illumina HiSeq 2500 | The study consists of peripheral blood samples from 51 patients with Rheumatoid Arthritis (RA), 49 healthy individuals, 11 patients with PsA, and 8 with AS. | Sex, age, phenotype, origin, cell type, disease activity, and medication use (such as NSAIDs, steroids, immunomodulators, and biological therapies) | Only samples from healthy individuals, patients with PsA, and patients with AS were selected. |
| GSE205748 | JJohnsson et al., 2023, 2024 | Illumina NovaSeq 6000 | Skin biopsies from three groups: nine healthy volunteers without psoriasis or any inflammatory disease, nine samples of skin taken directly from psoriatic plaques of PsA patients, and nine samples of healthy-appearing skin taken from an unaffected area of the same PsA patients. | Tissue type, experimental group. | All samples |
| GSE221786 | Lee et al., 2023 | Illumina NextSeq 500 platform | Comprises peripheral blood samples from 19 AS patients and eight healthy individuals. Peripheral blood mononuclear cells (PBMCs) were isolated and stimulated in vitro with CytoStim for four hours, aiming to enhance the activation of the interleukin-17 (IL-17) mediated pathway, before total RNA isolation and subsequent sequencing. | Sex, disease status, tissue type, and treatment conditions. | All Samples |

*This table provides a comprehensive overview of the Gene Expression Omnibus (GEO) datasets utilized in this study, detailing their references, sequencing platforms, brief descriptions of their contents, associated metadata, and the specific samples selected for analysis from each dataset.*

### Gene Expression Data Collection and Pre-processing

Gene expression data from patients with AS and PsA were retrieved from public databases, primarily the Gene Expression Omnibus (GEO). Dataset selection adhered to stringent criteria, including the availability of RNASeq data, the presence of appropriate control groups, high read quality, sufficient sample size, and direct relevance to the diseases under investigation. Following data acquisition, a comprehensive pre-processing pipeline was implemented, which encompassed normalization, noise filtering, and, where applicable, the removal of batch effects. The identification of differentially expressed genes (DEGs) between disease and control groups was performed using established statistical packages in R, such as DESeq2, applying a rigorous threshold of an adjusted p-value (p<0.05) and a Log2 Fold Change (Log2FC) greater than 1 or less than -1. These stringent criteria ensure that only statistically robust and biologically meaningful changes in gene expression are considered, thereby increasing the reliability of subsequent analyses.

### GO and KEGG Pathway Enrichment Analysis

Functional enrichment analysis was conducted using the clusterProfiler R package (Yu et al., 2012), leveraging the Kyoto Encyclopedia of Genes and Genomes (KEGG) and Gene Ontology (GO) databases. For KEGG enrichment, the KEGGREST (Tenenbaum, 2024) and pathview (Lou and Brouwer, 2013) packages were employed, while topGO and GOstats were utilized for Gene Ontology analysis. (Gene Ontology Consortium, 2021).

### Protein-Protein Interaction (PPI) Network Construction

The identified DEGs were used as input for constructing a protein-protein interaction (PPI) network, utilizing the STRING database (Szklarczyk et al., 2021). The PPI network was built by connecting DEGs that possess known or predicted direct physical interactions. To minimize the inclusion of false positives, only interactions with a moderate confidence score (threshold = 400) were considered. Topological analysis of the PPI network was performed using visNetwork (Almende et al., 2024), which facilitated the identification of highly connected nodes, often indicative of key functional proteins.

### Identification of Gene Hubs

Within the constructed protein-protein interaction (PPI) network, gene hubs were identified based on classical centrality metrics, which reflect their structural and functional importance within the network’s topology. The metrics employed included: (i) degree, representing the total number of direct interactions a node possesses, encompassing both incoming and outgoing connections; and (ii) betweenness centrality, which quantifies the frequency with which a node lies on the shortest paths between other nodes, thereby suggesting a mediating functional role. In parallel, master regulators were identified, defined as transcription factors (TFs) exhibiting a high out-degree in directed networks. This dual approach, distinguishing between structural hubs and master regulators, provides a comprehensive perspective on both the global connectivity and the hierarchical functional organization of the regulatory network. The selection of transcription factors that regulate the DEGs was analyzed using various R packages, including GenomicFeatures (Lawrence et al., 2013), TxDb.Hsapiens.UCSC.hg38.knownGene/org.Hs.eg.db (Carlson, 2024), JASPAR2020 (Fernández-Blanco et al., 2020), TFBSTools (Tan and Lenhard, 2016), and SummarizedExperiment (Huber et al., 2015).

### Gene Regulatory Network (GRN) Construction and Master Regulator (MR) Identification

The Gene Regulatory Network (GRN) was constructed to infer transcriptional control relationships between transcription factors (TFs) and their target genes. Inference tools based on gene expression data and information from established TF-gene interaction databases, such as TRRUST (Transcriptional Regulatory Relationships Unraveled by Sentence-based Text mining) (Han et al., 2018) or ENCODE (Encyclopedia of DNA Elements) (ENCODE Project Consortium, 2020), were utilized. Transcription factors were identified by comparing the experimental groups. They were subsequently summarized in a table detailing the TFs, their regulated genes, associated motifs, number of occurrences, and potential biological functions. This table served as the basis for inferring the gene regulatory network and for confirming hub genes, as well as identifying master regulators. Master Regulators (MRs) were identified within the GRN using algorithms such as ggraph (Pedersen, 2022) and visNetwork. These algorithms infer the activity of transcription factors based on the expression of their target genes, even if the expression of the TF itself is not significantly altered. TFs were defined as MRs if their regulatory activity significantly impacted more than 5 DEGs within the network, indicating their central role in coordinating the disease’s transcriptional response.

### Integration and Comparative Analysis

The results obtained for AS and PsA were integrated and subjected to a comparative analysis to identify both similarities and differences in their molecular profiles. The overlap of gene hubs, master regulators (MRs), and enriched pathways was meticulously analyzed to highlight standard mechanisms and the distinctive characteristics of each disease. Network visualizations and enrichment results were generated using tools such as visNetwork and specific graphics created with the ggplot2 package (Wickham, 2016).

## RESULTS

To identify key genes involved in Psoriatic Arthritis (PsA) and Ankylosing Spondylitis (AS), a systematic literature mining was conducted in the PubMed database, covering the period from 2014 to 2024. The initial analysis focused on the first 150 selected articles, which were complemented by a functional enrichment analysis using the Enrichr-KG platform. This comprehensive approach led to the identification of 3,660 genes associated with any of the two diseases, of which 433 were found to be shared between both conditions (Supplementary Table 1). This curated set of 433 shared genes served as a refined starting point for subsequent transcriptomic analyses, focusing the investigation on the common molecular ground between PsA and AS. Among the genes of highest relevance to both PsA and AS, HLA-B27 and HLA-C (particularly the 06:02 allele) stand out as fundamental for AS susceptibility and its involvement in axial PsA (Ranganathan et al., 2017; Bowes et al., 2015; Brown & Wordsworth, 2016). Genes within the IL-23/IL-17 axis, such as IL23R and IL12B, were consistently implicated in the inflammation observed in both diseases, underscoring their importance as therapeutic targets (Chimenti et al., 2015; Martin et al., 2019; Raychaudhuri et al., 2017; Baliwag et al., 2015). Additionally, ERAP1 was noted for its significant interaction with HLA-B27 in AS, influencing antigen processing. Functional analysis of the shared gene set revealed substantial enrichment in KEGG pathways related to *immune response*, *inflammatory signaling*, and *Th17 cell differentiation*. Concurrently, Gene Ontology (GO) biological processes, such as “*response to cytokines*,” “*regulation of adaptive immune response,*” and “*leukocyte migration*,” were also highly enriched.

### Transcriptomic Dataset Selection and Analysis

This study meticulously selected four public transcriptomic datasets from the Gene Expression Omnibus (GEO) database, maintained by the NCBI platform: GSE186063 (Deng et al., 2022), GSE117769 (Zhang L. and Li Z., 2019), GSE205748 (Johnsson et al., 2023, 2024), and GSE221786 (Lee et al., 2023). The selection criteria prioritized datasets obtained exclusively via RNA-Seq, ensuring a minimally adequate number of samples and clinical relevance for investigating Psoriatic Arthritis (PsA) and Ankylosing Spondylitis (AS). This strategic dataset selection, encompassing both skin and peripheral blood samples, was designed to provide insights into both localized (tissue-specific) and systemic inflammatory profiles of the diseases. Table 1 lists the details and features of each dataset and the samples used for subsequent analyses.

### Differentially Expressed Genes (DEGs) in AS and PsA with and without skin lesions

The 433 genes identified in the shared list served as the foundation for subsequent transcriptomic analyses (Supplementary Table 1). In the GSE186063 dataset, clustering analysis of the twenty most differentially expressed genes accurately segregated samples into lesional PsA, non-lesional PsA, and non-lesional AS, with the latter two conditions forming a single cluster (Supplementary Figure 1). A specific comparison between non-lesional skin samples from patients with Psoriatic Arthritis (PsA) and Ankylosing Spondylitis (AS) samples from the same dataset yielded no differentially expressed genes (DEGs) when applying the predefined adjusted p-value and log2FC criteria. This lack of DEGs suggests that at a basal, clinically non-inflamed state of the skin, the molecular profiles of these conditions, or at least the shared gene set, are remarkably similar to each other. In contrast, a markedly distinct expression profile was observed when comparing non-lesional skin with lesional skin from PsA patients (Figure 2), resulting in the identification of 121 DEGs from the shared gene list. In this analysis, non-lesional skin exhibited overexpression of 38 genes and robust underexpression of 83 genes.

**Figure 2.**
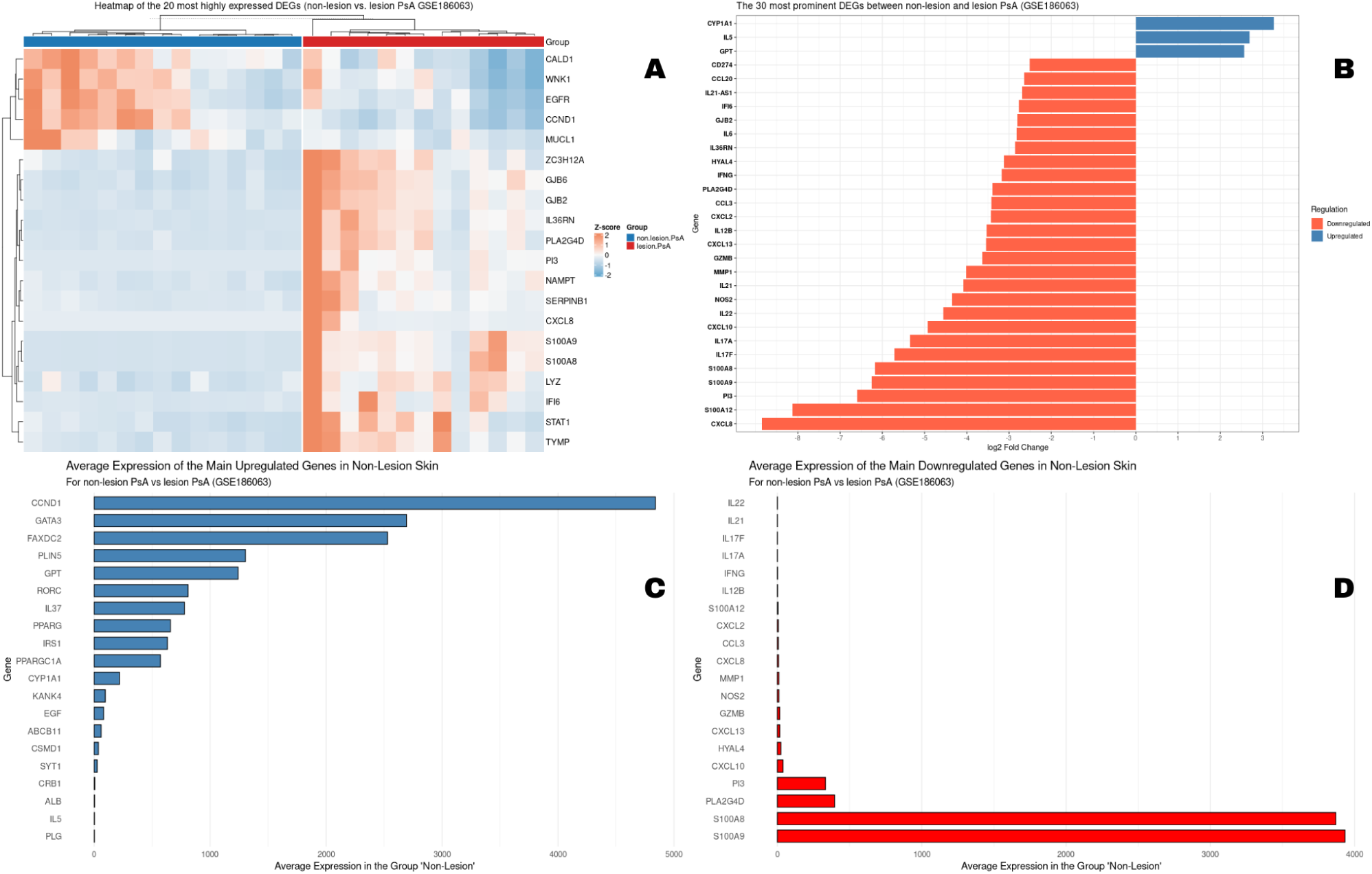
Differential Expression Profile in Psoriatic Arthritis (PsA) Non-Lesional and Lesional Skin Reveals Immune Suppression and Dissociation between Magnitude of Change and Expression Levels. Analysis of the GSE186063 dataset comparing non-lesional vs. lesional skin tissue. (A) Heatmap of the 20 genes with the highest average expression, showing the relative expression pattern (Z-score) that separates non-lesional (blue) and lesional (red) samples. (B) Bar chart showing the Log2FC of the 30 most prominent differentially expressed genes (DEGs). Genes with increased expression in non-lesional skin are in blue, and genes with decreased expression are in orange. (C) Average expression level of the main overexpressed genes in the comparison. (D) Average expression level of the main underexpressed genes.

A heatmap of the 20 genes with the highest differential expression clearly showed a segregation between the two sample groups, reflecting distinct and consistent transcriptional profiles (Figure 2A). Analysis of the top 30 differentially expressed genes highlighted a predominance of underexpressed genes in non-lesions, such as CXCL8 and S100A12, which displayed the lowest Log2FC values (Figure 2B). Interestingly, a dissociation between the magnitude of relative variation (Log2FC) and absolute expression levels was noted. While CCND1 was among the most abundant overexpressed genes in non-lesional tissue (Figure 2C), IL22 and IL21 were the least abundant genes in non-lesional PsA skin (Figure 2D).

The analysis of differential gene expression between normal skin from AS patients and lesional skin from PsA patients revealed a profoundly asymmetric and biologically relevant transcriptional landscape (Figure 3), leading to the identification of 129 DEGs, with 39 upregulated and 89 downregulated. Evaluation of the 20 most differentially expressed genes showed a clear separation between the two sample groups, reflecting distinctly modulated basal transcriptional profiles (Figure 3A). The analysis further indicated an asymmetric pattern, with a predominance of significantly underexpressed genes in AS skin compared to PsA lesions (Figure 3B), such as CXCL8, S100A12, and PI3, which exhibited intense transcriptional repression (log2FC <-1). Conversely, only a limited number of genes showed overexpression in AS skin, including CYP1A1 and ALB among the 30 most differentially expressed genes. Similar to the previous paired analysis, a dissociation between log2FC and transcriptional abundance was observed. For instance, MUCL1 and CCND1 stood out as the most abundant overexpressed genes in non-lesional skin of AS patients (Figure 3C), while IL21 and IL22 showed low basal expression and were strongly repressed in the context of this disease (Figure 3D).

**Figure 3.**
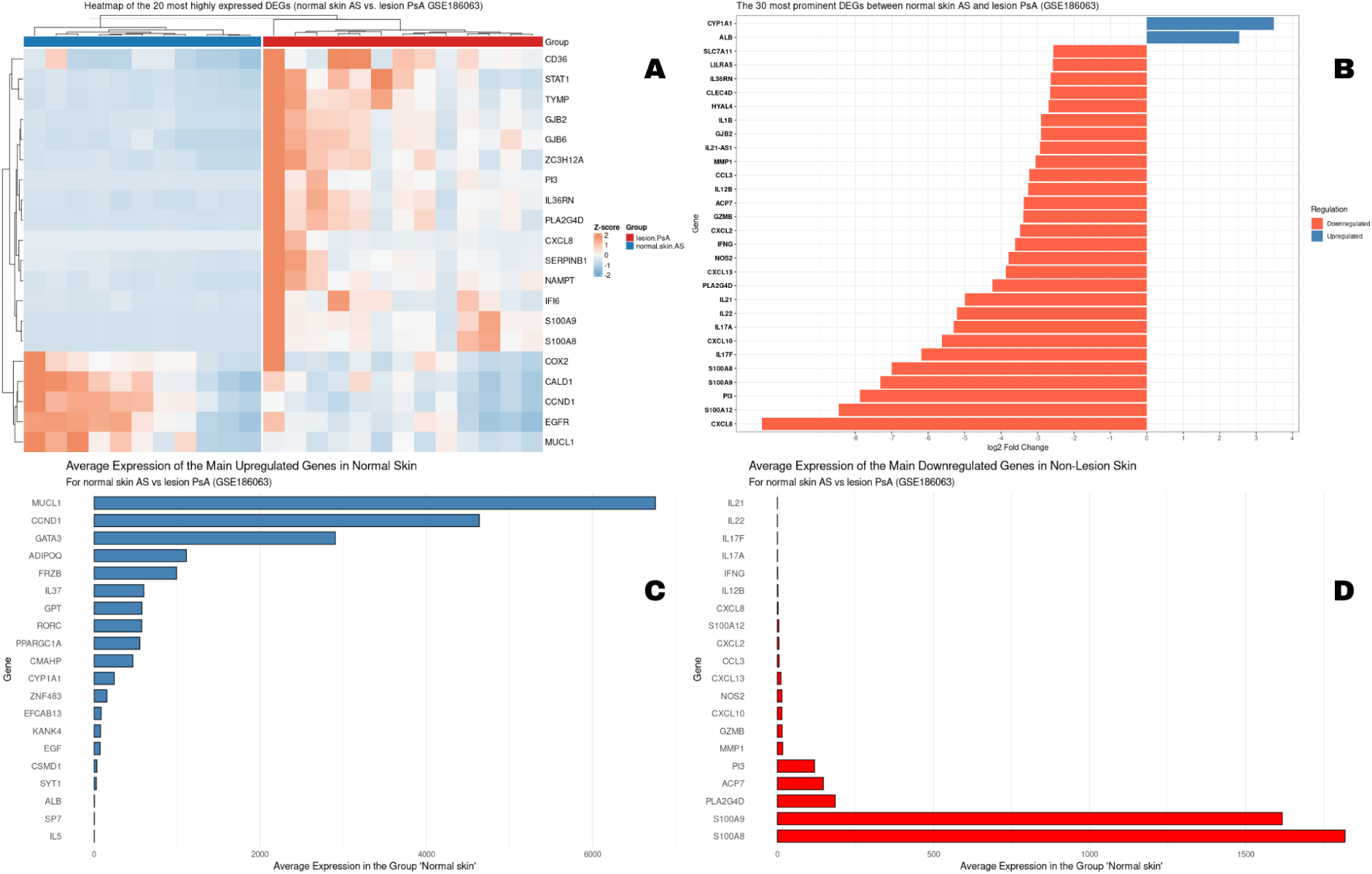
Comparative Analysis of Transcriptional Profile between Normal Skin of Ankylosing Spondylitis (AS) Patients and Psoriatic Arthritis (PsA) Lesions. The analysis was performed using RNA-Seq data from the GSE186063 dataset. (A) Heatmap of the expression of the 20 genes with the highest average expression in the dataset. Expression levels are represented as Z-score, with red indicating expression above the gene’s average and blue, below the average. Samples (columns) clearly separate by group (lesion.PsA and normal.skin.AS). (B) Bar chart of the Log2 Fold Change of the 30 most prominent differentially expressed genes (DEGs) (adjusted p-value < 0.05) in the comparison between AS skin and PsA lesion. Blue bars indicate upregulation and orange bars indicate downregulation in AS skin. (C) Average expression level in normalized counts of the main overexpressed genes in AS skin. (D) Average expression level of the main underexpressed genes in AS skin.

From the integration of these analyses, a standard set of genes was identified that distinguishes the shared basal state of non-lesional PsA skin and AS skin from the active inflammatory profile of psoriatic lesions. This set includes 33 upregulated and 75 downregulated genes, respectively (Supplementary Table 2).

Despite the smaller number of PsA and AS samples, the GSE117769 dataset enabled the comparison of gene expression from the initial list between patients with these diseases and healthy controls (Figure 4). The comparison between PsA patients and the control group (Figure 4A) revealed a modest alteration profile, with only seven differentially expressed genes. PRTN3, OLR1, and CLEC4D were identified as overexpressed, while DUSP4, TNFAIP3, CD69, and IL6 were underexpressed. The analysis between AS and controls identified 20 differentially expressed genes (Figure 4B). Notably, CCL2, MYC, CCR6, CCR7, and RORC were overexpressed, while CXCL8, DDIT3, GJB6, IFNG, FOS, JUNB, and ZFP36 were underexpressed. The direct comparison between the two diseases revealed a remarkable molecular similarity, with only five differentially expressed genes (Figure 4C). PRTN3, CLEC4D, MMP9, and RETN were underexpressed in AS, while CCR6 was overexpressed in AS and underexpressed in PsA. Once again, the differential expression of genes from the shared list clustered patients according to their pathological condition.

**Figure 04.**
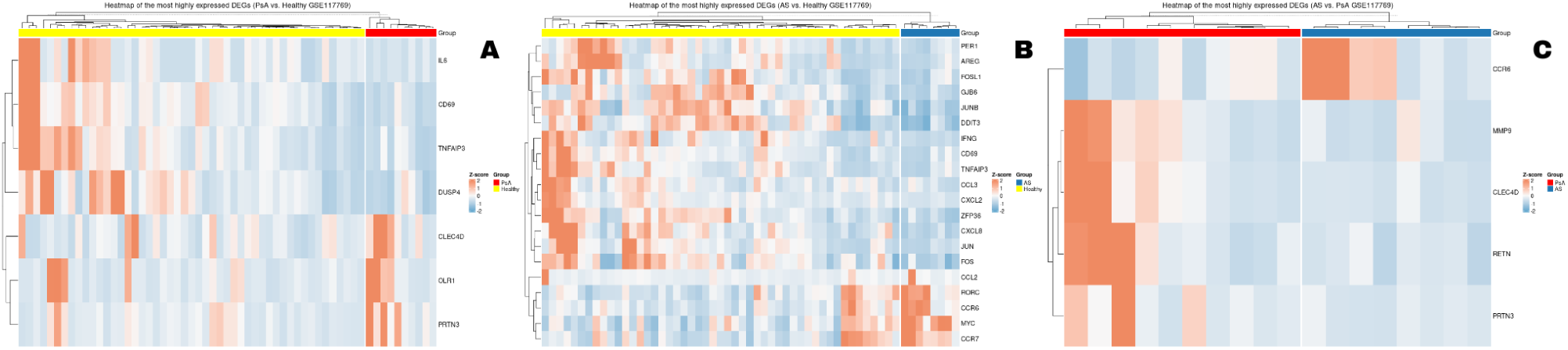
Comparative Gene Expression Analysis by Heatmaps in Study GSE117769 between Psoriatic Arthritis (PsA), Ankylosing Spondylitis (AS), and Healthy Controls. The figure presents three heatmaps, each illustrating the expression profile of the most prominent differentially expressed genes (DEGs) for a specific comparison. In all panels, rows represent genes and columns represent samples, both of which are organized by hierarchical clustering. The color scale indicates relative gene expression (red/orange: high; blue: low). (A) Comparison between PsA (red) and healthy controls (yellow), demonstrating a more focused profile, with a smaller number of genes. (B) Comparison between AS (blue) and healthy controls (yellow), revealing a complex and robust transcriptional signature that clearly distinguishes the two groups. (C) Direct comparison between PsA (red) and AS (blue), showing a subtle molecular difference, with a minimal set of genes distinguishing the two pathologies.

Analysis of the GSE205748 dataset revealed distinct gene expression profiles when comparing skin samples from patients with Psoriatic Arthritis (PsA) and healthy controls. The comparison between clinically non-lesional skin from PsA patients and control skin identified only one differentially expressed gene: SPP1 (osteopontin), which was found to be overexpressed. This finding, despite the absence of visible clinical alterations, suggests the presence of subtle molecular modifications in the cutaneous microenvironment, potentially reflecting a state of pre-activation or subclinical inflammation. In contrast, analysis of lesional PsA skin versus controls demonstrated a robust inflammatory profile, with 94 genes overexpressed, including CD8A, CTLA4, STAT1, STAT3, S100A8/9, and PPARGC1B, and 55 genes underexpressed, such as FOS, GATA3, IL18, PPARGC1A, and TGFBR3 (Figure 5).

**Figure 5.**
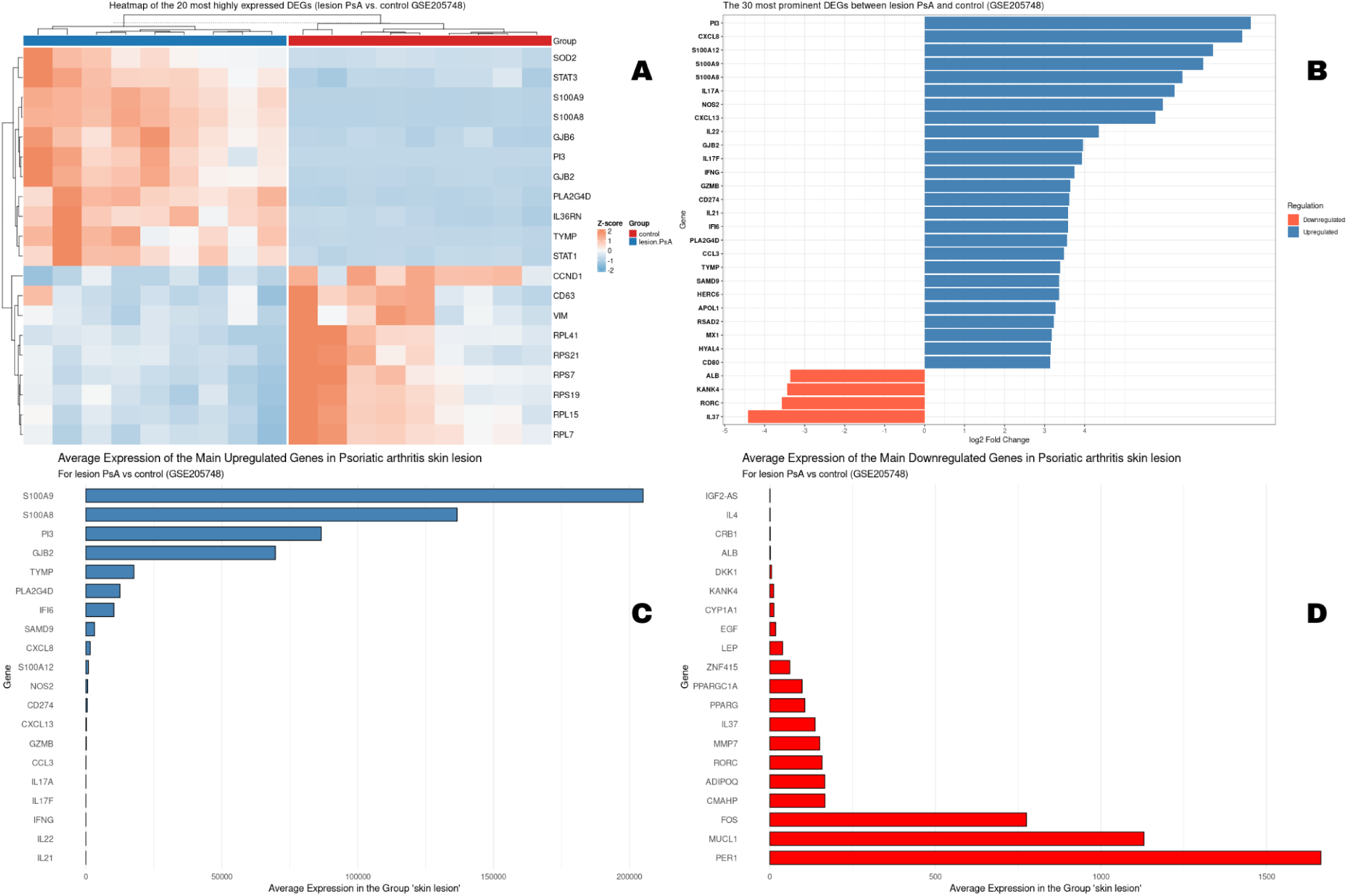
Analysis of Differential Gene Expression between Psoriatic Arthritis (PsA) Lesional Tissue and Controls in Study GSE205748. The figure presents a multifaceted analysis of the main differentially expressed genes (DEGs). (A) Heatmap of the average expression of the 20 most prominent DEGs, with hierarchical clustering of genes (rows) and samples (columns), clearly separating the PsA and Control groups. Colors indicate relative expression (red/orange: high; blue: low). (B) Bar chart of the 20 most significant DEGs, classified by their log2 fold change (L2FC), showing the magnitude of deregulation. Blue bars represent upregulated genes and red bars, downregulated genes in PsA lesions. (C) Average expression levels of the main overexpressed genes. (D) Average expression levels of the main underexpressed genes.

A direct comparison between non-lesional and lesional skin from the same PsA patients confirmed this dichotomy, identifying 63 genes that were overexpressed and 101 genes that were underexpressed in the non-lesional tissue. Non-lesional tissue exhibited higher expression of genes such as EGF, FOS, GATA3, IL1A, PPARG, RORC, and TGFBR3, while showing repression of genes like STAT1, MX1, GBP1, FOXP3, PDCD1, IL36RN, and NOD2 (Figure 6).

**Figure 6.**
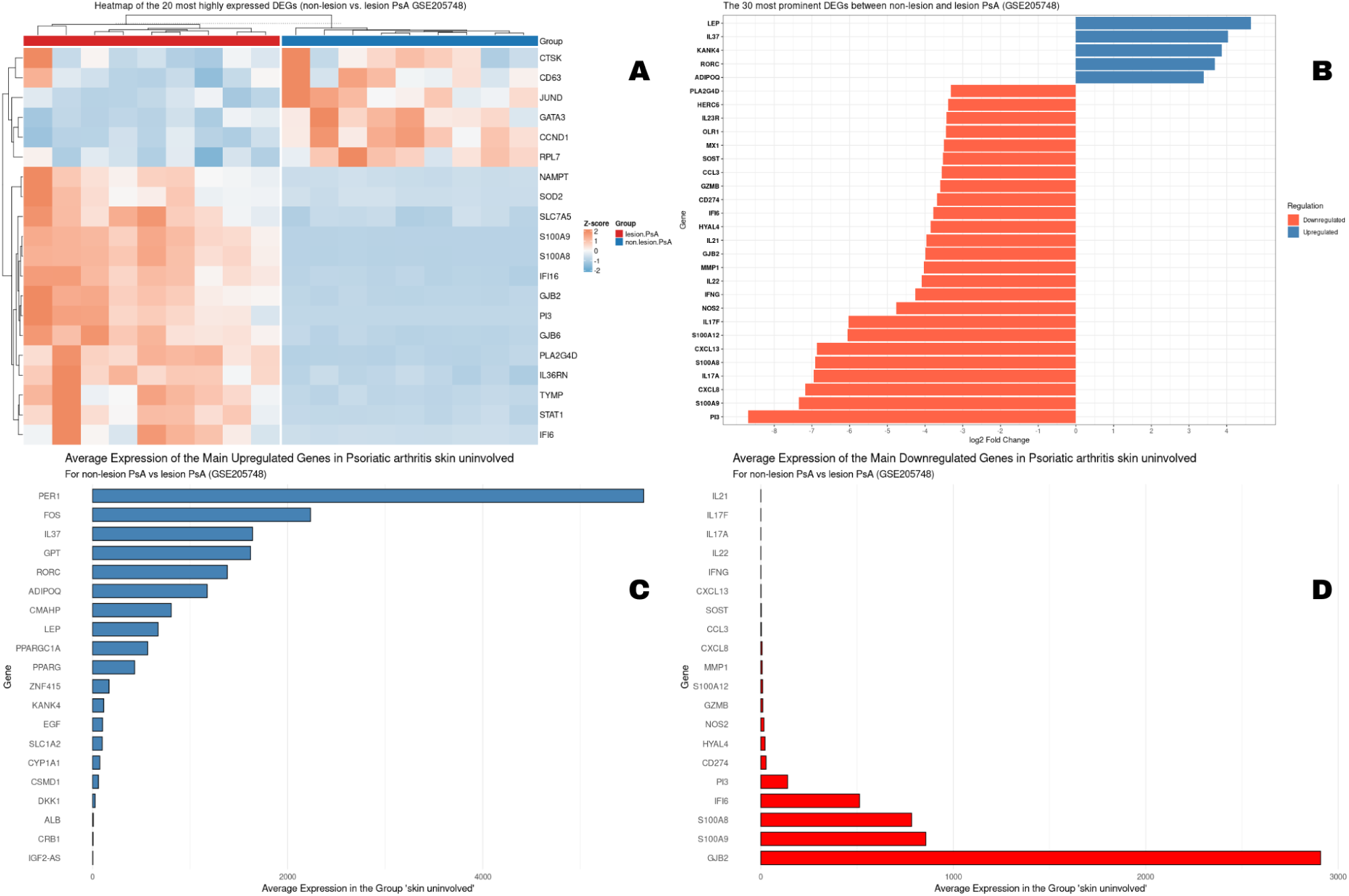
Characterization of Transcriptional Signature in the Transition from Non-Lesional to Lesional Tissue in Psoriatic Arthritis (GSE205748). The figure presents a detailed analysis of differentially expressed genes (DEGs) between lesional and non-lesional skin in patients with PsA. (A) Heatmap of the average expression of the 20 most prominent DEGs, demonstrating a clear separation between groups through hierarchical clustering of genes (rows) and samples (columns). Colors represent relative gene expression (orange: high; blue: low). (B) Bar chart of the 20 most significant DEGs classified by their log2 fold change (L2FC). Blue bars indicate upregulated genes, and orange/red bars indicate downregulated genes in non-lesional tissue. (C) Average expression levels of the main overexpressed genes in non-lesional tissue. (D) Average expression levels of the main underexpressed genes in non-lesional tissue.

In the GSE221786 dataset, only one gene from the initial list, DYSF, showed differential expression, being overexpressed in individuals with AS when compared to healthy controls.

### Protein-Protein Interaction (PPI) Network Analysis and Hub Gene Identification

To investigate the functional relationships among the differentially expressed genes (DEGs) identified in each analysis, protein-protein interaction (PPI) networks were constructed (Figure 07). Comparative study of these networks revealed considerable heterogeneity in their complexity and density of interactions. Comparisons involving non-lesional psoriatic arthritis (PsA) tissue against lesional tissue, AS patient tissue against lesional PsA tissue (from GSE186063), non-lesional PsA tissue against lesional PsA tissue, and lesional PsA tissue against control (from GSE205748) resulted in extensive and highly interconnected networks. This pattern suggests a profound perturbation of the transcriptome, with a large number of interacting proteins driving the active pathology within the lesion. Conversely, networks generated from comparisons between PsA and AS, AS and healthy controls, or PsA and healthy controls within the GSE117769 dataset were considerably smaller and more sparse.

Topological analysis of the PPI networks derived from the GSE186063 dataset elucidated the intricate molecular architecture underlying psoriatic lesions. In both the comparison between non-lesional and lesional PsA tissue (Figure 7A) and the comparison between normal Ankylosing Spondylitis (AS) skin and PsA lesions (Figure 7B), the networks demonstrated a high density of connections In both analyses, genes such as STAT1, IL6, CXCL8, IL1B, and TNF consistently emerged as high-centrality hub nodes. These findings underscore the role of these genes as primary orchestrators of inflammatory and proliferative pathways, which are fundamental to the pathogenesis and exacerbated phenotype of psoriatic lesions.

**Figure 07.**
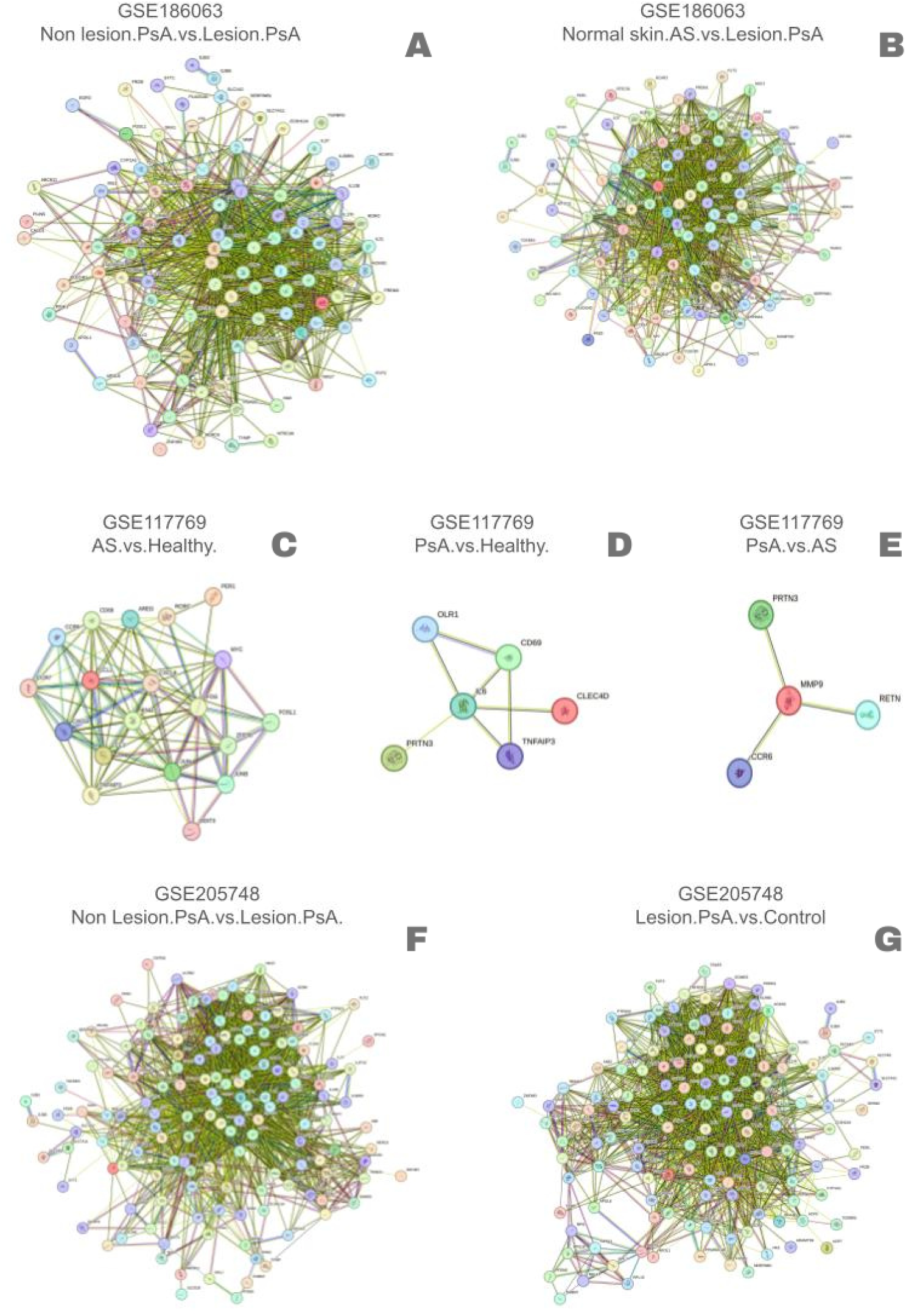
Protein-Protein Interaction (PPI) Networks of Differentially Expressed Genes in Multiple Cohorts and Pathological Conditions. The figure displays seven distinct PPI networks, each constructed from lists of differentially expressed genes (DEGs) derived from three transcriptomic datasets (GSE186063, GSE117769, GSE205748). The networks were generated using the STRING database to visualize predicted functional interactions between proteins encoded by the DEGs. In each panel, nodes represent proteins and edges (lines) indicate interactions. The density and size of each network reflect the number of DEGs and the connectivity among them for the specific comparison, as shown in the title of each panel, allowing for a visual assessment of the molecular complexity of each condition.

Comparative analysis of the PPI networks for the GSE117769 dataset revealed markedly distinct molecular signatures across PsA, AS, and healthy control conditions. The comparison between AS and healthy controls generated the most complex and densely connected network, indicating a systemic perturbation (Figure 7C). Key interactions within this network were orchestrated by multiple hubs, including the cytokine IFNG, the FOS/JUN complex, and various chemokines such as CXCL8 and CCL2. In stark contrast, the network comparing PsA with healthy controls was considerably more sparse, organized around a single central hub, IL6 (Figure 7D). Even more striking was the minimal network resulting from the direct comparison between PsA and AS, which was reduced to a single hub, MMP9, interacting with only a few partners (Figure 7E).

Analysis of the PPI networks in the GSE205748 dataset highlighted the molecular complexity involved in Psoriatic Arthritis (PsA), revealing a consistent pattern of dysregulation in both lesional and non-lesional skin areas. Comparisons between non-lesional and lesional skin (Figure 7F), as well as between lesional skin and healthy controls (Figure 7G), resulted in dense and highly interconnected networks. This reflects an extensive activation of immunological and inflammatory pathways. Proteins such as STAT1, IL6, TNF, S100A8, and S100A9 emerged as central hubs in these networks, reinforcing their critical role in mediating the characteristic inflammatory phenotype of PsA. These findings suggest that, even in clinically unaffected regions, cutaneous tissue already exhibits a latent pro-inflammatory profile, predisposing it to lesion manifestation under additional stimuli.

### GO and KEGG Pathway Enrichment Analysis

A functional enrichment analysis using Gene Ontology (GO) terms and KEGG pathways was conducted on the DEGs from the datasets to identify the most relevant biological processes and molecular pathways in the different clinical contexts of the two diseases. The GO analysis of the GSE186063 dataset revealed that in both comparisons, PsA non-lesional vs. lesional tissue (Figure 8A) and AS normal skin vs. PsA lesional tissue (Figure 8B), the DEGs were predominantly enriched for biological processes such as “*Inflammatory response*,” “*Response to lipopolysaccharid*e,” “*Response to bacterium*,” and “*Immune response*,” thereby confirming the immunomediated nature of the pathology. More specifically, KEGG pathway analysis identified a remarkably consistent set of pathogenic mechanisms across both comparisons (Figure 8C and 8D). The “*IL-17 signaling pathway*,” “*Inflammatory bowel diseas*e,” “*Cytokine-cytokine receptor interaction*,” and “*Rheumatoid arthritis*” consistently emerged as the most significantly enriched pathways in both scenarios.

**Figure 08.**
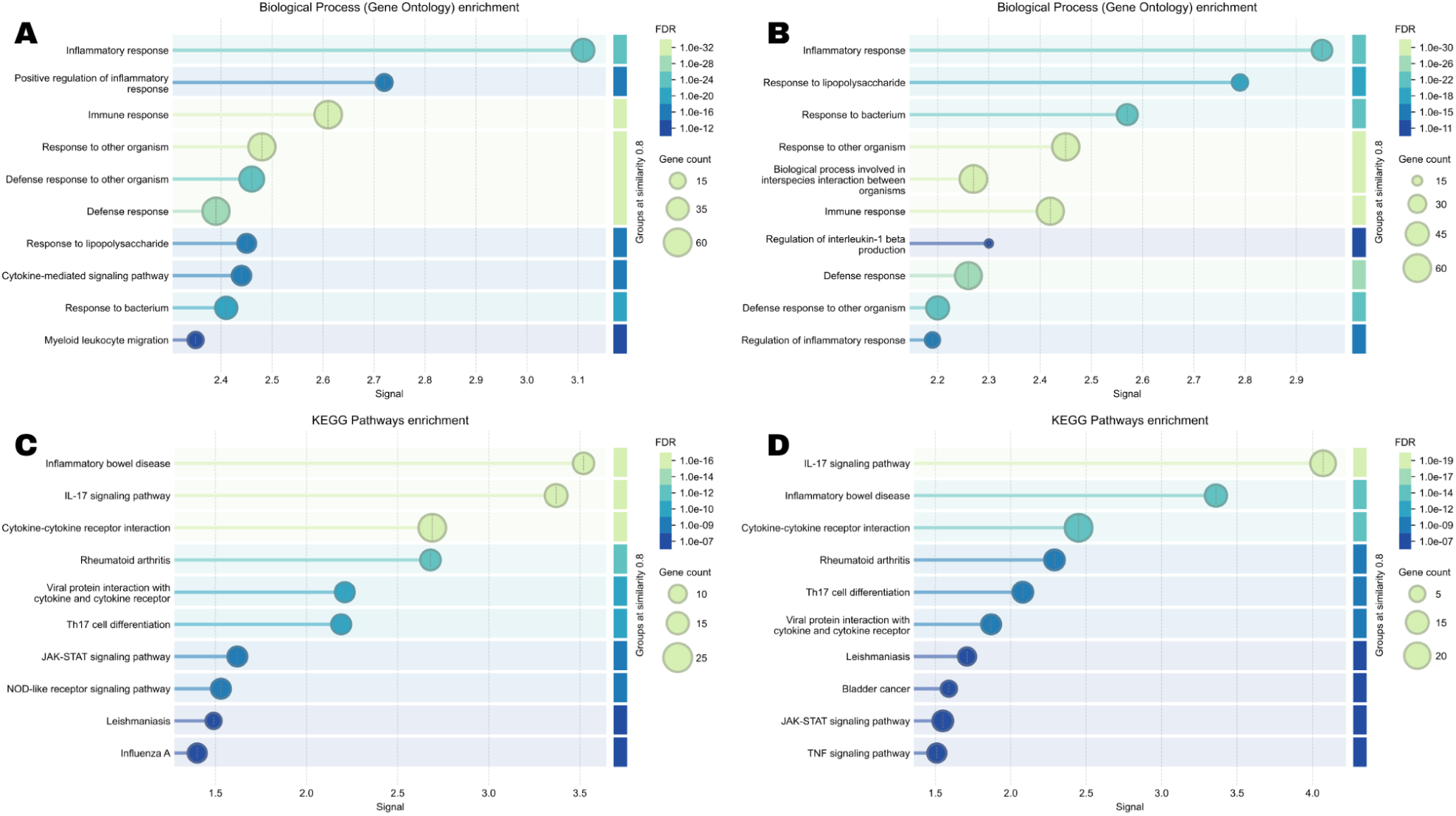
Comparative Functional Enrichment Analysis for Psoriatic Arthritis (PsA) Non-Lesional, PsA Lesional, and Ankylosing Spondylitis Normal Skin in Study GSE186063. The figure presents the results of the enrichment analysis of Gene Ontology (GO) Biological Process terms and KEGG pathways for the differentially expressed genes (DEGs). In all panels, the X-axis represents the enrichment signal strength, the size of the points corresponds to the number of genes associated with the term (Gene count), and the color indicates statistical significance (FDR). Panels (A) and (C) show the analysis for DEGs from the comparison between non-lesional and lesional tissue of PsA patients, with (A) showing GO terms and (C) showing KEGG pathways. Panels (B) and (D) show the analysis for DEGs from the comparison between normal skin of Ankylosing Spondylitis (AS) patients and PsA lesional tissue, with (B) showing GO terms and (D) showing KEGG pathways.

The functional enrichment analysis of the GSE117769 study elucidated both shared pathogenic axes and distinct molecular signatures between AS and PsA, as detailed in Figure 9. A central common mechanism was identified, with the “*IL-17 signaling pathway*” emerging as a cornerstone in both diseases when compared to healthy controls (Figure 9B and 9D). Despite the similarities, analysis indicates that ankylosing spondylitis (AS) and psoriatic arthritis (PsA) have distinct inflammatory mechanisms. In AS, inflammation appears to result from a broader immune dysregulation that promotes generalized “*leukocyte chemotaxis*” (Figure 9A). In contrast, in PsA, the inflammatory response is more targeted, focusing mainly on “*T-helper 17 cell differentiation*” (Figure 9C), considered a key element in the pathogenesis of the disease. The confirmation of molecular overlap between PsA and AS is evidenced by the direct analysis between the two diseases (Figure 9E), which notably showed no differential enrichment for major inflammatory pathways. Instead, enrichment was observed for catabolic processes, such as “*collagen metabolism*,” suggesting that the mechanisms distinguishing the two diseases may be more subtle and linked to tissue metabolism rather than acute inflammatory pathways.

**Figure 9.**
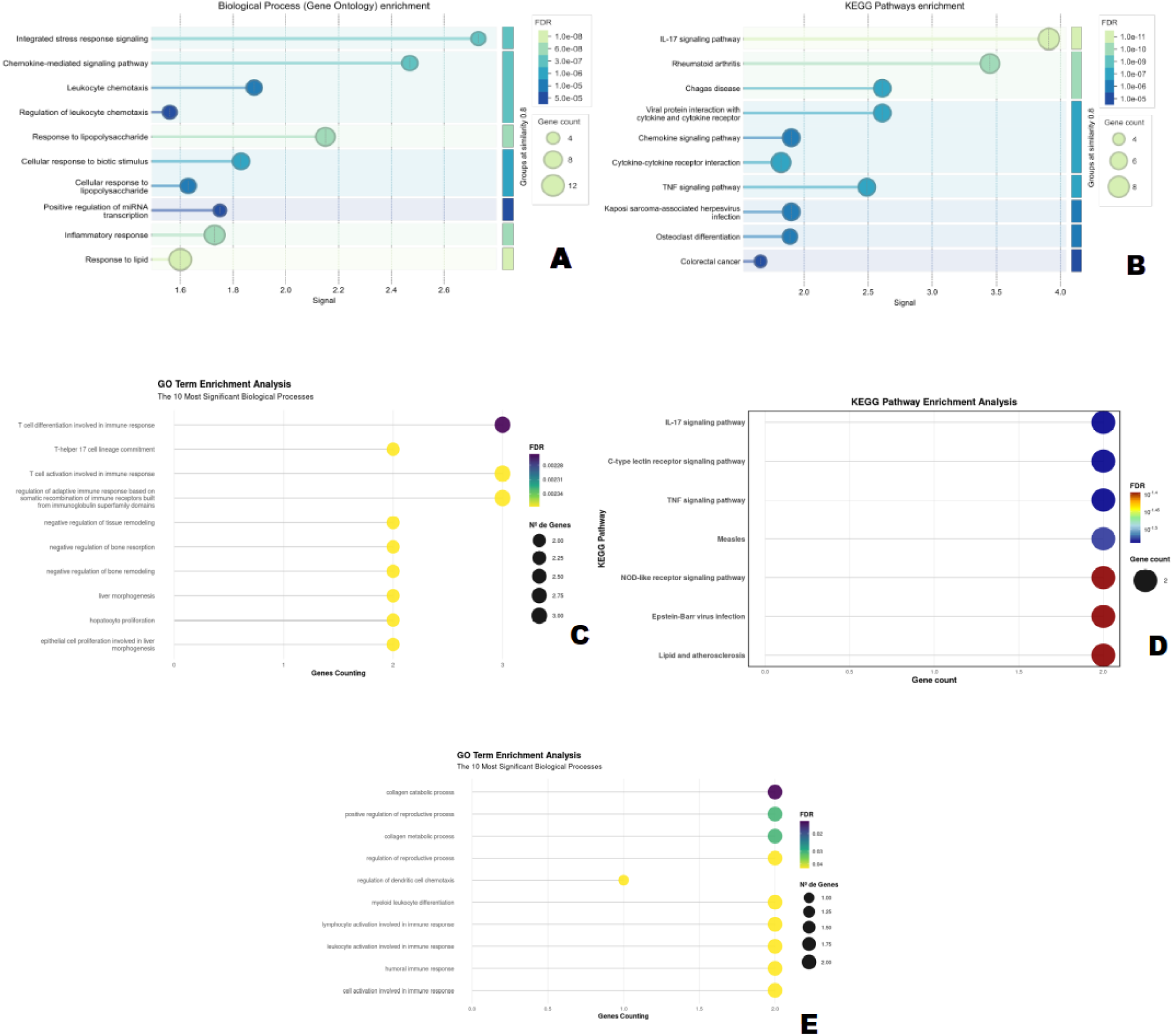
Comparative Functional Enrichment Analysis for Psoriatic Arthritis (PsA) and Ankylosing Spondylitis (AS) in Study GSE117769. The figure details the results of the functional enrichment analysis (Gene Ontology - GO and KEGG pathways) for three distinct comparisons. In panels A and B, the X-axis represents the enrichment ’Signal’, while in panels C, D, and E, it represents the gene count. In all graphs, the size and color of the points correspond to the gene count and statistical significance (FDR). Panels (A) and (B) present the analysis for differentially expressed genes (DEGs) between AS and healthy controls, highlighting ’leukocyte chemotaxis’ processes (GO) and the ’IL-17 signaling pathway’ (KEGG). Panels (C) and (D) show the analysis for DEGs between PsA and healthy controls, with strong enrichment for ’T-helper 17 cell differentiation’ (GO) and, consistently, the ’IL-17 signaling pathway’ (KEGG). Panel (E) illustrates the GO analysis for the direct comparison between PsA and AS, revealing a more subtle enrichment for immune pathways and highlighting metabolic processes.

Functional analysis of skin samples from PsA patients, AS patients, and healthy controls, utilizing datasets GSE205748 and GSE186063, yielded highly consistent results in comparisons between non-lesional vs. lesional skin, as well as lesional skin vs. healthy controls. The most frequently recurring Gene Ontology (GO) enrichment terms included “*inflammatory response*,” “*immune response*,” “*response to lipopolysaccharide,*” and “*response to bacterium*,” collectively demonstrating a robust inflammatory pattern within the lesions. Equally consistent, KEGG pathway analyses indicated the activation of the IL-17 signaling pathway, cytokine-cytokine receptor interaction, and associations with inflammatory diseases such as rheumatoid arthritis and inflammatory bowel disease. These findings were reproducible across different cohorts, highlighting a convergent functional signature characteristic of lesional PsA skin. Notably, the IL-17 pathway is solidified as a central axis in the pathophysiology of spondyloarthropathies.

### Gene Regulatory Network (GRN) Construction and Master Regulator (MR) Identification

The gene regulatory network was constructed from a matrix containing transcription factors (TFs), their target genes, associated regulatory motifs and frequency of occurrence (Supplementary Table 3). In the visualization, genes are represented in red and TFs in blue, allowing the functional classes of the nodes to be distinguished. The network topology shows a highly connected core, suggesting regions of coordinated regulation, while peripheral connections indicate possible independent regulatory modules, forming hubs with specific and potentially autonomous functions.

Topological analysis of the regulatory network unveiled nodes of high centrality, which can be categorized into two primary functional classes: connectivity hubs and master regulators. To identify the central regulatory elements within this biological network, a topology analysis was performed, and the connectivity degree of each node was calculated. Figure 11A presents the top 10 hub nodes, ranked by their total degree. After identifying the central hub nodes, a master regulator analysis was conducted to determine which transcription factors (TFs) exert the most significant regulatory influence over the network (Figure 11B). Notably, the gene PPARG emerged as the most prominent hub, exhibiting the highest degree of connectivity (total degree ≈ 10.0), suggesting a central and pleiotropic role in the investigated process. Furthermore, the presence of other significant hubs, such as STAT1, FOS, and STAT3 — well-known for their involvement in crucial cellular signaling pathways — reinforces the biological relevance of this network.

**Figure 10.**
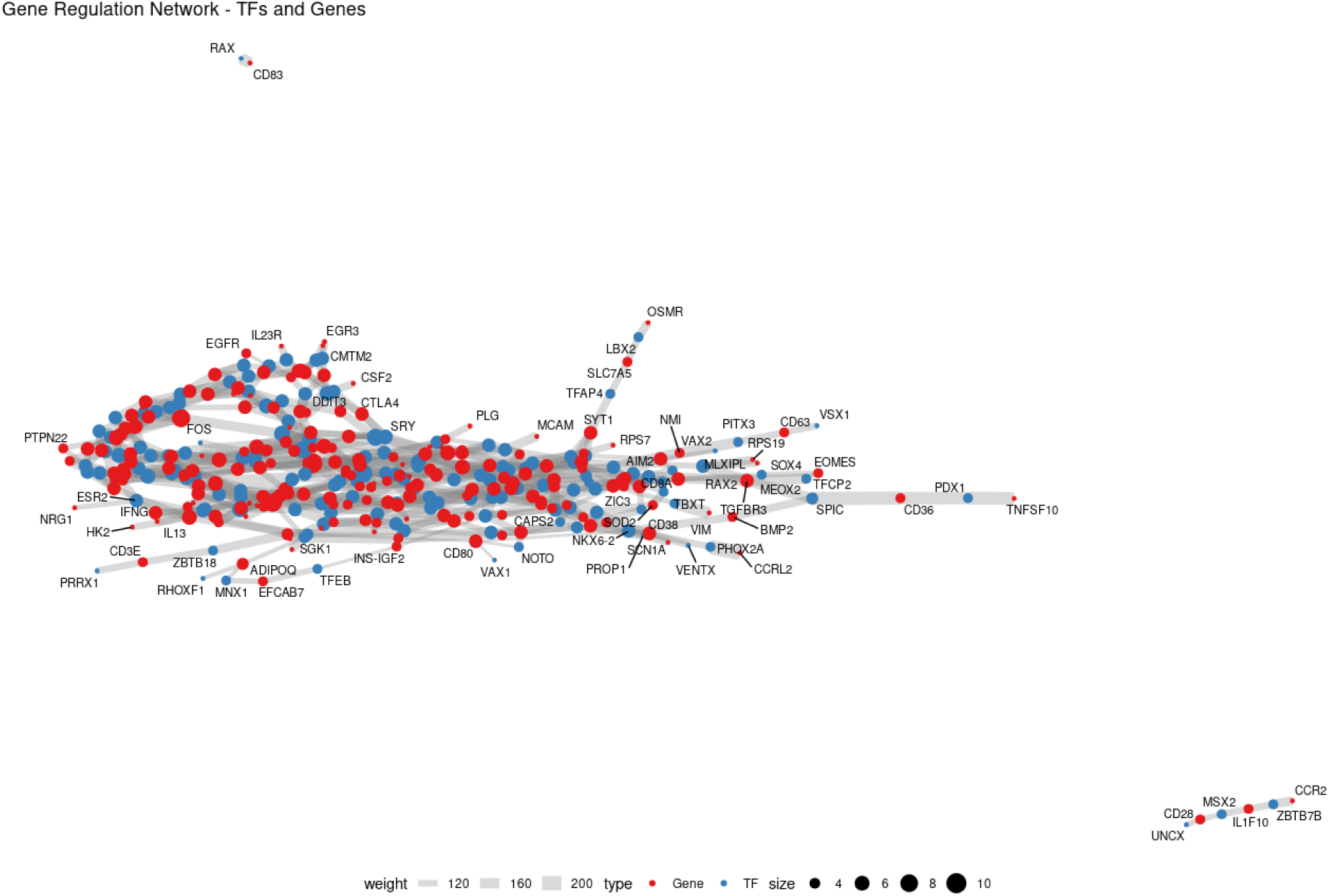
Visualization of the Gene Regulatory Network. The network illustrates the predicted regulatory interactions between Transcription Factors (TFs) and their target genes. Nodes in blue indicate TFs, and nodes in red represent genes (targets). The size of TF nodes is proportional to their connectivity degree in the network (centrality), highlighting the most influential TFs. Edges (lines) symbolize regulatory interactions, and their thickness (weight) is proportional to the strength or confidence of that interaction. The network topology reveals a densely connected main component and some disconnected gene clusters, suggesting the existence of distinct co-regulated modules.

**Figure 11.**
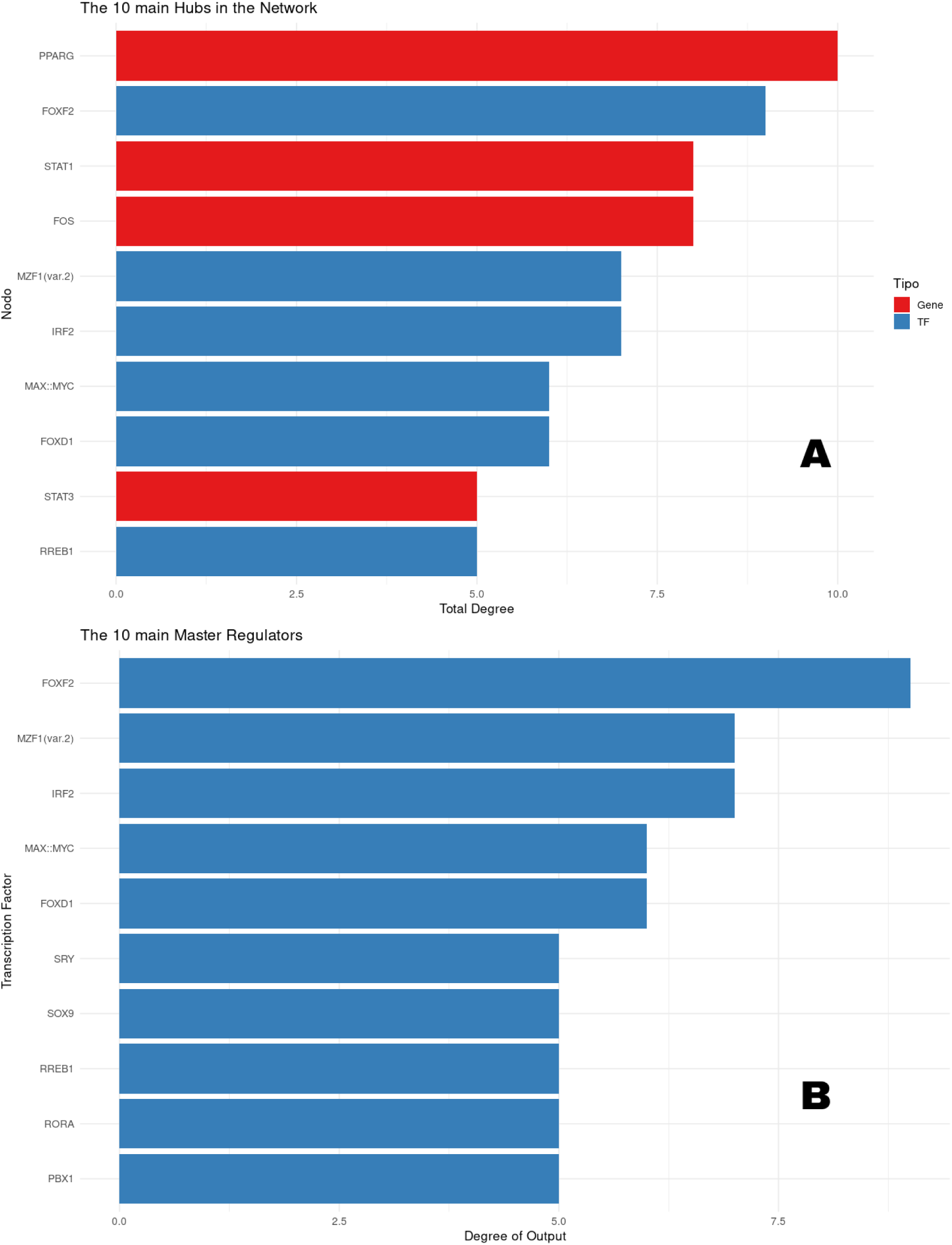
Identification and Classification of the Top 10 Hub Nodes in the Regulatory Network (A) and the rank of the Top 10 Transcription Factors (TFs) Identified as Master Regulators (B). (A) The 10 nodes (’hubs’) with the highest connectivity degree (’Total degree’) identified in the network analysis. The X-axis represents the total degree, which quantifies the number of interactions each node has within the network. The nodes (Y-axis) are categorized as ’Gene’ (red bars) and Transcription Factors (’TF’, blue bars). The nodes are ordered in decreasing fashion based on their total degree, highlighting the most influential elements of the network. (B) The 10 TFs with the highest master regulation potential, based on their “Degree of Output.” The X-axis quantifies the number of distinct target genes regulated by each TF (regulon size), serving as a measure of their downstream regulatory influence. The TFs (Y-axis) are ordered in decreasing fashion, highlighting those with the most significant capacity to control gene expression modules in the network.

Transcription factors (TFs) such as FOXF2 (out-degree ≈ 9.0), MZF1 (variant 2), and IRF2, with an out-degree of 7, indicate their fundamental role as direct regulators of gene expression. The MAX::MYC complex and FOXD1, each with an out-degree of 6, were also identified as significant master regulators, each controlling a substantial regulon. The identification of these TFs as master regulators implies they are primary drivers of the observed phenotypic state and represent high-priority candidates for experimental validation to confirm their causal role in network regulation.

## DISCUSSION

The present analysis investigated the molecular signatures and biological pathways associated with Psoriatic Arthritis (PsA) and Ankylosing Spondylitis (AS), integrating gene expression data from four public transcriptomic datasets. The analysis focused on evaluating the expression of 433 genes identified as potentially biologically relevant based on a text mining approach performed here. The results show that there are inflammatory mechanisms common to both diseases as well as molecular peculiarities that may reflect their distinct clinical manifestations.

The main molecular characteristic of PsA lies in a profound and consistent alteration of the cutaneous transcriptome (Veale and Fearon, 2018). Based on the comparative analysis of different skin specimens (non-lesional from PsA patients and normal skin from AS patients) with cutaneous lesions from PsA patients, it is possible to highlight the main similarities and differences in transcriptomic profiles. In all comparisons, the genes S100A8, S100A9, CXCL8, S100A12, PI3, IL17F, IL17A, CXCL13, IL21, and IL22 are consistently overexpressed in lesions, indicating a robust inflammatory pattern characteristic of lesional PsA skin. These genes are linked to neutrophil activation, Th1/Th17 cytokine production, and cell recruitment (Lowes et al., 2008; Chiricozzi et al., 2011). This reinforces the central role of the IL-23/IL-17 pathway in psoriatic inflammation (Baliwag et al., 2015). Genes like CCND1, GATA3, IL37, GPT, RORC, PPARGC1A, CYP1A1, KANK4, EGF, CSMD1, and ALB are elevated in non-lesional skin samples from PsA patients or normal skin from AS patients. This expression suggests a role in tissue maintenance and a less inflammatory cutaneous metabolism (Swindell et al., 2017). Additionally, the PPARG gene is also increased in non-lesional psoriatic arthritis (PsA) samples, and the FRZB and EFCA13 genes are increased in normal skin from AS patients.

The most prominent finding in the comparison of lesional PsA skin samples versus healthy controls is the massive overexpression of the S100A8 and S100A9 genes, as seen in the mean expression graphs (Figure 5C). These genes encode the calprotectin complex, a known marker of neutrophil activity and innate inflammation (Foell et al., 2009). Their dominance suggests that neutrophil infiltration and the calprotectin pathway are central events in psoriatic lesions. The overexpression of genes such as PI3 (Elafin) serves as an additional indicator of an intense innate immune response. Elafin, a protease inhibitor expressed by keratinocytes, has been consistently associated with inflammation and tissue repair in psoriasis (Inaba et al., 2003).

On the other hand, significant underexpression was observed in the genes IGF2-AS, IL4, CRB1, ALB, DKK1, KANK4, and CYP1A1 (Figure 5D). The reduction of IL-4, a cytokine classically associated with the Th2 immune response, corroborates the predominantly Th1/Th17 nature of psoriatic inflammation. The underexpression of ALB (albumin) in lesional skin may reflect changes in the skin barrier and systemic inflammatory response, since albumin is a negative acute phase protein: during inflammatory processes, the liver reduces its production to prioritize pro-inflammatory proteins (Garcia-Martínez et al., 2014; Kayser; Meyer; Morris, 2015). CYP1A1, a cytochrome P450 enzyme regulated by the aryl hydrocarbon receptor (AhR), is overexpressed in non-lesional skin, acting in metabolic defense and oxidative homeostasis (Sovenyák et al., 2019), but is underexpressed in lesional skin, possibly due to inhibition of the AhR pathway by proinflammatory cytokines such as TNF-α and IL-17, compromising detoxification and oxidative control (Vonta et al., 2019; Di Meglio; Villanova; Nestle, 2014).

In the non-lesional skin of individuals with PsA, only the SPP1 (osteopontin) gene was identified as differentially expressed compared to controls. Although this is a single marker, this finding is biologically relevant, since osteopontin plays central roles in modulating the immune response, activating T lymphocytes, and differentiating osteoclasts, in addition to being associated with chronic inflammatory processes (Chimenti et al., 2013; Ritchlin et al., 2017; Xu et al., 2021; Veale & Fearon, 2018). The presence of SPP1 in all comparisons performed (non-lesional skin vs. control, non-lesional vs. lesional, and lesional vs. control) suggests that its expression may reflect a continuous mechanism of inflammatory activation, regardless of the clinical stage of the lesion (Johnsson et al., 2023; Zhang et al., 2021). In this sense, osteopontin may act as an early sign of subclinical inflammation in PsA, anticipating the disruption of tissue homeostasis and contributing to the transition between asymptomatic states and clinical manifestations (Taheri et al., 2023; Chandran & Ritchlin, 2017).

From the data obtained from the GEO repository (GSE221786) (Zhao et al., 2023), which evaluated peripheral blood mononuclear cells (PBMCs) from ankylosing spondylitis (AS) patients after 4 hours of CytoStim stimulation (enriched with IL-17), only one differentially expressed gene (DEG) was identified (DYSF, overexpressed). The scarcity of DEGs may be related to restrictive analytical criteria (p.adj < 0.05 and |log2FC| > 1), the nature of the samples (*in vitro* stimulated PBMCs), or high biological variability among patients. Thus, in this cohort, the transcriptomic signature of AS in peripheral blood is discrete compared to our initial gene list, suggesting that pathological processes predominate in tissue compartments (such as entheses and joints), with systemic impact difficult to detect at the gene expression level (Baeten et al., 2015; Gracey et al., 2020). Alternatively, the molecular response to stimulation may be heterogeneous and insufficient to generate a robust differential profile.

A notable overlap is observed when comparing the DEGs obtained in the analyses of patients with PsA versus healthy individuals (Figure 4A) and AS versus healthy individuals (Figure 4B). Genes such as TNFAIP3 (a regulator of the TNF-alpha pathway) and CD69 are underexpressed in both pathologies. The former encodes the A20 protein, which acts as a negative inhibitor of the TNF-α/NF-κB signaling pathway, essential for controlling inflammation. The reduction in its expression has been associated with the persistence of inflammatory activity in autoimmune diseases, including spondyloarthritis (Catalán et al., 2015; Ma & Malone, 2020). CD69, in addition to being an early marker of lymphocyte activation, participates in the retention of T cells in the tissue by antagonizing the S1P1 receptor. Its underexpression observed in the peripheral blood of patients may indicate that a larger number of lymphocytes have already migrated to inflammatory sites, reflecting a process of cell activation and recruitment directed to target tissues. Thus, rather than suggesting an absence of inflammation, the decrease in CD69 in the blood can be interpreted as an indirect marker of lymphocyte redistribution to sites of active inflammation (Castro-Gómez et al., 2020; Czarnecki et al., 2021). Thus, the concomitant decrease in TNFAIP3 and CD69 may reflect distinct but convergent mechanisms that favor the maintenance of a chronic inflammatory state in PsA and AS, highlighting possible targets for immunomodulatory therapies.

The direct comparison between AS and PsA shows that, when the inflammatory signature common to both diseases is removed, the remaining transcriptomic differences are discrete. This suggests a basal similarity in systemic gene expression profiles, which corroborates the hypothesis of a partially shared pathogenic origin, especially in the Th17/IL-23 axis (Raychaudhuri et al., 2017; Veale & Fearon, 2018). Despite the global similarity, some genes on our list showed exclusive expression patterns. In AS, overexpression of the genes CCR6, CCR7, RORC, MYC, and CCL2 was observed, suggesting a more prominent activation of the Th17 axis and lymphocyte recruitment pathways (Lin et al., 2020; Menon et al., 2021). On the other hand, in PsA, the genes CLEC4D, OLR1, and PRTN3 were differentially overexpressed, possibly reflecting greater activation of myeloid cells and processes related to cutaneous inflammation and oxidative stress (Gudjonsson et al., 2019; Zhang et al., 2021). These findings reinforce that, although AS and PsA share a common immunological basis, discrete but biologically relevant differences may reflect the tissue, immunological, and clinical specificity of each condition (Ritchlin et al., 2017; McGonagle et al., 2020).

The construction of protein-protein interaction (PPI) networks from differentially expressed genes (DEGs) allows for the identification of molecular hubs and functional modules central to the pathophysiology of these diseases. We observed different profiles of connectivity and network density, which likely reflect varying degrees of inflammatory and immunological activation between tissues and clinical conditions (Barabási et al., 2011). The comparison between non-lesional and lesional skin in PsA (GSE186063) revealed a highly interconnected PPI network, with genes such as STAT1, CCL2, CD8A, GBP5, MX1, PPARG, and GBP1 positioned as functional hubs, involved in interferon-dependent responses, leukocyte chemotaxis, and immune response regulation (Appel et al., 2021). The similarity with the network from the comparison between normal AS skin and lesional PsA suggests a shared inflammatory core between spondyloarthritis, especially in components such as STAT1, GBP1, and CD8A. Additionally, the comparison between non-lesional and lesional skin in PsA (GSE205748) reinforced the presence of classic inflammatory hubs such as FOS, STAT1, CCL2, RORC, FOXP3, IL1A, CCL5, GBP5, CD8A, and PPARG, highlighting their participation in the modulation of active cutaneous inflammation (Richmond et al., 2020). The analysis of samples from PsA patients with lesions versus healthy controls further revealed the genes SPP1, HIF1A, CTLA4, STAT3, FOS, and FCGR3A as central, suggesting involvement of immunoregulatory and cell activation mechanisms (Müller et al., 2019), particularly in the context of active lesions (Smith et al., 2021).

Interestingly, the expression analyses of PBMC samples showed marked differences in network density and connectivity between the diseases. In comparison to healthy controls, the DEGs from AS patients formed a denser network, with relevant inflammatory hubs such as CXCL8, FOS, IFNG, MYC, and RORC, all associated with T-cell activation, inflammatory response, and transcriptional regulation (Ranganathan et al., 2021; Zhang et al., 2019; Chen et al., 2015). In contrast, in a similar comparison, the network formed by DEGs from PsA patients was sparse, with only a few connected genes, such as CD69, OLR1, TNFAIP3, and CLEC4D, also suggesting a more discrete or localized peripheral signature (Castro-Gómez et al., 2020; Catalán et al., 2015; Czarnecki et al., 2021). In a direct comparison between PsA and AS, the PPI network was restricted to PRTN3, MMP9, and RETN. The coexpression of these genes constitutes a signature of pathological neutrophilia, characterized by the activation and release of proteolytic enzymes and inflammatory mediators with high destructive potential for tissues (Zhang et al., 2023; Taheri et al., 2023; Chimenti et al., 2013; Zhang et al., 2019; Veale & Fearon, 2018). This profile reinforces the central role of innate immunity, especially neutrophil activation, as a shared mechanism in systemic inflammation and chronic tissue damage in both diseases (Baeten et al., 2015; Ritchlin et al., 2017; Colbert et al., 2010; Zhang et al., 2021).

Integrated, the four analyzed datasets reveal that PPARG showed increased expression specifically in PsA patients without skin lesions, suggesting a possible regulatory or protective function in the non-inflammatory cutaneous microenvironment (Müller et al., 2019; Veale & Fearon, 2018). STAT1 and FOS were identified as recurrent central axes in the networks of both diseases, reinforcing their role as regulators of chronic inflammation. Additionally, the RORC gene, widely recognized for encoding the RORγt transcription factor, was consistently overexpressed in both AS and non-lesional PsA, indicating its participation in Th17 cell differentiation. This finding is consistent with studies associating RORC with the activation of the IL-23/IL-17 axis, synovial inflammation, and epigenetic modulation via H3K27me3 (Chen et al., 2015; Bowness, 2015; Ivanov et al., 2006).

Functional enrichment analysis revealed that PsA and AS share a spectrum of inflammatory processes, but the nature of the differences depends heavily on the type of sample analyzed. In PsA lesion tissue samples, compared with non-lesion tissue from the same disease or from AS, the most prominent GO biological processes included “*inflammatory response*,” “*immune response*,” “*lipopolysaccharide response*” and *“bacterial response”* (Veale & Fearon, 2018; Taheri et al., 2023). The most enriched KEGG pathways were *“IL-17 signaling pathway,” “inflammatory bowel disease,” “rheumatoid arthritis,” “Th17 cell differentiation,”* and “*JAK-STAT signaling pathway*” (Zhang et al., 2019; Chandran & Ritchlin, 2017; Grotzinger et al., 2021). It is worth noting that some pathways, such as the “*TNF signaling pathway*”, only appeared when non-lesional AS tissue was compared with PsA lesion tissue, while the “*JAK-STAT signaling pathway*” was not observed when PsA lesion tissue was compared with healthy controls alone. These findings indicate that the functional signature of lesion tissue predominantly reflects local inflammatory activity and is strongly influenced by the microenvironment of the lesion (Veale & Fearon, 2018; Taheri et al., 2023). Thus, pathways that appear exclusively in comparisons involving lesion tissue likely represent responses associated with tissue damage and local repair, rather than intrinsic systemic differences between PsA and AS.

In peripheral blood samples, differences between PsA and AS better reflect systemic disease responses. In AS, specific biological processes included “*integrated stress response signaling*”, “*chemokine-mediated signaling pathway*”, “*leukocyte chemotaxis*”, “*inflammatory response*”, “*lipopolysaccharide response*” and “*lipid response*” while the most enriched pathways were “*viral protein interaction with cytokine receptor*”, “*Chagas disease*”, “*chemokine signaling pathway*”, “*herpesvirus infection associated with Kaposi’s sarcoma*” and “*osteoclast differentiation*” (Appel et al., 2021; Liu et al., 2020; Akdis et al., 2016; KEGG, 2024). In PsA, biological processes emphasized the immune response involving T cell activation and differentiation, as well as the regulation of the adaptive immune response, while the most prominent pathways included “*C-type lectin receptor signaling pathway*”, “*measles”,* “*Epstein-Barr virus infection*” and “*lipids and atherosclerosis*” (Chandran & Ritchlin, 2017; Grotzinger et al., 2021; Zhang et al., 2023).

The pathways shared between AS and PsA in peripheral blood mainly included IL-17 signaling and TNF signaling, confirming the presence of a common central inflammatory axis (Chandran & Ritchlin, 2017; Grotzinger et al., 2021). At the same time, associations with lipid metabolism were observed: in AS, differentially expressed genes showed enrichment for the biological process of “*response to lipids*” (GO), while in PsA there was enrichment of the “*lipid and atherosclerosis*” pathway (KEGG). These findings suggest that, in addition to cytokine-mediated inflammation, lipid metabolic dysfunction constitutes a shared molecular mechanism, with possible implications for increased cardiovascular risk in both diseases (Wang et al., 2022; Huang et al., 2023).

Direct comparison between AS and PsA in blood revealed biological processes such as *“collagen catabolic process”, “upregulation of reproductive process”* and *“collagen metabolic process”* (Zhang et al., 2019; Taheri et al., 2023). These processes suggest subtle systemic differences between the diseases.

The construction of the Gene Regulatory Network (GRN) allowed for the identification of central genes and transcription factors (TFs) in controlling the differential gene expression observed in PsA and AS. Our analysis integrated transcriptomic data with network inference models to identify hubs and master regulators (MRs), which play central roles in orchestrating the inflammatory and immunological responses characteristic of these diseases. The general network structure revealed a dense core of interactions, with genes such as PPARG, STAT1, FOS, and FOXF2 figuring among the main hubs. The PPARG gene, widely recognized for its anti-inflammatory function, appeared as the most prominent hub, suggesting an attempt to counter-regulate inflammation in affected tissues, particularly in lesional skin of PsA patients (Grotzinger et al., 2021). This finding reinforces the idea that there is an active adaptive immunometabolic response in this group. STAT1 and FOS, in turn, are classic mediators of signaling induced by inflammatory cytokines (such as IFN-γ and TNF-α), being directly associated with the activation of immunological pathways in both PsA and AS (Zhang et al., 2020). Their presence among the main hubs confirms their role in amplifying chronic inflammation. FOXF2, a less studied factor, showed high connectivity and may be involved in cell differentiation and tissue integrity maintenance, especially in the early or non-lesional phases of PsA, indicating a potential role in the transition between latent and manifest inflammation (Hannenhalli & Kaestner, 2009).

The analysis of master regulators revealed FOXF2 as the transcription factor with the highest out-degree, regulating a significant number of differentially expressed genes. This TF, along with FOXD1, both belonging to the FOX family, is involved in tissue homeostasis, epithelial differentiation, and structural organization of tissues, especially in inflammatory and repair contexts (HANNENHALLI & KAESTNER, 2009). In Psoriatic Arthritis (PsA), these factors emerge as relevant regulators in non-lesional tissues, suggesting a possible preventive or organizing role in epithelial architecture before the onset of inflammation (Hannenhalli & Kaestner, 2009). Similarly, in Ankylosing Spondylitis (AS), although the affected tissues are predominantly articular, members of the FOX family have also been associated with mesenchymal cell differentiation and aberrant osteogenesis, central processes in AS pathophysiology (WANG et al., 2018; XU et al., 2021). The MZF1 (var.2) factor simultaneously stood out as a master regulator and hub, having been previously implicated in myeloid differentiation, epigenetic remodeling, and hematopoietic control (YANG et al., 2017). In PsA, its relevance is corroborated by the intense infiltration of neutrophils and myeloid cells into cutaneous lesions, suggesting a role in maintaining a persistent inflammatory microenvironment. In AS, where innate immune cells (such as monocytes and neutrophils) are involved in enthesitis and sacroiliac joints, MZF1 may contribute to the modulation of the local immune response and favor the transition from inflammation to pathological bone remodeling (CHEN et al., 2019).

Another relevant factor was the MAX::MYC complex, whose regulatory activity is related to cell proliferation, metabolism, and adaptive immune responses (Dang, 2012). Its participation in the network suggests involvement with the expansion of activated T cells, a process widely described in AS and also present in PsA, especially in joint tissues. The action of this complex may contribute to the metabolic reprogramming of inflammatory cells, a phenomenon increasingly recognized in the pathogenesis of spondyloarthritis. IRF2, also identified as a master regulator and hub, is a key modulator of the type I and type II interferon response. Its activation is consistently associated with hyperactivity of the interferon pathway in AS, especially in patients with extra-articular manifestations and an exacerbated response to endogenous or environmental viral stimuli (Ranganathan et al., 2021). Its presence reinforces the role of exacerbated innate immunity in AS, aligning with the functional findings of enrichment in antiviral pathways. Finally, RORA is a crucial regulator of Th17 cell differentiation, being directly implicated in the IL-23/IL-17 pathway, one of the central immunological axes in both diseases, but particularly important in cutaneous and articular PsA (Ivanov et al., 2006). Its role as a master regulator suggests that transcriptional changes in RORA can significantly influence the intensity and type of Th17 response, impacting the severity and location of inflammation.

The GRN data reveal robust interconnectivity between classic inflammatory hubs (STAT1, FOS, PPARG) and new transcriptional regulators with possibly modulating functions (FOXF2, MZF1, MAX::MYC). This complex regulatory network reflects the overlap of common immunological mechanisms between PsA and AS — such as the activation of inflammatory cytokines and Th17 cells — but also suggests essential differences in the predominant regulatory cell type and the tissue context of inflammation. In summary, the mapping of the gene regulatory network shows that FOXF2, MZF1 (var.2), MAX::MYC, IRF2, and RORA are central transcription factors that, in addition to regulating the expression of multiple genes involved in inflammation, functionally differentiate the molecular signatures of PsA and AS. This reinforces the potential of these regulators as molecular biomarkers or specific therapeutic targets for each subtype of spondyloarthritis.

It is important to emphasize that the present study and its approach have significant limitations that impact the interpretation of transcriptomic results and their application in both diseases. The main restriction refers to the scarcity and heterogeneity of public RNA-Seq data available for these diseases. Only four datasets were analyzed, with incomplete clinical information, a lack of standardization regarding disease stage, therapies in use, comorbidities, and patient lifestyle, which compromises sample stratification and the external validity of the findings (Barendregt et al., 2020; Mease et al., 2021). The tissue diversity among the datasets also constitutes a critical source of bias, given that gene expression is highly dependent on the type of tissue analyzed (Rajapakse and Anderson, 2020). Furthermore, most of the data do not include longitudinal analyses, limiting the evaluation of the temporal dynamics of inflammatory processes (Liu et al., 2023). Such factors highlight the need for more robust, longitudinal studies with greater methodological standardization to deepen the understanding of the molecular signatures of these inflammatory autoimmune diseases.

## CONCLUSION

The analyzed gene expression data demonstrate the effective involvement of many of the previously selected 433 genes, corroborating several findings described in the literature. At the same time, our analysis revealed less-explored elements with potential as biomarkers. The integrated analysis of transcriptomic data, protein-protein interaction (PPI) networks, functional enrichment, and master regulator prediction provides a comprehensive view of the molecular mechanisms underpinning Psoriatic Arthritis and Ankylosing Spondylitis. Although both diseases share a common inflammatory axis, the observed differences in differentially expressed gene (DEG) profiles, hub genes identified in PPI networks, and functionally enriched pathways highlight immunopathological particularities that contribute to their distinct clinical manifestations. The identification of central regulatory genes like PPARG, STAT1, FOS, and STAT3 reinforces their role in modulating the immune and inflammatory response. At the same time, the implication of less-explored transcription factors, such as FOXF2, points to emerging regulatory pathways that may be involved in the differential pathogenesis of these spondyloarthropathies. These findings not only deepen our biological understanding of these diseases but also open new perspectives for translational research and the development of more specific and targeted therapeutic strategies.

## ACKNOWLEDGMENTS

The authors are indebted to the High-Performance Computing Center (NPAD/UFRN) and the Multidisciplinary Bioinformatics Environment (BioME/UFRN) for providing computing resources. The text’s language was reviewed using AI-assisted tools, including Grammarly (Premium, version 2.0) and Gemini 2.5. The authors express their gratitude to the developers of these tools.

## Supplementary Material

**Supplementary Table 1** - 433 genes in common with AS and PsA.

**Supplementary Table 2** - Gene expression profile conserved in non-lesion skin states (PsA and AS) compared to psoriatic lesions.

**Supplementary Table 3 -** Associations between Genes and Transcription Factors Identified by Gene Expression Analysis for Psoriatic Arthritis (PsA) and Ankylosing Spondylitis (AS).

**Supplementary Figure 1.**
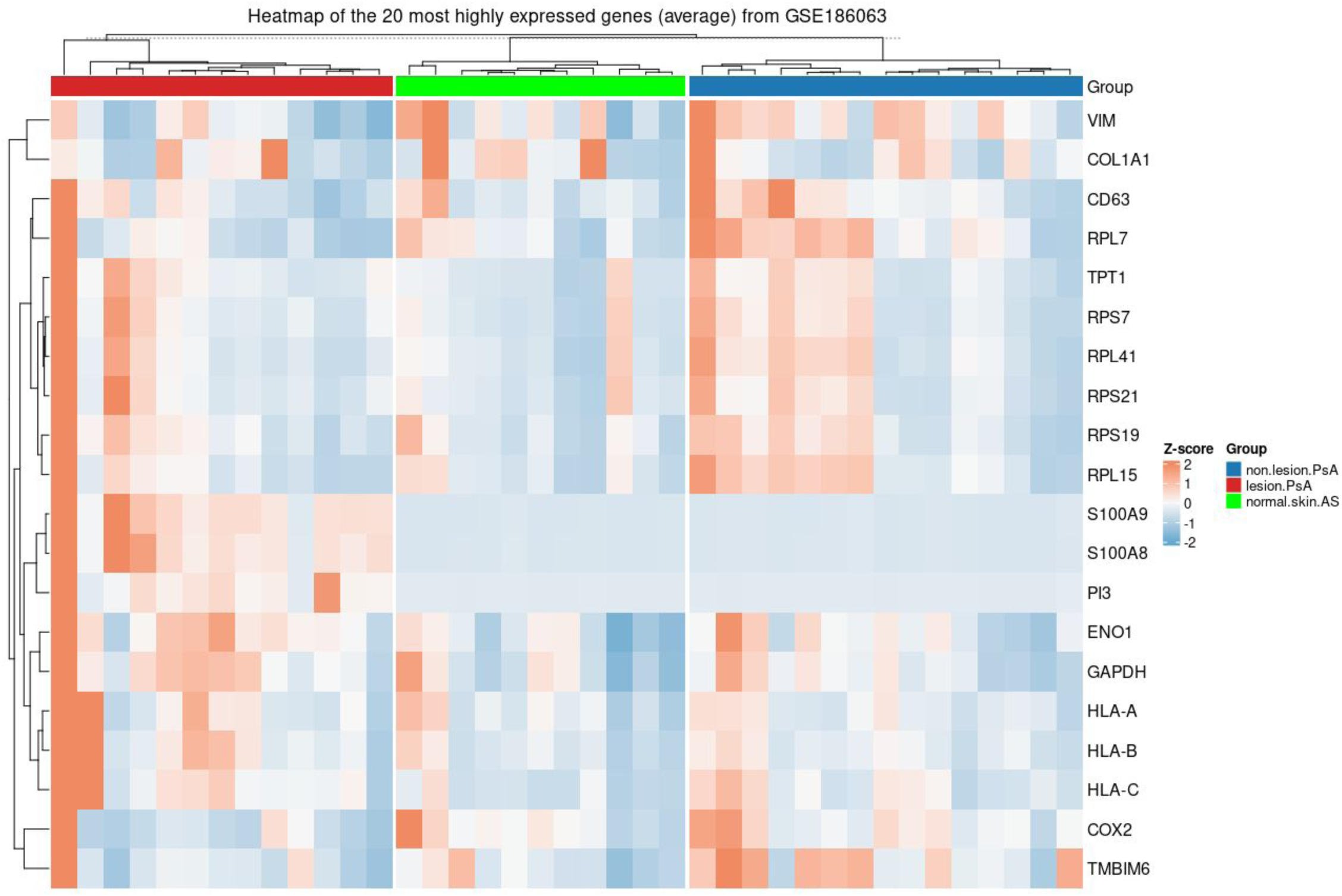
Heatmap of the most 20 highly expressed genes for our list (average) from GSE186063.

**Supplementary Figure 2.**
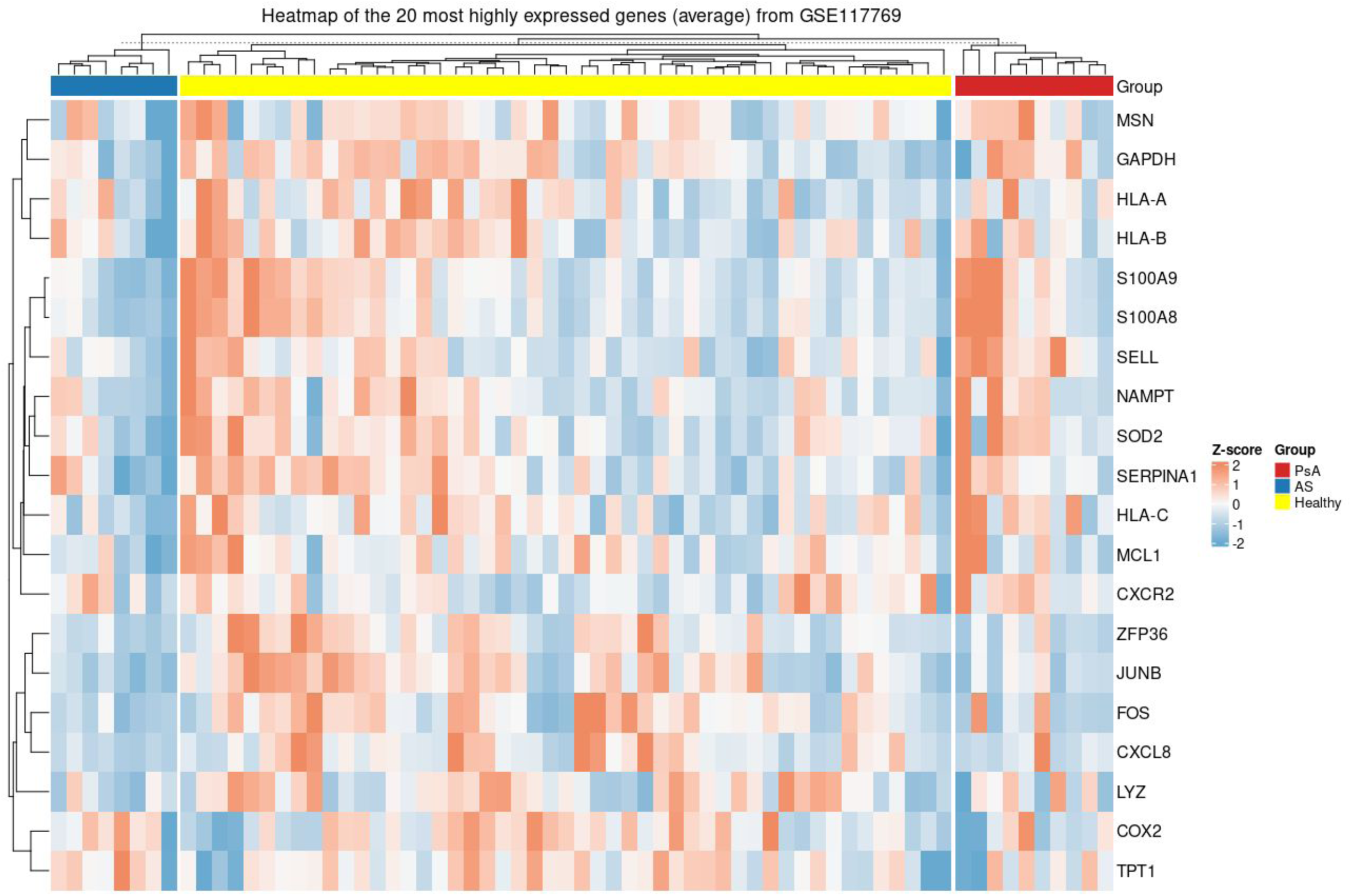
Heatmap of the most 20 highly expressed genes for our list (average) from GSE117769.

**Supplementary Figure 3.**
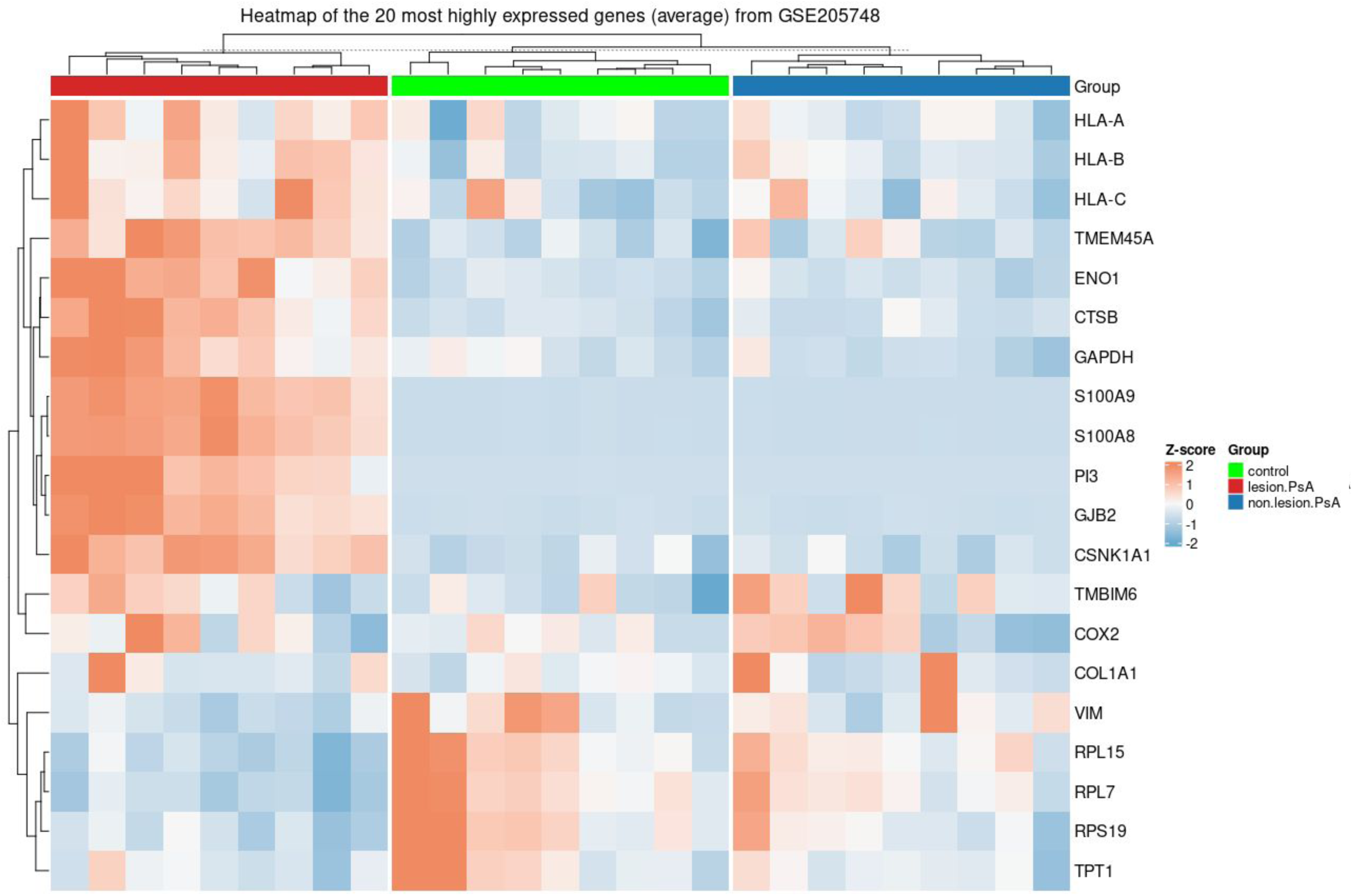
Heatmap of the most 20 highly expressed genes for our list (average) from GSE205748.

