## Supplementary Table 1 for "Beyond the Shared Inflammatory Axis: Differentiating Molecular Signatures in Psoriatic Arthritis and Ankylosing Spondylitis through Integrated Omics"

**Supplementary Table 01 - 433 genes in common with AS and PsA**

| Genes | gene accession | CHROMOSOME | Start position | End position | Strand | Description |
|---|---|---|---|---|---|---|
| ABCB11 | ENSG00000073734 | 2 | 168915498 | 169031324 | (-) | ATP binding cassette subfamily B member 11 |
| ABCD2 | ENSG00000173208 | 12 | 39550033 | 39619803 | (-) | ATP binding cassette subfamily D member 2 |
| ACE | ENSG00000159640 | 17 | 63477061 | 63498380 | (+) | angiotensin I converting enzyme |
| ACKR2 | ENSG00000144648 | 3 | 42804752 | 42887974 | (+) | atypical chemokine receptor 2 |
| ACP5 | ENSG00000102575 | 19 | 11574660 | 11579008 | (-) | acid phosphatase 5, tartrate resistant |
| ACP7 | ENSG00000183760 | 19 | 39083913 | 39111493 | (+) | acid phosphatase 7, tartrate resistant (putative) |
| ADAMTS9 | ENSG00000163638 | 3 | 64515654 | 64688000 | (-) | ADAM metallopeptidase with thrombospondin type 1 motif 9 |
| ADCY7 | ENSG00000121281 | 16 | 50246137 | 50318135 | (+) | adenylate cyclase 7 |

| | | | | | | |
|---|---|---|---|---|---|---|
| ADGRL2 | ENSG00000117114 | 1 | 81306147 | 81992436 | (+) | adhesion G protein-coupled receptor L2 |
| ADIPOQ | ENSG00000181092 | 3 | 186842704 | 186858463 | (+) | adiponectin, C1Q and collagen domain containing |
| AHR | ENSG00000106546 | 7 | 16916359 | 17346152 | (+) | aryl hydrocarbon receptor |
| AIM2 | ENSG00000163568 | 1 | 159062484 | 159147096 | (-) | absent in melanoma 2 |
| AKAP13 | ENSG00000170776 | 15 | 85380571 | 85749358 | (+) | A-kinase anchoring protein 13 |
| ALB | ENSG00000163631 | 4 | 73397114 | 73421482 | (+) | albumin |
| ANKRD55 | ENSG00000164512 | 5 | 56099680 | 56233330 | (-) | ankyrin repeat domain 55 |
| ANXA1 | ENSG00000135046 | 9 | 73151865 | 73170393 | (+) | annexin A1 |
| APOL1 | ENSG00000100342 | 22 | 36253071 | 36267530 | (+) | apolipoprotein L1 |
| APOL6 | ENSG00000221963 | 22 | 35648446 | 35668404 | (+) | apolipoprotein L6 |
| AQP1 | ENSG00000240583 | 7 | 30911853 | 30925517 | (+) | aquaporin 1 (Colton blood group) |

| | | | | | | |
|---|---|---|---|---|---|---|
| AREG | ENSG00000109321 | 4 | 74445136 | 74455005 | (+) | amphiregulin |
| ARHGEF3 | ENSG00000163947 | 3 | 56727418 | 57079329 | (-) | Rho guanine nucleotide exchange factor 3 |
| ATG16L1 | ENSG00000085978 | 2 | 233210051 | 233295674 | (+) | autophagy related 16 like 1 |
| ATG5 | ENSG00000057663 | 6 | 106045423 | 106325791 | (-) | autophagy related 5 |
| ATOX1 | ENSG00000177556 | 5 | 151742316 | 151772532 | (-) | antioxidant 1 copper chaperone |
| ATXN2L | ENSG00000168488 | 16 | 28822999 | 28837237 | (+) | ataxin 2 like |
| BEND2 | ENSG00000177324 | X | 18162931 | 18220886 | (-) | BEN domain containing 2 |
| BGLAP | ENSG00000242252 | 1 | 156242184 | 156243317 | (+) | bone gamma-carboxyglutamate protein |
| BMP2 | ENSG00000125845 | 20 | 6767686 | 6780246 | (+) | bone morphogenetic protein 2 |
| BMP7 | ENSG00000101144 | 20 | 57168753 | 57266641 | (-) | bone morphogenetic protein 7 |
| BNIP3L | ENSG00000104765 | 8 | 26383054 | 26505636 | (+) | BCL2 interacting protein 3 like |

| | | | | | | |
|---|---|---|---|---|---|---|
| BRAF | ENSG00000157764 | 7 | 140719327 | 140924929 | (-) | B-Raf proto-oncogene, serine/threonine kinase |
| BTG1 | ENSG00000133639 | 12 | 92140278 | 92145846 | (-) | BTG anti-proliferation factor 1 |
| C15ORF48 | ENSG00000166920 | 15 | 45430579 | 45448761 | (+) | chromosome 15 open reading frame 48 |
| C7ORF57 | ENSG00000164746 | 7 | 48035511 | 48061304 | (+) | chromosome 7 open reading frame 57 |
| CALD1 | ENSG00000122786 | 7 | 134744252 | 134970729 | (+) | caldesmon 1 |
| CAPS2 | ENSG00000180881 | 12 | 75275979 | 75390928 | (-) | calcyphosine 2 |
| CARD9 | ENSG00000187796 | 9 | 136363956 | 136373681 | (-) | caspase recruitment domain family member 9 |
| CASP1 | ENSG00000137752 | 11 | 105025397 | 105035250 | (-) | caspase 1 |
| CASP10 | ENSG00000003400 | 2 | 201182872 | 201229428 | (+) | caspase 10 |
| CAST | ENSG00000153113 | 5 | 95962001 | 96631085 | (+) | calpastatin |
| CCL2 | ENSG00000108691 | 17 | 34255274 | 34257208 | (+) | C-C motif chemokine ligand 2 |

| | | | | | | |
|---|---|---|---|---|---|---|
| CCL20 | ENSG00000115009 | 2 | 227805739 | 227817564 | (+) | C-C motif chemokine ligand 20 |
| CCL25 | ENSG00000131142 | 19 | 8052318 | 8062660 | (+) | C-C motif chemokine ligand 25 |
| CCL3 | ENSG00000277632 | 17 | 36088256 | 36090169 | (-) | C-C motif chemokine ligand 3 |
| CCL5 | ENSG00000271503 | 17 | 35871491 | 35880793 | (-) | C-C motif chemokine ligand 5 |
| CCND1 | ENSG00000110092 | 11 | 69641156 | 69654474 | (+) | cyclin D1 |
| CCND3 | ENSG00000112576 | 6 | 41934934 | 42050357 | (-) | cyclin D3 |
| CCR2 | ENSG00000121807 | 3 | 46353864 | 46360940 | (+) | C-C motif chemokine receptor 2 |
| CCR6 | ENSG00000112486 | 6 | 167111807 | 167139141 | (+) | C-C motif chemokine receptor 6 |
| CCR7 | ENSG00000126353 | 17 | 40551081 | 40565472 | (-) | C-C motif chemokine receptor 7 |
| CCRL2 | ENSG00000121797 | 3 | 46407166 | 46412997 | (+) | C-C motif chemokine receptor like 2 |
| CD14 | ENSG00000170458 | 5 | 140631728 | 140633700 | (-) | CD14 molecule |

| | | | | | | |
|---|---|---|---|---|---|---|
| CD160 | ENSG00000117281 | 1 | 145719471 | 145739288 | (+) | CD160 molecule |
| CD19 | ENSG00000177455 | 16 | 28931965 | 28939342 | (+) | CD19 molecule |
| CD274 | ENSG00000120217 | 9 | 5450503 | 5470566 | (+) | CD274 molecule |
| CD28 | ENSG00000178562 | 2 | 203706517 | 203739756 | (+) | CD28 molecule |
| CD36 | ENSG00000135218 | 7 | 80369575 | 80679277 | (+) | CD36 molecule (CD36 blood group) |
| CD38 | ENSG00000004468 | 4 | 15778275 | 15853232 | (+) | CD38 molecule |
| CD3E | ENSG00000198851 | 11 | 118304730 | 118316175 | (+) | CD3 epsilon subunit of T-cell receptor complex |
| CD4 | ENSG00000010610 | 12 | 6786858 | 6820799 | (+) | CD4 molecule |
| CD40 | ENSG00000101017 | 20 | 46118271 | 46129863 | (+) | CD40 molecule |
| CD40LG | ENSG00000102245 | X | 136648158 | 136660390 | (+) | CD40 ligand |
| CD52 | ENSG00000169442 | 1 | 26317958 | 26320523 | (+) | CD52 molecule |

| | | | | | | |
|---|---|---|---|---|---|---|
| CD63 | ENSG00000135404 | 12 | 55725323 | 55729707 | (-) | CD63 molecule |
| CD68 | ENSG00000129226 | 17 | 7579491 | 7582111 | (+) | CD68 molecule |
| CD69 | ENSG00000110848 | 12 | 9752486 | 9760901 | (-) | CD69 molecule |
| CD80 | ENSG00000121594 | 3 | 119524293 | 119559614 | (-) | CD80 molecule |
| CD83 | ENSG00000112149 | 6 | 14117256 | 14140682 | (+) | CD83 molecule |
| CD86 | ENSG00000114013 | 3 | 122055362 | 122121139 | (+) | CD86 molecule |
| CD8A | ENSG00000153563 | 2 | 86784610 | 86808396 | (-) | CD8 subunit alpha |
| CD8B | ENSG00000172116 | 2 | 86815339 | 86861924 | (-) | CD8 subunit beta |
| CDC42BPB | ENSG00000198752 | 14 | 102932380 | 103057549 | (-) | CDC42 binding protein kinase beta |
| CEBPA | ENSG00000245848 | 19 | 33299934 | 33302534 | (-) | CCAAT enhancer binding protein alpha |
| CEBPG | ENSG00000153879 | 19 | 33373685 | 33382686 | (+) | CCAAT enhancer binding protein gamma |

| | | | | | | |
|---|---|---|---|---|---|---|
| CENPK | ENSG00000123219 | 5 | 65517766 | 65563168 | (-) | centromere protein K |
| CLEC2B | ENSG00000110852 | 12 | 9852369 | 9869386 | (-) | C-type lectin domain family 2 member B |
| CLEC4D | ENSG00000166527 | 12 | 8509475 | 8522366 | (+) | C-type lectin domain family 4 member D |
| CLIC3 | ENSG00000169583 | 9 | 136994608 | 136996568 | (-) | chloride intracellular channel 3 |
| CMAHP | ENSG00000168405 | 6 | 25061626 | 25452263 | (-) | cytidine monophospho-N-acetylneuraminic acid hydroxylase, pseudogene;uncharacterized LOC101928663 |
| CMTM2 | ENSG00000140932 | 16 | 66579448 | 66588275 | (+) | CKLF like MARVEL transmembrane domain containing 2 |
| COL1A1 | ENSG00000108821 | 17 | 50184101 | 50201632 | (-) | collagen type I alpha 1 chain |
| CRB1 | ENSG00000134376 | 1 | 197268204 | 197478455 | (+) | crumbs cell polarity complex component 1 |
| CREM | ENSG00000095794 | 10 | 35126791 | 35212958 | (+) | cAMP responsive element modulator |
| CRP | ENSG00000132693 | 1 | 159712289 | 159714589 | (-) | C-reactive protein |

| | | | | | | |
|---|---|---|---|---|---|---|
| CSF2 | ENSG00000164400 | 5 | 132073789 | 132076170 | (+) | colony stimulating factor 2 |
| CSMD1 | ENSG00000183117 | 8 | 2935353 | 4994972 | (-) | CUB and Sushi multiple domains 1 |
| CSN3 | ENSG00000171209 | 4 | 70238382 | 70251474 | (+) | casein kappa |
| CSNK1A1 | ENSG00000113712 | 5 | 149492982 | 149551471 | (-) | casein kinase 1 alpha 1 |
| CTLA4 | ENSG00000163599 | 2 | 203853888 | 203873965 | (+) | cytotoxic T-lymphocyte associated protein 4 |
| CTSB | ENSG00000164733 | 8 | 11842524 | 11869533 | (-) | cathepsin B |
| CTSK | ENSG00000143387 | 1 | 150794880 | 150809577 | (-) | cathepsin K |
| CUX1 | ENSG00000257923 | 7 | 101815904 | 102283958 | (+) | cut like homeobox 1 |
| CX3CR1 | ENSG00000168329 | 3 | 39263495 | 39281735 | (-) | C-X3-C motif chemokine receptor 1 |
| CXCL10 | ENSG00000169245 | 4 | 76021118 | 76023497 | (-) | C-X-C motif chemokine ligand 10 |
| CXCL13 | ENSG00000156234 | 4 | 77511753 | 77611834 | (+) | C-X-C motif chemokine ligand 13 |

| | | | | | | |
|---|---|---|---|---|---|---|
| CXCL16 | ENSG00000161921 | 17 | 4733533 | 4739928 | (-) | C-X-C motif chemokine ligand 16 |
| CXCL2 | ENSG00000081041 | 4 | 74097040 | 74099196 | (-) | C-X-C motif chemokine ligand 2 |
| CXCL8 | ENSG00000169429 | 4 | 73740519 | 73747379 | (+) | C-X-C motif chemokine ligand 8 |
| CXCR2 | ENSG00000180871 | 2 | 218125289 | 218137251 | (+) | C-X-C motif chemokine receptor 2 |
| CYCS | ENSG00000172115 | 7 | 25118656 | 25125260 | (-) | cytochrome c, somatic |
| CYP1A1 | ENSG00000140465 | 15 | 74719542 | 74725536 | (-) | cytochrome P450 family 1 subfamily A member 1 |
| CYP4F22 | ENSG00000171954 | 19 | 15508525 | 15552317 | (+) | cytochrome P450 family 4 subfamily F member 22 |
| DDIT3 | ENSG00000175197 | 12 | 57516588 | 57521737 | (-) | DNA damage inducible transcript 3 |
| DDX60 | ENSG00000137628 | 4 | 168216291 | 168318883 | (-) | DExD/H-box helicase 60 |
| DKK1 | ENSG00000107984 | 10 | 52314281 | 52318042 | (+) | dickkopf WNT signaling pathway inhibitor 1 |

| | | | | | | |
|---|---|---|---|---|---|---|
| DLAT | ENSG00000150768 | 11 | 112025033 | 112064404 | (+) | dihydrolipoamide S-acetyltransferase |
| DNAJA2 | ENSG00000069345 | 16 | 46955362 | 46973674 | (-) | DnaJ heat shock protein family (Hsp40) member A2 |
| DNAJB6 | ENSG00000105993 | 7 | 157335381 | 157417439 | (+) | DnaJ heat shock protein family (Hsp40) member B6 |
| DUSP4 | ENSG00000120875 | 8 | 29333064 | 29350684 | (-) | dual specificity phosphatase 4 |
| DYSF | ENSG00000135636 | 2 | 71453561 | 71686763 | (+) | dysferlin |
| EDNRA | ENSG00000151617 | 4 | 147480917 | 147544954 | (+) | endothelin receptor type A |
| EFCAB13 | ENSG00000178852 | 17 | 47323290 | 47441312 | (+) | EF-hand calcium binding domain 13 |
| EFCAB7 | ENSG00000203965 | 1 | 63523372 | 63572693 | (+) | EF-hand calcium binding domain 7 |
| EGF | ENSG00000138798 | 4 | 109912883 | 110013766 | (+) | epidermal growth factor |
| EGFR | ENSG00000146648 | 7 | 55019017 | 55211628 | (+) | epidermal growth factor receptor |

| | | | | | | |
|---|---|---|---|---|---|---|
| EGR3 | ENSG00000179388 | 8 | 22687659 | 22693480 | (-) | early growth response 3 |
| EIF5B | ENSG00000158417 | 2 | 99337389 | 99401326 | (+) | eukaryotic translation initiation factor 5B |
| ENO1 | ENSG00000074800 | 1 | 8861000 | 8879190 | (-) | enolase 1 |
| EOMES | ENSG00000163508 | 3 | 27715949 | 27722711 | (-) | eomesodermin |
| ERAP1 | ENSG00000164307 | 5 | 96760810 | 96808100 | (-) | endoplasmic reticulum aminopeptidase 1 |
| ERAP2 | ENSG00000164308 | 5 | 96875986 | 96919703 | (+) | endoplasmic reticulum aminopeptidase 2 |
| ERN1 | ENSG00000178607 | 17 | 64039080 | 64130819 | (-) | endoplasmic reticulum to nucleus signaling 1 |
| ERP44 | ENSG00000023318 | 9 | 99979185 | 100099052 | (-) | endoplasmic reticulum protein 44 |
| EZH2 | ENSG00000106462 | 7 | 148807257 | 148884321 | (-) | enhancer of zeste 2 polycomb repressive complex 2 subunit |
| FAXDC2 | ENSG00000170271 | 5 | 154818492 | 154859252 | (-) | fatty acid hydroxylase domain containing 2 |

| | | | | | | |
|---|---|---|---|---|---|---|
| FCGR1A | ENSG00000150337 | 1 | 149782671 | 149791675 | (+) | Fc gamma receptor Ia |
| FCGR2A | ENSG00000143226 | 1 | 161505430 | 161524013 | (+) | Fc gamma receptor IIa |
| FCGR3A | ENSG00000203747 | 1 | 161541759 | 161550968 | (-) | Fc gamma receptor IIIa |
| FGB | ENSG00000171564 | 4 | 154563011 | 154572807 | (+) | fibrinogen beta chain |
| FNBP1 | ENSG00000187239 | 9 | 129887187 | 130043189 | (-) | formin binding protein 1 |
| FOS | ENSG00000170345 | 14 | 75278826 | 75283190 | (+) | Fos proto-oncogene, AP-1 transcription factor subunit |
| FOSL1 | ENSG00000175592 | 11 | 65892049 | 65900573 | (-) | FOS like 1, AP-1 transcription factor subunit |
| FOSL2 | ENSG00000075426 | 2 | 28392448 | 28417317 | (+) | FOS like 2, AP-1 transcription factor subunit |
| FOXP3 | ENSG00000049768 | X | 49250438 | 49264800 | (-) | forkhead box P3 |
| FRZB | ENSG00000162998 | 2 | 182833275 | 182866637 | (-) | frizzled related protein |

| | | | | | | |
|---|---|---|---|---|---|---|
| FUT2 | ENSG00000176920 | 19 | 48695971 | 48705951 | (+) | fucosyltransferase 2 (H blood group) |
| GAPDH | ENSG00000111640 | 12 | 6534512 | 6538374 | (+) | glyceraldehyde-3-phosphate dehydrogenase |
| GATA3 | ENSG00000107485 | 10 | 8045378 | 8075198 | (+) | GATA binding protein 3 |
| GBP1 | ENSG00000117228 | 1 | 89051882 | 89065360 | (-) | guanylate binding protein 1 |
| GBP3 | ENSG00000117226 | 1 | 89006666 | 89022894 | (-) | guanylate binding protein 3 |
| GBP5 | ENSG00000154451 | 1 | 89256189 | 89272860 | (-) | guanylate binding protein 5 |
| GEM | ENSG00000164949 | 8 | 94249253 | 94262350 | (-) | GTP binding protein overexpressed in skeletal muscle |
| GINS1 | ENSG00000101003 | 20 | 25391008 | 25452700 | (+) | GINS complex subunit 1 |
| GJB2 | ENSG00000165474 | 13 | 20187463 | 20192938 | (-) | gap junction protein beta 2 |
| GJB6 | ENSG00000121742 | 13 | 20221962 | 20232365 | (-) | gap junction protein beta 6 |

| | | | | | | |
|---|---|---|---|---|---|---|
| GLUL | ENSG00000135821 | 1 | 182378098 | 182392206 | (-) | glutamate-ammonia ligase |
| GOLIM4 | ENSG00000173905 | 3 | 168008689 | 168095924 | (-) | golgi integral membrane protein 4 |
| GPR35 | ENSG00000178623 | 2 | 240605430 | 240633159 | (+) | G protein-coupled receptor 35 |
| GPT | ENSG00000167701 | 8 | 144502973 | 144507174 | (+) | glutamic--pyruvic transaminase |
| GZMA | ENSG00000145649 | 5 | 55102646 | 55110252 | (+) | granzyme A |
| GZMB | ENSG00000100453 | 14 | 24630954 | 24634267 | (-) | granzyme B |
| GZMK | ENSG00000113088 | 5 | 55024256 | 55034570 | (+) | granzyme K |
| HCAR3 | ENSG00000255398 | 12 | 122714756 | 122716811 | (-) | hydroxycarboxylic acid receptor 3 |
| HERC6 | ENSG00000138642 | 4 | 88378739 | 88443097 | (+) | HECT and RLD domain containing E3 ubiquitin protein ligase family member 6 |
| HHAT | ENSG00000054392 | 1 | 210328252 | 210676296 | (+) | hedgehog acyltransferase |

| | | | | | | |
|---|---|---|---|---|---|---|
| HIF1A | ENSG00000100644 | 14 | 61695513 | 61748259 | (+) | hypoxia inducible factor 1 subunit alpha |
| HK2 | ENSG00000159399 | 2 | 74834127 | 74893359 | (+) | hexokinase 2 |
| HLA-A | ENSG00000206503 | 6 | 29941260 | 29949572 | (+) | major histocompatibility complex, class I, A |
| HLA-B | ENSG00000234745 | 6 | 31353872 | 31367067 | (-) | major histocompatibility complex, class I, B |
| HLA-C | ENSG00000204525 | 6 | 31268749 | 31272130 | (-) | major histocompatibility complex, class I, C |
| HLA-DQB1 | ENSG00000179344 | 6 | 32659467 | 32668383 | (-) | major histocompatibility complex, class II, DQ beta 1 |
| HLA-DRB1 | ENSG00000196126 | 6 | 32577902 | 32589848 | (-) | major histocompatibility complex, class II, DR beta 1 |
| HSPA5 | ENSG00000044574 | 9 | 125234853 | 125241382 | (-) | heat shock protein family A (Hsp70) member 5 |
| HSPA6 | ENSG00000173110 | 1 | 161524540 | 161526894 | (+) | heat shock protein family A (Hsp70) member 6 |
| HYAL4 | ENSG00000106302 | 7 | 123828983 | 123877481 | (+) | hyaluronidase 4 |

| | | | | | | |
|---|---|---|---|---|---|---|
| ICAM1 | ENSG00000090339 | 19 | 10271093 | 10286615 | (+) | intercellular adhesion molecule 1 |
| ICOS | ENSG00000163600 | 2 | 203936763 | 203961577 | (+) | inducible T cell costimulator |
| IFI16 | ENSG00000163565 | 1 | 158999968 | 159055155 | (+) | interferon gamma inducible protein 16 |
| IFI6 | ENSG00000126709 | 1 | 27666064 | 27672212 | (-) | interferon alpha inducible protein 6 |
| IFIH1 | ENSG00000115267 | 2 | 162267074 | 162318684 | (-) | interferon induced with helicase C domain 1 |
| IFIT3 | ENSG00000119917 | 10 | 89327307 | 89377473 | (+) | interferon induced protein with tetratricopeptide repeats 3 |
| IFNA1 | ENSG00000197919 | 9 | 21440439 | 21441316 | (+) | interferon alpha 1 |
| IFNG | ENSG00000111537 | 12 | 68154768 | 68159740 | (-) | interferon gamma |
| IFNGR1 | ENSG00000027697 | 6 | 137197483 | 137219449 | (-) | interferon gamma receptor 1 |
| IFNLR1 | ENSG00000185436 | 1 | 24154168 | 24187959 | (-) | interferon lambda receptor 1 |

| | | | | | | |
|---|---|---|---|---|---|---|
| IGF2 | ENSG00000167244 | 11 | 2129112 | 2158391 | (-) | insulin like growth factor 2 |
| IGF2-AS | ENSG00000099869 | 11 | 2140501 | 2148666 | (+) | IGF2 antisense RNA |
| IL10 | ENSG00000136634 | 1 | 206767602 | 206774541 | (-) | interleukin 10 |
| IL12B | ENSG00000113302 | 5 | 159314780 | 159330863 | (-) | interleukin 12B |
| IL13 | ENSG00000169194 | 5 | 132656263 | 132661110 | (+) | interleukin 13 |
| IL15 | ENSG00000164136 | 4 | 141636583 | 141733987 | (+) | interleukin 15 |
| IL17A | ENSG00000112115 | 6 | 52186375 | 52190638 | (+) | interleukin 17A |
| IL17F | ENSG00000112116 | 6 | 52236681 | 52245689 | (-) | interleukin 17F |
| IL17RA | ENSG00000177663 | 22 | 17084954 | 17115693 | (+) | interleukin 17 receptor A |
| IL18 | ENSG00000150782 | 11 | 112143251 | 112164096 | (-) | interleukin 18 |
| IL1A | ENSG00000115008 | 2 | 112773925 | 112784493 | (-) | interleukin 1 alpha |

| | | | | | | |
|---|---|---|---|---|---|---|
| IL1B | ENSG00000125538 | 2 | 112829751 | 112836816 | (-) | interleukin 1 beta |
| IL1F10 | ENSG00000136697 | 2 | 113067970 | 113075843 | (+) | interleukin 1 family member 10 |
| IL1R1 | ENSG00000115594 | 2 | 102064544 | 102179874 | (+) | interleukin 1 receptor type 1 |
| IL1RN | ENSG00000136689 | 2 | 113099315 | 113134016 | (+) | interleukin 1 receptor antagonist |
| IL2 | ENSG00000109471 | 4 | 122451470 | 122456725 | (-) | interleukin 2 |
| IL21 | ENSG00000138684 | 4 | 122610108 | 122621066 | (-) | interleukin 21 |
| IL21-AS1 | ENSG00000227145 | 4 | 122618983 | 122689164 | (+) | IL21 antisense RNA 1 |
| IL22 | ENSG00000127318 | 12 | 68248242 | 68253604 | (-) | interleukin 22 |
| IL23A | ENSG00000110944 | 12 | 56334174 | 56340410 | (+) | interleukin 23 subunit alpha |
| IL23R | ENSG00000162594 | 1 | 67138907 | 67259979 | (+) | interleukin 23 receptor |
| IL2RA | ENSG00000134460 | 10 | 6010689 | 6062370 | (-) | interleukin 2 receptor subunit alpha |

| | | | | | | |
|---|---|---|---|---|---|---|
| IL33 | ENSG00000137033 | 9 | 6215786 | 6257983 | (+) | interleukin 33 |
| IL36RN | ENSG00000136695 | 2 | 113058638 | 113065382 | (+) | interleukin 36 receptor antagonist |
| IL37 | ENSG00000125571 | 2 | 112911165 | 112918882 | (+) | interleukin 37 |
| IL4 | ENSG00000113520 | 5 | 132673986 | 132682678 | (+) | interleukin 4 |
| IL5 | ENSG00000113525 | 5 | 132541445 | 132556838 | (-) | interleukin 5 |
| IL6 | ENSG00000136244 | 7 | 22725884 | 22732002 | (+) | interleukin 6 |
| IL6R | ENSG00000160712 | 1 | 154405193 | 154469450 | (+) | interleukin 6 receptor |
| IL7 | ENSG00000104432 | 8 | 78675743 | 78805523 | (-) | interleukin 7 |
| INS | ENSG00000254647 | 11 | 2159779 | 2161221 | (-) | insulin |
| INS-IGF2 | ENSG00000129965 | 11 | 2132538 | 2161209 | (-) | INS-IGF2 readthrough |
| IRAK1 | ENSG00000184216 | X | 154010506 | 154019902 | (-) | interleukin 1 receptor associated kinase 1 |

| | | | | | | |
|---|---|---|---|---|---|---|
| IRS1 | ENSG00000169047 | 2 | 226731312 | 226799820 | (-) | insulin receptor substrate 1 |
| ITGA2B | ENSG00000005961 | 17 | 44372180 | 44389649 | (-) | integrin subunit alpha 2b |
| ITGAL | ENSG00000005844 | 16 | 30472658 | 30523567 | (+) | integrin subunit alpha L |
| ITGAM | ENSG00000169896 | 16 | 31259967 | 31332892 | (+) | integrin subunit alpha M |
| ITGAX | ENSG00000140678 | 16 | 31355134 | 31382999 | (+) | integrin subunit alpha X |
| JAK2 | ENSG00000096968 | 9 | 4984390 | 5129948 | (+) | Janus kinase 2 |
| JAK3 | ENSG00000105639 | 19 | 17824780 | 17848071 | (-) | Janus kinase 3 |
| JDP2 | ENSG00000140044 | 14 | 75427716 | 75474111 | (+) | Jun dimerization protein 2 |
| JRKL | ENSG00000183340 | 11 | 96389989 | 96507574 | (+) | JRK like |
| JUN | ENSG00000177606 | 1 | 58776845 | 58784048 | (-) | Jun proto-oncogene, AP-1 transcription factor subunit |

| | | | | | | |
|---|---|---|---|---|---|---|
| JUNB | ENSG00000171223 | 19 | 12791486 | 12793315 | (+) | JunB proto-oncogene, AP-1 transcription factor subunit |
| JUND | ENSG00000130522 | 19 | 18279694 | 18281622 | (-) | JunD proto-oncogene, AP-1 transcription factor subunit |
| KANK4 | ENSG00000132854 | 1 | 62236165 | 62319434 | (-) | KN motif and ankyrin repeat domains 4 |
| KDM5B | ENSG00000117139 | 1 | 202724495 | 202808487 | (-) | lysine demethylase 5B |
| KIR2DS1 | ENSG00000283937 | HSCHR19KIR_CA01-TB01_CTG3_1 | 163173 | 177001 | (+) | killer cell immunoglobulin like receptor, two Ig domains and short cytoplasmic tail 1 |
| KIR3DL1 | ENSG00000167633 | 19 | 54816468 | 54830778 | (+) | killer cell immunoglobulin like receptor, three Ig domains and long cytoplasmic tail 1 |
| KIR3DL2 | ENSG00000240403 | 19 | 54850443 | 54867207 | (+) | killer cell immunoglobulin like receptor, three Ig domains and long cytoplasmic tail 2 |
| KLRB1 | ENSG00000111796 | 12 | 9594551 | 9607916 | (-) | killer cell lectin like receptor B1 |
| LAMP1 | ENSG00000185896 | 13 | 113297239 | 113323672 | (+) | lysosomal associated membrane protein 1 |

| | | | | | | |
|---|---|---|---|---|---|---|
| LAMP2 | ENSG00000005893 | X | 120426148 | 120469365 | (-) | lysosomal associated membrane protein 2 |
| LEP | ENSG00000174697 | 7 | 128241278 | 128257629 | (+) | leptin |
| LILRA5 | ENSG00000187116 | 19 | 54307070 | 54313166 | (-) | leukocyte immunoglobulin like receptor A5 |
| LILRB2 | ENSG00000131042 | 19 | 54273812 | 54281184 | (-) | leukocyte immunoglobulin like receptor B2 |
| LINC01185 | ENSG00000228414 | 2 | 60823069 | 60881317 | (-) | long intergenic non-protein coding RNA 1185 |
| LINC01250 | ENSG00000234423 | 2 | 2893569 | 3147934 | (-) | long intergenic non-protein coding RNA 1250 |
| LMO7 | ENSG00000136153 | 13 | 75620434 | 75859870 | (+) | LIM domain 7 |
| LPAL2 | ENSG00000213071 | 6 | 160453428 | 160520269 | (-) | lipoprotein(a) like 2, pseudogene |
| LRRK2 | ENSG00000188906 | 12 | 40196744 | 40369285 | (+) | leucine rich repeat kinase 2 |
| LTA | ENSG00000226979 | 6 | 31572054 | 31574324 | (+) | lymphotoxin alpha |
| LURAP1L | ENSG00000153714 | 9 | 12775020 | 12823060 | (+) | leucine rich adaptor protein 1 like |

| | | | | | | |
|---|---|---|---|---|---|---|
| LURAP1L-AS 1 | ENSG00000235448 | 9 | 12631434 | 12814382 | (-) | LURAP1L antisense RNA 1 |
| LYZ | ENSG00000090382 | 12 | 69348381 | 69354234 | (+) | lysozyme |
| MAF | ENSG00000178573 | 16 | 79585843 | 79600737 | (-) | MAF bZIP transcription factor |
| MBL2 | ENSG00000165471 | 10 | 52765380 | 52772784 | (-) | mannose binding lectin 2 |
| MCAM | ENSG00000076706 | 11 | 119308529 | 119321521 | (-) | melanoma cell adhesion molecule |
| MCL1 | ENSG00000143384 | 1 | 150560895 | 150579738 | (-) | MCL1 apoptosis regulator, BCL2 family member |
| MEFV | ENSG00000103313 | 16 | 3242027 | 3256633 | (-) | MEFV innate immunity regulator, pyrin |
| MICA | ENSG00000204520 | 6 | 31399784 | 31415315 | (+) | MHC class I polypeptide-related sequence A |
| MIR146A | ENSG00000283733 | 5 | 160485352 | 160485450 | (+) | microRNA 146a |
| MIR21 | ENSG00000284190 | 17 | 59841266 | 59841337 | (+) | microRNA 21 |

| | | | | | | |
|---|---|---|---|---|---|---|
| MIX23 | ENSG00000160124 | 3 | 122359591 | 122383231 | (-) | mitochondrial matrix import factor 23 |
| MMP1 | ENSG00000196611 | 11 | 102789401 | 102798160 | (-) | matrix metallopeptidase 1 |
| MMP3 | ENSG00000149968 | 11 | 102835801 | 102843609 | (-) | matrix metallopeptidase 3 |
| MMP7 | ENSG00000137673 | 11 | 102520508 | 102530750 | (-) | matrix metallopeptidase 7 |
| MMP9 | ENSG00000100985 | 20 | 46008908 | 46016561 | (+) | matrix metallopeptidase 9 |
| MRPS23 | ENSG00000181610 | 17 | 57834781 | 57850056 | (-) | mitochondrial ribosomal protein S23 |
| MSN | ENSG00000147065 | X | 65588377 | 65741931 | (+) | moesin |
| MT-CO2 | ENSG00000198712 | M | 7586 | 8269 | (+) | cytochrome c oxidase subunit II / A MESMA COISA QUE COX2 |
| MTHFR | ENSG00000177000 | 1 | 11785723 | 11806455 | (-) | methylenetetrahydrofolate reductase |
| MUCL1 | ENSG00000172551 | 12 | 54830518 | 54896008 | (+) | mucin like 1 |

| | | | | | | |
|---|---|---|---|---|---|---|
| MX1 | ENSG00000157601 | 21 | 41420020 | 41470071 | (+) | MX dynamin like GTPase 1 |
| MYC | ENSG00000136997 | 8 | 127735434 | 127742951 | (+) | MYC proto-oncogene, bHLH transcription factor |
| MYNN | ENSG00000085274 | 3 | 169773396 | 169789716 | (+) | myoneurin |
| NABP1 | ENSG00000173559 | 2 | 191678068 | 191741097 | (+) | nucleic acid binding protein 1 |
| NAMPT | ENSG00000105835 | 7 | 106248298 | 106285966 | (-) | nicotinamide phosphoribosyltransferase |
| NCOA7 | ENSG00000111912 | 6 | 125781161 | 125932034 | (+) | nuclear receptor coactivator 7 |
| NDUFS1 | ENSG00000023228 | 2 | 206114817 | 206159509 | (-) | NADH:ubiquinone oxidoreductase core subunit S1 |
| NFKB1 | ENSG00000109320 | 4 | 102501330 | 102617302 | (+) | nuclear factor kappa B subunit 1 |
| NKG7 | ENSG00000105374 | 19 | 51371606 | 51372701 | (-) | natural killer cell granule protein 7 |
| NLRP3 | ENSG00000162711 | 1 | 247332331 | 247449108 | (+) | NLR family pyrin domain containing 3 |
| NMI | ENSG00000123609 | 2 | 151270470 | 151289894 | (-) | N-myc and STAT interactor |

| | | | | | | |
|---|---|---|---|---|---|---|
| NOD2 | ENSG00000167207 | 16 | 50693588 | 50733077 | (+) | nucleotide binding oligomerization domain containing 2 |
| NOS2 | ENSG00000007171 | 17 | 27756766 | 27800529 | (-) | nitric oxide synthase 2 |
| NOXRED1 | ENSG00000165555 | 14 | 77394021 | 77423523 | (-) | NADP dependent oxidoreductase domain containing 1 |
| NPEPPS | ENSG00000141279 | 17 | 47522942 | 47624665 | (+) | aminopeptidase puromycin sensitive |
| NRG1 | ENSG00000157168 | 8 | 31639222 | 32855666 | (+) | neuregulin 1 |
| NT5C3A | ENSG00000122643 | 7 | 33014113 | 33062796 | (-) | 5'-nucleotidase, cytosolic IIIA |
| OLR1 | ENSG00000173391 | 12 | 10158301 | 10172138 | (-) | oxidized low density lipoprotein receptor 1 |
| OSMR | ENSG00000145623 | 5 | 38845858 | 38945596 | (+) | oncostatin M receptor |
| PDCD1 | ENSG00000188389 | 2 | 241849884 | 241858894 | (-) | programmed cell death 1 |
| PDLIM7 | ENSG00000196923 | 5 | 177483394 | 177497606 | (-) | PDZ and LIM domain 7 |

| | | | | | | |
|---|---|---|---|---|---|---|
| PER1 | ENSG00000179094 | 17 | 8140467 | 8156506 | (-) | period circadian regulator 1 |
| PFDN4 | ENSG00000101132 | 20 | 54207921 | 54228052 | (+) | prefoldin subunit 4 |
| PFDN5 | ENSG00000123349 | 12 | 53295291 | 53299452 | (+) | prefoldin subunit 5 |
| PFKL | ENSG00000141959 | 21 | 44300051 | 44327376 | (+) | phosphofructokinase, liver type |
| PGD | ENSG00000142657 | 1 | 10398592 | 10420511 | (+) | phosphogluconate dehydrogenase |
| PGK1 | ENSG00000102144 | X | 77910739 | 78129295 | (+) | phosphoglycerate kinase 1 |
| PHEX | ENSG00000102174 | X | 22032325 | 22494713 | (+) | phosphate regulating endopeptidase X-linked |
| PI3 | ENSG00000124102 | 20 | 45174902 | 45176544 | (+) | peptidase inhibitor 3 |
| PIK3CD | ENSG00000171608 | 1 | 9629889 | 9729114 | (+) | phosphatidylinositol-4,5-bisphosphate 3-kinase catalytic subunit delta |
| PINK1 | ENSG00000158828 | 1 | 20633458 | 20651511 | (+) | PTEN induced kinase 1 |

| | | | | | | |
|---|---|---|---|---|---|---|
| PLA2G4D | ENSG00000159337 | 15 | 42067009 | 42094562 | (-) | phospholipase A2 group IVD |
| PLCG1 | ENSG00000124181 | 20 | 41136960 | 41196801 | (+) | phospholipase C gamma 1 |
| PLG | ENSG00000122194 | 6 | 160702194 | 160754097 | (+) | plasminogen |
| PLIN5 | ENSG00000214456 | 19 | 4522531 | 4535224 | (-) | perilipin 5 |
| PLS1 | ENSG00000120756 | 3 | 142596393 | 142713664 | (+) | plastin 1 |
| PPARD | ENSG00000112033 | 6 | 35342558 | 35428191 | (+) | peroxisome proliferator activated receptor delta |
| PPARG | ENSG00000132170 | 3 | 12287368 | 12434356 | (+) | peroxisome proliferator activated receptor gamma |
| PPARGC1A | ENSG00000109819 | 4 | 23755041 | 23904089 | (-) | PPARG coactivator 1 alpha |
| PPARGC1B | ENSG00000155846 | 5 | 149730298 | 149855022 | (+) | PPARG coactivator 1 beta |
| PRDM1 | ENSG00000057657 | 6 | 105993463 | 106109939 | (+) | PR/SET domain 1 |

| | | | | | | |
|---|---|---|---|---|---|---|
| PRF1 | ENSG00000180644 | 10 | 70597348 | 70602759 | (-) | perforin 1 |
| PRTN3 | ENSG00000196415 | 19 | 840999 | 848175 | (+) | proteinase 3 |
| PSG2 | ENSG00000242221 | 19 | 43064209 | 43083045 | (-) | pregnancy specific beta-1-glycoprotein 2 |
| PSMC2 | ENSG00000161057 | 7 | 103328570 | 103370346 | (+) | proteasome 26S subunit, ATPase 2 |
| PSMD7 | ENSG00000103035 | 16 | 74296814 | 74306288 | (+) | proteasome 26S subunit, non-ATPase 7 |
| PSME2 | ENSG00000100911 | 14 | 24143362 | 24147570 | (-) | proteasome activator subunit 2 |
| PTGS1 | ENSG00000095303 | 9 | 122370530 | 122395703 | (+) | prostaglandin-endoperoxide synthase 1 |
| PTGS2 | ENSG00000073756 | 1 | 186671791 | 186680922 | (-) | prostaglandin-endoperoxide synthase 2 |
| PTH | ENSG00000152266 | 11 | 13492054 | 13496181 | (-) | parathyroid hormone |
| PTPN22 | ENSG00000134242 | 1 | 113813811 | 113871753 | (-) | protein tyrosine phosphatase non-receptor type 22 |

| | | | | | | |
|---|---|---|---|---|---|---|
| PTX3 | ENSG00000163661 | 3 | 157436850 | 157443633 | (+) | pentraxin 3 |
| PYGL | ENSG00000100504 | 14 | 50857891 | 50944483 | (-) | glycogen phosphorylase L |
| RAC1 | ENSG00000136238 | 7 | 6374527 | 6403967 | (+) | Rac family small GTPase 1 |
| RBM45 | ENSG00000155636 | 2 | 178112424 | 178139011 | (+) | RNA binding motif protein 45 |
| REL | ENSG00000162924 | 2 | 60881491 | 60931612 | (+) | REL proto-oncogene, NF-kB subunit |
| RETN | ENSG00000104918 | 19 | 7669049 | 7670455 | (+) | resistin |
| RGPD6 | ENSG00000183054 | 2 | 110513802 | 110610840 | (-) | RANBP2 like and GRIP domain containing 6 |
| RIT1 | ENSG00000143622 | 1 | 155897808 | 155911404 | (-) | Ras like without CAAX 1 |
| RORC | ENSG00000143365 | 1 | 151806071 | 151831845 | (-) | RAR related orphan receptor C |
| RPL15 | ENSG00000174748 | 3 | 23916591 | 23924374 | (+) | ribosomal protein L15 |
| RPL36AL | ENSG00000165502 | 14 | 49618530 | 49620626 | (-) | ribosomal protein L36a like |

| | | | | | | |
|---|---|---|---|---|---|---|
| RPL41 | ENSG00000229117 | 12 | 56116590 | 56117967 | (+) | ribosomal protein L41 |
| RPL7 | ENSG00000147604 | 8 | 73290242 | 73295789 | (-) | ribosomal protein L7 |
| RPS19 | ENSG00000105372 | 19 | 41860255 | 41872925 | (+) | ribosomal protein S19 |
| RPS21 | ENSG00000171858 | 20 | 62387103 | 62388520 | (+) | ribosomal protein S21 |
| RPS26 | ENSG00000197728 | 12 | 56041351 | 56044697 | (+) | ribosomal protein S26 |
| RPS6KB1 | ENSG00000108443 | 17 | 59893046 | 59950574 | (+) | ribosomal protein S6 kinase B1 |
| RPS7 | ENSG00000171863 | 2 | 3575260 | 3580920 | (+) | ribosomal protein S7 |
| RSAD2 | ENSG00000134321 | 2 | 6865557 | 6898239 | (+) | radical S-adenosyl methionine domain containing 2 |
| RUNX2 | ENSG00000124813 | 6 | 45328157 | 45664349 | (+) | RUNX family transcription factor 2 |
| RUNX3 | ENSG00000020633 | 1 | 24899511 | 24965121 | (-) | RUNX family transcription factor 3 |

| | | | | | | |
|---|---|---|---|---|---|---|
| S100A12 | ENSG00000163221 | 1 | 153373711 | 153375621 | (-) | S100 calcium binding protein A12 |
| S100A8 | ENSG00000143546 | 1 | 153390032 | 153391073 | (-) | S100 calcium binding protein A8 |
| S100A9 | ENSG00000163220 | 1 | 153357854 | 153361023 | (+) | S100 calcium binding protein A9 |
| S100P | ENSG00000163993 | 4 | 6693878 | 6697170 | (+) | S100 calcium binding protein P |
| SAA1 | ENSG00000173432 | 11 | 18266260 | 18269977 | (+) | serum amyloid A1 |
| SAMD9 | ENSG00000205413 | 7 | 93099513 | 93118023 | (-) | sterile alpha motif domain containing 9 |
| SAR1A | ENSG00000079332 | 10 | 70147289 | 70170523 | (-) | secretion associated Ras related GTPase 1A |
| SCN1A | ENSG00000144285 | 2 | 165984641 | 166182806 | (-) | sodium voltage-gated channel alpha subunit 1 |
| SEC14L2 | ENSG00000100003 | 22 | 30396941 | 30425303 | (+) | SEC14 like lipid binding 2 |
| SEC24B | ENSG00000138802 | 4 | 109433772 | 109540896 | (+) | SEC24 homolog B, COPII coat complex component |

| | | | | | | |
|---|---|---|---|---|---|---|
| SELL | ENSG00000188404 | 1 | 169690665 | 169711702 | (-) | selectin L |
| SERPINA1 | ENSG00000197249 | 14 | 94376747 | 94390693 | (-) | serpin family A member 1 |
| SERPINB1 | ENSG00000021355 | 6 | 2832332 | 2841959 | (-) | serpin family B member 1 |
| SERPINE1 | ENSG00000106366 | 7 | 101127104 | 101139247 | (+) | serpin family E member 1 |
| SF3B1 | ENSG00000115524 | 2 | 197388515 | 197435079 | (-) | splicing factor 3b subunit 1 |
| SF3B3 | ENSG00000189091 | 16 | 70523791 | 70577670 | (+) | splicing factor 3b subunit 3 |
| SGK1 | ENSG00000118515 | 6 | 134169248 | 134318112 | (-) | serum/glucocorticoid regulated kinase 1 |
| SH3BGRL3 | ENSG00000142669 | 1 | 26280086 | 26281522 | (+) | SH3 domain binding glutamate rich protein like 3 |
| SIAH1 | ENSG00000196470 | 16 | 48356364 | 48448402 | (-) | siah E3 ubiquitin protein ligase 1 |
| SLC1A2 | ENSG00000110436 | 11 | 35251205 | 35420063 | (-) | solute carrier family 1 member 2 |
| SLC2A3 | ENSG00000059804 | 12 | 7919230 | 8019007 | (-) | solute carrier family 2 member 3 |

| | | | | | | |
|---|---|---|---|---|---|---|
| SLC51B | ENSG00000186198 | 15 | 65045387 | 65053397 | (+) | SLC51 subunit beta |
| SLC7A11 | ENSG00000151012 | 4 | 138164097 | 138242349 | (-) | solute carrier family 7 member 11 |
| SLC7A5 | ENSG00000103257 | 16 | 87830016 | 87869507 | (-) | solute carrier family 7 member 5 |
| SMAD3 | ENSG00000166949 | 15 | 67063763 | 67195173 | (+) | SMAD family member 3 |
| SMARCA4 | ENSG00000127616 | 19 | 10960932 | 11079426 | (+) | SWI/SNF related, matrix associated, actin dependent regulator of chromatin, subfamily a, member 4 |
| SMOX | ENSG00000088826 | 20 | 4120980 | 4187747 | (+) | spermine oxidase |
| SOCS1 | ENSG00000185338 | 16 | 11254417 | 11256204 | (-) | suppressor of cytokine signaling 1 |
| SOD2 | ENSG00000291237 | 6 | 159669069 | 159762529 | (-) | superoxide dismutase 2 |
| SOST | ENSG00000167941 | 17 | 43753738 | 43758791 | (-) | sclerostin |
| SOX4 | ENSG00000124766 | 6 | 21593751 | 21598619 | (+) | SRY-box transcription factor 4 |

| | | | | | | |
|---|---|---|---|---|---|---|
| SP7 | ENSG00000170374 | 12 | 53326575 | 53345315 | (-) | Sp7 transcription factor |
| SPCS3 | ENSG00000129128 | 4 | 176319966 | 176332245 | (+) | signal peptidase complex subunit 3 |
| SPON2 | ENSG00000159674 | 4 | 1166932 | 1208962 | (-) | spondin 2 |
| SPP1 | ENSG00000118785 | 4 | 87975667 | 87983532 | (+) | secreted phosphoprotein 1 |
| SSR1 | ENSG00000124783 | 6 | 7268306 | 7347446 | (-) | signal sequence receptor subunit 1 |
| STAT1 | ENSG00000115415 | 2 | 190908460 | 191020960 | (-) | signal transducer and activator of transcription 1 |
| STAT3 | ENSG00000168610 | 17 | 42313324 | 42388568 | (-) | signal transducer and activator of transcription 3 |
| STIM1 | ENSG00000167323 | 11 | 3854527 | 4093210 | (+) | stromal interaction molecule 1 |
| SUOX | ENSG00000139531 | 12 | 55997180 | 56006641 | (+) | sulfite oxidase |
| SYT1 | ENSG00000067715 | 12 | 78863993 | 79452008 | (+) | synaptotagmin 1 |
| TALDO1 | ENSG00000177156 | 11 | 747415 | 765012 | (+) | transaldolase 1 |

| | | | | | | |
|---|---|---|---|---|---|---|
| TBX21 | ENSG00000073861 | 17 | 47733236 | 47746122 | (+) | T-box transcription factor 21 |
| TEK | ENSG00000120156 | 9 | 27109141 | 27230174 | (+) | TEK receptor tyrosine kinase |
| TFPI | ENSG00000003436 | 2 | 187464230 | 187565760 | (-) | tissue factor pathway inhibitor |
| TGFA | ENSG00000163235 | 2 | 70447284 | 70554193 | (-) | transforming growth factor alpha |
| TGFB1 | ENSG00000105329 | 19 | 41301587 | 41353922 | (-) | transforming growth factor beta 1 |
| TGFBR3 | ENSG00000069702 | 1 | 91680343 | 91906335 | (-) | transforming growth factor beta receptor 3 |
| TIMP1 | ENSG00000102265 | X | 47582408 | 47586789 | (+) | TIMP metallopeptidase inhibitor 1 |
| TLR2 | ENSG00000137462 | 4 | 153684050 | 153706260 | (+) | toll like receptor 2 |
| TLR3 | ENSG00000164342 | 4 | 186068911 | 186088073 | (+) | toll like receptor 3 |
| TLR4 | ENSG00000136869 | 9 | 117704175 | 117724735 | (+) | toll like receptor 4 |
| TLR9 | ENSG00000239732 | 3 | 52221080 | 52225645 | (-) | toll like receptor 9 |

| | | | | | | |
|---|---|---|---|---|---|---|
| TMBIM6 | ENSG00000139644 | 12 | 49707725 | 49764934 | (+) | transmembrane BAX inhibitor motif containing |
| TMEM45A | ENSG00000181458 | 3 | 100492619 | 100577444 | (+) | transmembrane protein 45A |
| TMPRSS11B | ENSG00000185873 | 4 | 68226653 | 68245694 | (-) | transmembrane serine protease 11B |
| TNF | ENSG00000232810 | 6 | 31575565 | 31578336 | (+) | tumor necrosis factor |
| TNFAIP3 | ENSG00000118503 | 6 | 137867214 | 137883314 | (+) | TNF alpha induced protein 3 |
| TNFAIP6 | ENSG00000123610 | 2 | 151357592 | 151380046 | (+) | TNF alpha induced protein 6 |
| TNFAIP8 | ENSG00000145779 | 5 | 119268692 | 119399688 | (+) | TNF alpha induced protein 8 |
| TNFRSF10A | ENSG00000104689 | 8 | 23190452 | 23225102 | (-) | TNF receptor superfamily member 10a |
| TNFRSF1A | ENSG00000067182 | 12 | 6328757 | 6342114 | (-) | TNF receptor superfamily member 1A |
| TNFRSF1B | ENSG00000028137 | 1 | 12166991 | 12209228 | (+) | TNF receptor superfamily member 1B |
| TNFRSF9 | ENSG00000049249 | 1 | 7915871 | 7943165 | (-) | TNF receptor superfamily member 9 |

| | | | | | | |
|---|---|---|---|---|---|---|
| TNFSF10 | ENSG00000121858 | 3 | 172505508 | 172523475 | (-) | TNF superfamily member 10 |
| TNFSF11 | ENSG00000120659 | 13 | 42562736 | 42608013 | (+) | TNF superfamily member 11 |
| TNFSF13B | ENSG00000102524 | 13 | 108251240 | 108308484 | (+) | TNF superfamily member 13b |
| TNFSF15 | ENSG00000181634 | 9 | 114784652 | 114806039 | (-) | TNF superfamily member 15 |
| TNIP1 | ENSG00000145901 | 5 | 151029942 | 151093577 | (-) | TNFAIP3 interacting protein 1 |
| TOMM5 | ENSG00000175768 | 9 | 37582646 | 37592604 | (-) | translocase of outer mitochondrial membrane 5 |
| TOMM7 | ENSG00000196683 | 7 | 22812628 | 22822849 | (-) | translocase of outer mitochondrial membrane 7 |
| TPST2 | ENSG00000128294 | 22 | 26521996 | 26596717 | (-) | tyrosylprotein sulfotransferase 2 |
| TPT1 | ENSG00000133112 | 13 | 45333471 | 45341284 | (-) | tumor protein, translationally-controlled 1 |
| TRAF2 | ENSG00000127191 | 9 | 136881912 | 136929221 | (+) | TNF receptor associated factor 2 |
| TRAF3IP2 | ENSG00000056972 | 6 | 111555381 | 111606906 | (-) | TRAF3 interacting protein 2 |

| | | | | | | |
|---|---|---|---|---|---|---|
| TRAF4 | ENSG00000076604 | 17 | 28744011 | 28750956 | (+) | TNF receptor associated factor 4 |
| TRAF5 | ENSG00000082512 | 1 | 211326615 | 211374946 | (+) | TNF receptor associated factor 5 |
| TRBV20OR9-2 | ENSG00000205274 | 9 | 33617845 | 33618508 | (+) | T cell receptor beta variable 20/OR9-2 (non-functional) |
| TRIM22 | ENSG00000132274 | 11 | 5689610 | 5737089 | (+) | tripartite motif containing 22 |
| TRIM69 | ENSG00000185880 | 15 | 44728988 | 44767829 | (+) | tripartite motif containing 69 |
| TTC39B | ENSG00000155158 | 9 | 15163622 | 15307360 | (-) | tetratricopeptide repeat domain 39B |
| TYK2 | ENSG00000105397 | 19 | 10350528 | 10380608 | (-) | tyrosine kinase 2 |
| TYMP | ENSG00000025708 | 22 | 50525752 | 50530032 | (-) | thymidine phosphorylase |
| UBE2L3 | ENSG00000185651 | 22 | 21549447 | 21624034 | (+) | ubiquitin conjugating enzyme E2 L3 |
| UQCR10 | ENSG00000184076 | 22 | 29767369 | 29770413 | (+) | ubiquinol-cytochrome c reductase, complex III subunit X |

| | | | | | | |
|---|---|---|---|---|---|---|
| UTP11 | ENSG00000183520 | 1 | 38009258 | 38024820 | (+) | UTP11 small subunit processome component |
| VCAM1 | ENSG00000162692 | 1 | 100719742 | 100739045 | (+) | vascular cell adhesion molecule 1 |
| VDR | ENSG00000111424 | 12 | 47841537 | 47943048 | (-) | vitamin D receptor |
| VEGFA | ENSG00000112715 | 6 | 43770184 | 43786487 | (+) | vascular endothelial growth factor A |
| VIM | ENSG00000026025 | 10 | 17228241 | 17237593 | (+) | vimentin |
| WDR1 | ENSG00000071127 | 4 | 10068089 | 10116972 | (-) | WD repeat domain 1 |
| WNK1 | ENSG00000060237 | 12 | 752579 | 911452 | (+) | WNK lysine deficient protein kinase 1 |
| WWOX | ENSG00000186153 | 16 | 78099400 | 79212667 | (+) | WW domain containing oxidoreductase |
| XAF1 | ENSG00000132530 | 17 | 6755447 | 6775647 | (+) | XIAP associated factor 1 |
| XBP1 | ENSG00000100219 | 22 | 28794555 | 28800597 | (-) | X-box binding protein 1 |
| YOD1 | ENSG00000180667 | 1 | 207043849 | 207052980 | (-) | YOD1 deubiquitinase |

| | | | | | | |
|---|---|---|---|---|---|---|
| YWHAB | ENSG00000166913 | 20 | 44885702 | 44908532 | (+) | tyrosine 3-monooxygenase/tryptophan 5-monooxygenase activation protein beta |
| ZC3H12A | ENSG00000163874 | 1 | 37474580 | 37484377 | (+) | zinc finger CCCH-type containing 12A |
| ZFP36 | ENSG00000128016 | 19 | 39406847 | 39409407 | (+) | ZFP36 ring finger protein |
| ZMIZ1 | ENSG00000108175 | 10 | 79068966 | 79316519 | (+) | zinc finger MIZ-type containing 1 |
| ZNF316 | ENSG00000205903 | 7 | 6637318 | 6658279 | (+) | zinc finger protein 316 |
| ZNF415 | ENSG00000170954 | 19 | 53107879 | 53133077 | (-) | zinc finger protein 415 |
| ZNF483 | ENSG00000173258 | 9 | 111525159 | 111577844 | (+) | zinc finger protein 483 |
