## Supplementary Table 2 for "Beyond the Shared Inflammatory Axis: Differentiating Molecular Signatures in Psoriatic Arthritis and Ankylosing Spondylitis through Integrated Omics"

**Supplementary Table 2 - Gene expression profile conserved in non-lesion skin states (PsA and AS) compared to psoriatic lesions.**

| Upregulated | Downregulated |
|---|---|
| KANK4 | TNFRSF9 |
| TGFBR3 | IFI6 |
| RORC | ZC3H12A |
| CRB1 | GBP1 |
| IL37 | GBP5 |
| ABCB11 | FCGR1A |
| FRZB | S100A9 |
| IRS1 | S100A12 |
| PPARG | S100A8 |
| PPARGC1A | AIM2 |
| ALB | FCGR3A |
| EGF | SELL |
| IL5 | PTGS2 |
| FAXDC2 | RSAD2 |
| CMAHP | CD8A |
| ZNF316 | IL1B |
| EGFR | IL36RN |
| CALD1 | STAT1 |
| CSMD1 | CTLA4 |
| GPT | ICOS |
| ZNF483 | CXCR2 |
| GATA3 | CCL20 |
| DKK1 | PDCD1 |
| SLC1A2 | ACKR2 |
| CCND1 | CD80 |
| JRKL | S100P |
| MUCL1 | CD38 |
| CAPS2 | CXCL8 |
| SYT1 | CXCL2 |
| CYP1A1 | CXCL10 |
| AKAP13 | CXCL13 |
| EFCAB13 | HERC6 |
| ZNF415 | IL21 |
| | IL21-AS1 |
| | SLC7A11 |
| | GZMA |

| | |
|---|---|
| | IL12B |
| | SERPINB1 |
| | IL17A |
| | IL17F |
| | PRDM1 |
| | IL6 |
| | NT5C3A |
| | SAMD9 |
| | NAMPT |
| | HYAL4 |
| | CD274 |
| | IL2RA |
| | PRF1 |
| | IFIT3 |
| | FOSL1 |
| | MMP1 |
| | CLEC4D |
| | OLR1 |
| | IFNG |
| | IL22 |
| | HCAR3 |
| | GJB2 |
| | GJB6 |
| | GZMB |
| | SERPINA1 |
| | PLA2G4D |
| | MEFV |
| | NOD2 |
| | NOS2 |
| | CCL3 |
| | CCR7 |
| | ACP7 |
| | FUT2 |
| | NKG7 |
| | LILRA5 |
| | PI3 |
| | MX1 |
| | APOL1 |
| | TYMP |

*The table details the 33 overexpressed genes and 75 underexpressed genes that are common when non-lesional PsA skin and AS skin are independently compared to PsA lesions. The analysis was based on the GSE186063 dataset.*
