## Supplementary Table 3 for "Beyond the Shared Inflammatory Axis: Differentiating Molecular Signatures in Psoriatic Arthritis and Ankylosing Spondylitis through Integrated Omics"

**Supplementary Table 3 - Associations between Genes and Transcription Factors Identified by Gene Expression Analysis for Psoriatic Arthritis (PsA) and Ankylosing Spondylitis (AS)**

| Gene | Transcription Factors | Occurrences | Datasets |
|---|---|---|---|
| CCL20 | IRF8 | 236 | Non lesion PsA vs Lesion PsA GSE186063 |
| CXCL8 | YY1 | 204 | Non lesion PsA vs Lesion PsA GSE186063 |
| CCR7 | RREB1 | 196 | Non lesion PsA vs Lesion PsA GSE186063 |
| CCND1 | GSC | 181 | Non lesion PsA vs Lesion PsA GSE186063 |
| GJB6 | MZF1(var.2) | 176 | Non lesion PsA vs Lesion PsA GSE186063 |
| CXCL2 | POU2F2 | 173 | Non lesion PsA vs Lesion PsA GSE186063 |
| PI3 | ELF5 | 170 | Non lesion PsA vs Lesion PsA GSE186063 |
| IL5 | STAT3 | 168 | Non lesion PsA vs Lesion PsA GSE186063 |

| | | | |
|---|---|---|---|
| FUT2 | FOS | 167 | Non lesion PsA vs Lesion PsA GSE186063 |
| IFI6 | FOSL2 | 167 | Non lesion PsA vs Lesion PsA GSE186063 |
| NMI | NKX2-3 | 166 | Non lesion PsA vs Lesion PsA GSE186063 |
| PRF1 | FOXL1 | 165 | Non lesion PsA vs Lesion PsA GSE186063 |
| DKK1 | RARA::RXRA | 165 | Non lesion PsA vs Lesion PsA GSE186063 |
| TNFRSF9 | SREBF2 | 164 | Non lesion PsA vs Lesion PsA GSE186063 |
| GZMA | STAT1::STAT2 | 163 | Non lesion PsA vs Lesion PsA GSE186063 |
| TGFBR3 | MLXIPL | 161 | Non lesion PsA vs Lesion PsA GSE186063 |
| ZNF316 | FOXF2 | 160 | Non lesion PsA vs Lesion PsA GSE186063 |
| FCGR3A | INSM1 | 159 | Non lesion PsA vs Lesion PsA GSE186063 |

| | | | |
|---|---|---|---|
| PINK1 | LHX6 | 159 | Non lesion PsA vs Lesion PsA GSE186063 |
| SAMD9 | ETV6 | 158 | Non lesion PsA vs Lesion PsA GSE186063 |
| EFCAB13 | SOX9 | 157 | Non lesion PsA vs Lesion PsA GSE186063 |
| MEFV | NFATC3 | 157 | Non lesion PsA vs Lesion PsA GSE186063 |
| PLA2G4D | MAF::NFE2 | 154 | Non lesion PsA vs Lesion PsA GSE186063 |
| PLIN5 | SHOX | 154 | Non lesion PsA vs Lesion PsA GSE186063 |
| PPARG | EN2 | 154 | Non lesion PsA vs Lesion PsA GSE186063 |
| RORC | HEY2 | 154 | Non lesion PsA vs Lesion PsA GSE186063 |
| IL12B | SREBF1 | 152 | Non lesion PsA vs Lesion PsA GSE186063 |
| PRDM1 | IRF9 | 152 | Non lesion PsA vs Lesion PsA GSE186063 |

| | | | |
|---|---|---|---|
| FAXDC2 | MAX::MYC | 150 | Non lesion PsA vs Lesion PsA GSE186063 |
| WNK1 | MEOX1 | 150 | Non lesion PsA vs Lesion PsA GSE186063 |
| MUCL1 | RORA(var.2) | 149 | Non lesion PsA vs Lesion PsA GSE186063 |
| IL17F | PBX1 | 148 | Non lesion PsA vs Lesion PsA GSE186063 |
| SLC7A11 | ARNT::HIF1A | 146 | Non lesion PsA vs Lesion PsA GSE186063 |
| APOL6 | NEUROD2 | 146 | Non lesion PsA vs Lesion PsA GSE186063 |
| IL6 | GATA1::TAL1 | 145 | Non lesion PsA vs Lesion PsA GSE186063 |
| SERPINB1 | EWSR1-FLI1 | 145 | Non lesion PsA vs Lesion PsA GSE186063 |
| IRS1 | DMRT3 | 143 | Non lesion PsA vs Lesion PsA GSE186063 |
| IL37 | MEF2C | 140 | Non lesion PsA vs Lesion PsA GSE186063 |

| | | | |
|---|---|---|---|
| OLR1 | CENPB | 140 | Non lesion PsA vs Lesion PsA GSE186063 |
| NOD2 | JDP2(var.2) | 140 | Non lesion PsA vs Lesion PsA GSE186063 |
| CTLA4 | SRY | 139 | Non lesion PsA vs Lesion PsA GSE186063 |
| RSAD2 | NKX2-8 | 139 | Non lesion PsA vs Lesion PsA GSE186063 |
| GJB2 | JUND(var.2) | 138 | Non lesion PsA vs Lesion PsA GSE186063 |
| AKAP13 | PAX6 | 136 | Non lesion PsA vs Lesion PsA GSE186063 |
| LYZ | LBX1 | 136 | Non lesion PsA vs Lesion PsA GSE186063 |
| SYT1 | MLX | 136 | Non lesion PsA vs Lesion PsA GSE186063 |
| S100A8 | HINFP | 135 | Non lesion PsA vs Lesion PsA GSE186063 |
| IL2RA | STAT1 | 135 | Non lesion PsA vs Lesion PsA GSE186063 |

| | | | |
|---|---|---|---|
| IL36RN | JUN(var.2) | 135 | Non lesion PsA vs Lesion PsA GSE186063 |
| JRKL | NFIX | 135 | Non lesion PsA vs Lesion PsA GSE186063 |
| PPARGC1A | PPARG | 134 | Non lesion PsA vs Lesion PsA GSE186063 |
| GBP5 | RORA | 134 | Non lesion PsA vs Lesion PsA GSE186063 |
| IL17A | THAP1 | 134 | Non lesion PsA vs Lesion PsA GSE186063 |
| IL21 | GRHL1 | 134 | Non lesion PsA vs Lesion PsA GSE186063 |
| CRB1 | NR4A2 | 133 | Non lesion PsA vs Lesion PsA GSE186063 |
| MMP1 | POU6F1 | 133 | Non lesion PsA vs Lesion PsA GSE186063 |
| XAF1 | ESX1 | 133 | Non lesion PsA vs Lesion PsA GSE186063 |
| HYAL4 | PLAG1 | 132 | Non lesion PsA vs Lesion PsA GSE186063 |

| | | | |
|---|---|---|---|
| NT5C3A | DBP | 132 | Non lesion PsA vs Lesion PsA GSE186063 |
| ALB | REST | 131 | Non lesion PsA vs Lesion PsA GSE186063 |
| EGR3 | CTCF | 130 | Non lesion PsA vs Lesion PsA GSE186063 |
| IL23A | ELF4 | 130 | Non lesion PsA vs Lesion PsA GSE186063 |
| NOS2 | BHLHE41 | 129 | Non lesion PsA vs Lesion PsA GSE186063 |
| PTGS2 | GCM1 | 129 | Non lesion PsA vs Lesion PsA GSE186063 |
| APOL1 | NFIA | 129 | Non lesion PsA vs Lesion PsA GSE186063 |
| CYP1A1 | TAL1::TCF3 | 128 | Non lesion PsA vs Lesion PsA GSE186063 |
| GATA3 | FOXH1 | 128 | Non lesion PsA vs Lesion PsA GSE186063 |
| IL22 | CREB3 | 128 | Non lesion PsA vs Lesion PsA GSE186063 |

| | | | |
|---|---|---|---|
| SELL | ISX | 128 | Non lesion PsA vs Lesion PsA GSE186063 |
| CENPK | JDP2 | 128 | Non lesion PsA vs Lesion PsA GSE186063 |
| YOD1 | FOXI1 | 126 | Non lesion PsA vs Lesion PsA GSE186063 |
| PLG | EN1 | 125 | Non lesion PsA vs Lesion PsA GSE186063 |
| EGF | NFIC::TLX1 | 125 | Non lesion PsA vs Lesion PsA GSE186063 |
| ZNF415 | FOXO3 | 125 | Non lesion PsA vs Lesion PsA GSE186063 |
| CD274 | NRF1 | 125 | Non lesion PsA vs Lesion PsA GSE186063 |
| GZMB | TCF7L2 | 125 | Non lesion PsA vs Lesion PsA GSE186063 |
| CAPS2 | NEUROG2 | 125 | Non lesion PsA vs Lesion PsA GSE186063 |
| HCAR3 | NHLH1 | 123 | Non lesion PsA vs Lesion PsA GSE186063 |

| | | | |
|---|---|---|---|
| CD38 | PAX4 | 122 | Non lesion PsA vs Lesion PsA GSE186063 |
| GBP1 | JUN | 122 | Non lesion PsA vs Lesion PsA GSE186063 |
| ICOS | SMAD2::SMAD3::SMAD4 | 122 | Non lesion PsA vs Lesion PsA GSE186063 |
| CXCR2 | FOXP2 | 122 | Non lesion PsA vs Lesion PsA GSE186063 |
| TYMP | NR1H2::RXRA | 121 | Non lesion PsA vs Lesion PsA GSE186063 |
| FRZB | DUX4 | 121 | Non lesion PsA vs Lesion PsA GSE186063 |
| ACP7 | FOXG1 | 121 | Non lesion PsA vs Lesion PsA GSE186063 |
| NKG7 | BARHL2 | 121 | Non lesion PsA vs Lesion PsA GSE186063 |
| ACKR2 | RXRA::VDR | 120 | Non lesion PsA vs Lesion PsA GSE186063 |
| GPT | NR2C2 | 120 | Non lesion PsA vs Lesion PsA GSE186063 |

| | | | |
|---|---|---|---|
| FOSL1 | MYF6 | 120 | Non lesion PsA vs Lesion PsA GSE186063 |
| IL1B | IRF1 | 119 | Non lesion PsA vs Lesion PsA GSE186063 |
| HERC6 | FLI1 | 119 | Non lesion PsA vs Lesion PsA GSE186063 |
| KANK4 | RELA | 117 | Non lesion PsA vs Lesion PsA GSE186063 |
| S100A12 | HNF1B | 117 | Non lesion PsA vs Lesion PsA GSE186063 |
| NAMPT | FOXD1 | 116 | Non lesion PsA vs Lesion PsA GSE186063 |
| CALD1 | MSC | 116 | Non lesion PsA vs Lesion PsA GSE186063 |
| IFIT3 | ELK4 | 115 | Non lesion PsA vs Lesion PsA GSE186063 |
| IFNG | ESR2 | 115 | Non lesion PsA vs Lesion PsA GSE186063 |
| CXCL10 | KLF5 | 114 | Non lesion PsA vs Lesion PsA GSE186063 |

| | | | |
|---|---|---|---|
| ZC3H12A | MSX1 | 114 | Non lesion PsA vs Lesion PsA GSE186063 |
| MX1 | ALX3 | 113 | Non lesion PsA vs Lesion PsA GSE186063 |
| FCGR1A | NFATC2 | 112 | Non lesion PsA vs Lesion PsA GSE186063 |
| ZNF483 | REL | 111 | Non lesion PsA vs Lesion PsA GSE186063 |
| S100A9 | HNF1A | 109 | Non lesion PsA vs Lesion PsA GSE186063 |
| CSMD1 | KLF13 | 109 | Non lesion PsA vs Lesion PsA GSE186063 |
| STAT1 | MIXL1 | 107 | Non lesion PsA vs Lesion PsA GSE186063 |
| S100P | HOXC11 | 105 | Non lesion PsA vs Lesion PsA GSE186063 |
| SLC1A2 | MEF2B | 104 | Non lesion PsA vs Lesion PsA GSE186063 |
| CD8A | NKX6-1 | 103 | Non lesion PsA vs Lesion PsA GSE186063 |

| | | | |
|---|---|---|---|
| CXCL13 | IRF2 | 102 | Non lesion PsA vs Lesion PsA GSE186063 |
| CCL2 | HSF1 | 102 | Non lesion PsA vs Lesion PsA GSE186063 |
| CLEC4D | ZBTB33 | 101 | Non lesion PsA vs Lesion PsA GSE186063 |
| AIM2 | OLIG2 | 100 | Non lesion PsA vs Lesion PsA GSE186063 |
| EGFR | ZNF354C | 90 | Non lesion PsA vs Lesion PsA GSE186063 |
| CD80 | NKX6-2 | 86 | Non lesion PsA vs Lesion PsA GSE186063 |
| OLR1 | EN1 | 140 | Normal Skin AS vs Lesion PsA GSE186063 |
| ZNF316 | FOXF2 | 162 | Normal Skin AS vs Lesion PsA GSE186063 |
| NAMPT | FOXD1 | 118 | Normal Skin AS vs Lesion PsA GSE186063 |
| PPARG | FOXL1 | 157 | Normal Skin AS vs Lesion PsA GSE186063 |

| | | | |
|---|---|---|---|
| SMOX | FOXI1 | 130 | Normal Skin AS vs Lesion PsA GSE186063 |
| PTGS2 | HNF1A | 130 | Normal Skin AS vs Lesion PsA GSE186063 |
| ZC3H12A | NHLH1 | 110 | Normal Skin AS vs Lesion PsA GSE186063 |
| CLEC4D | IRF1 | 101 | Normal Skin AS vs Lesion PsA GSE186063 |
| SPON2 | IRF2 | 126 | Normal Skin AS vs Lesion PsA GSE186063 |
| CXCL13 | MZF1(var.2) | 97 | Normal Skin AS vs Lesion PsA GSE186063 |
| GJB6 | MAX::MYC | 176 | Normal Skin AS vs Lesion PsA GSE186063 |
| FAXDC2 | PPARG | 149 | Normal Skin AS vs Lesion PsA GSE186063 |
| RSAD2 | PAX4 | 140 | Normal Skin AS vs Lesion PsA GSE186063 |
| PPARGC1A | PAX6 | 131 | Normal Skin AS vs Lesion PsA GSE186063 |

| | | | |
|---|---|---|---|
| AKAP13 | PBX1 | 134 | Normal Skin AS vs Lesion PsA GSE186063 |
| IL17F | RORA | 152 | Normal Skin AS vs Lesion PsA GSE186063 |
| GBP5 | RORA(var.2) | 134 | Normal Skin AS vs Lesion PsA GSE186063 |
| MUCL1 | RREB1 | 149 | Normal Skin AS vs Lesion PsA GSE186063 |
| NOXRED1 | RXRA::VDR | 120 | Normal Skin AS vs Lesion PsA GSE186063 |
| GZMB | ELK4 | 128 | Normal Skin AS vs Lesion PsA GSE186063 |
| CCR7 | SOX9 | 198 | Normal Skin AS vs Lesion PsA GSE186063 |
| CD38 | SRF | 122 | Normal Skin AS vs Lesion PsA GSE186063 |
| ACKR2 | SRY | 121 | Normal Skin AS vs Lesion PsA GSE186063 |
| EFCAB13 | TAL1::TCF3 | 159 | Normal Skin AS vs Lesion PsA GSE186063 |

| | | | |
|---|---|---|---|
| IL1B | YY1 | 120 | Normal Skin AS vs Lesion PsA GSE186063 |
| CTLA4 | REL | 136 | Normal Skin AS vs Lesion PsA GSE186063 |
| CYP1A1 | RELA | 131 | Normal Skin AS vs Lesion PsA GSE186063 |
| ZNF483 | NR1H2::RXRA | 115 | Normal Skin AS vs Lesion PsA GSE186063 |
| KANK4 | NFIC::TLX1 | 118 | Normal Skin AS vs Lesion PsA GSE186063 |
| TYMP | ZNF354C | 124 | Normal Skin AS vs Lesion PsA GSE186063 |
| ADAMTS9 | HINFP | 159 | Normal Skin AS vs Lesion PsA GSE186063 |
| NOS2 | ELF5 | 134 | Normal Skin AS vs Lesion PsA GSE186063 |
| IFIT3 | STAT1 | 115 | Normal Skin AS vs Lesion PsA GSE186063 |
| SERPINB1 | REST | 144 | Normal Skin AS vs Lesion PsA GSE186063 |

| | | | |
|---|---|---|---|
| EGF | CTCF | 126 | Normal Skin AS vs Lesion PsA GSE186063 |
| IGF2 | GATA1::TAL1 | 114 | Normal Skin AS vs Lesion PsA GSE186063 |
| IFNG | STAT3 | 114 | Normal Skin AS vs Lesion PsA GSE186063 |
| EGFR | EWSR1-FLI1 | 90 | Normal Skin AS vs Lesion PsA GSE186063 |
| ALB | NFATC2 | 131 | Normal Skin AS vs Lesion PsA GSE186063 |
| IL21 | HNF1B | 137 | Normal Skin AS vs Lesion PsA GSE186063 |
| FCGR1A | INSM1 | 116 | Normal Skin AS vs Lesion PsA GSE186063 |
| SAMD9 | FOXO3 | 161 | Normal Skin AS vs Lesion PsA GSE186063 |
| FCGR3A | RARA::RXRA | 160 | Normal Skin AS vs Lesion PsA GSE186063 |
| DKK1 | NR4A2 | 167 | Normal Skin AS vs Lesion PsA GSE186063 |

| | | | |
|---|---|---|---|
| CRB1 | PLAG1 | 136 | Normal Skin AS vs Lesion PsA GSE186063 |
| NRG1 | ESR2 | 108 | Normal Skin AS vs Lesion PsA GSE186063 |
| FOS | ARNT::HIF1A | 138 | Normal Skin AS vs Lesion PsA GSE186063 |
| HYAL4 | DUX4 | 136 | Normal Skin AS vs Lesion PsA GSE186063 |
| PI3 | FLI1 | 174 | Normal Skin AS vs Lesion PsA GSE186063 |
| SLC7A11 | FOS | 144 | Normal Skin AS vs Lesion PsA GSE186063 |
| FRZB | FOSL2 | 122 | Normal Skin AS vs Lesion PsA GSE186063 |
| FUT2 | FOXH1 | 169 | Normal Skin AS vs Lesion PsA GSE186063 |
| RORC | HSF1 | 154 | Normal Skin AS vs Lesion PsA GSE186063 |
| IFI6 | JUN | 168 | Normal Skin AS vs Lesion PsA GSE186063 |

| | | | |
|---|---|---|---|
| GATA3 | JUN(var.2) | 127 | Normal Skin AS vs Lesion PsA GSE186063 |
| GBP1 | JUND(var.2) | 123 | Normal Skin AS vs Lesion PsA GSE186063 |
| IL36RN | MEF2C | 135 | Normal Skin AS vs Lesion PsA GSE186063 |
| GJB2 | MAF::NFE2 | 139 | Normal Skin AS vs Lesion PsA GSE186063 |
| IL37 | NR2C2 | 146 | Normal Skin AS vs Lesion PsA GSE186063 |
| PLA2G4D | NRF1 | 153 | Normal Skin AS vs Lesion PsA GSE186063 |
| GPT | POU2F2 | 119 | Normal Skin AS vs Lesion PsA GSE186063 |
| CD274 | SMAD2::SMAD3::SMAD4 | 129 | Normal Skin AS vs Lesion PsA GSE186063 |
| CXCL2 | STAT1::STAT2 | 175 | Normal Skin AS vs Lesion PsA GSE186063 |
| ICOS | TCF7L2 | 119 | Normal Skin AS vs Lesion PsA GSE186063 |

| | | | |
|---|---|---|---|
| GZMA | ZBTB33 | 167 | Normal Skin AS vs Lesion PsA GSE186063 |
| IL2RA | FOXP2 | 134 | Normal Skin AS vs Lesion PsA GSE186063 |
| IL5 | SREBF1 | 167 | Normal Skin AS vs Lesion PsA GSE186063 |
| IL6 | SREBF2 | 148 | Normal Skin AS vs Lesion PsA GSE186063 |
| CXCL8 | THAP1 | 206 | Normal Skin AS vs Lesion PsA GSE186063 |
| CXCR2 | KLF5 | 123 | Normal Skin AS vs Lesion PsA GSE186063 |
| IL12B | DMRT3 | 154 | Normal Skin AS vs Lesion PsA GSE186063 |
| IL13 | FOXG1 | 156 | Normal Skin AS vs Lesion PsA GSE186063 |
| TNFRSF9 | LBX1 | 167 | Normal Skin AS vs Lesion PsA GSE186063 |
| IL17A | NFATC3 | 133 | Normal Skin AS vs Lesion PsA GSE186063 |

| | | | |
|---|---|---|---|
| CXCL10 | POU6F1 | 115 | Normal Skin AS vs Lesion PsA GSE186063 |
| IRS1 | SHOX | 143 | Normal Skin AS vs Lesion PsA GSE186063 |
| ITGA2B | ALX3 | 120 | Normal Skin AS vs Lesion PsA GSE186063 |
| ACP7 | BARHL2 | 121 | Normal Skin AS vs Lesion PsA GSE186063 |
| MEFV | BHLHE41 | 160 | Normal Skin AS vs Lesion PsA GSE186063 |
| MMP1 | CENPB | 129 | Normal Skin AS vs Lesion PsA GSE186063 |
| MMP9 | CREB3 | 193 | Normal Skin AS vs Lesion PsA GSE186063 |
| MX1 | DBP | 113 | Normal Skin AS vs Lesion PsA GSE186063 |
| NKG7 | ELF4 | 123 | Normal Skin AS vs Lesion PsA GSE186063 |
| IL22 | EN2 | 128 | Normal Skin AS vs Lesion PsA GSE186063 |

| | | | |
|---|---|---|---|
| NT5C3A | ESX1 | 137 | Normal Skin AS vs Lesion PsA GSE186063 |
| PER1 | ETV6 | 137 | Normal Skin AS vs Lesion PsA GSE186063 |
| HERC6 | GCM1 | 120 | Normal Skin AS vs Lesion PsA GSE186063 |
| PRF1 | GRHL1 | 163 | Normal Skin AS vs Lesion PsA GSE186063 |
| ZNF415 | GSC | 126 | Normal Skin AS vs Lesion PsA GSE186063 |
| RETN | HEY2 | 113 | Normal Skin AS vs Lesion PsA GSE186063 |
| CCND1 | HOXC11 | 184 | Normal Skin AS vs Lesion PsA GSE186063 |
| S100A8 | IRF8 | 141 | Normal Skin AS vs Lesion PsA GSE186063 |
| S100A9 | IRF9 | 107 | Normal Skin AS vs Lesion PsA GSE186063 |
| S100A12 | ISX | 116 | Normal Skin AS vs Lesion PsA GSE186063 |

| | | | |
|---|---|---|---|
| S100P | JDP2 | 106 | Normal Skin AS vs Lesion PsA GSE186063 |
| BGLAP | JDP2(var.2) | 138 | Normal Skin AS vs Lesion PsA GSE186063 |
| CCL20 | KLF13 | 234 | Normal Skin AS vs Lesion PsA GSE186063 |
| PRDM1 | LHX6 | 155 | Normal Skin AS vs Lesion PsA GSE186063 |
| SELL | MEF2B | 124 | Normal Skin AS vs Lesion PsA GSE186063 |
| NOD2 | MEOX1 | 137 | Normal Skin AS vs Lesion PsA GSE186063 |
| CSMD1 | MIXL1 | 111 | Normal Skin AS vs Lesion PsA GSE186063 |
| SLC1A2 | MLX | 103 | Normal Skin AS vs Lesion PsA GSE186063 |
| SOX4 | MLXIPL | 129 | Normal Skin AS vs Lesion PsA GSE186063 |
| STAT1 | MSC | 108 | Normal Skin AS vs Lesion PsA GSE186063 |

| | | | |
|---|---|---|---|
| SYT1 | MSX1 | 141 | Normal Skin AS vs Lesion PsA GSE186063 |
| TGFBR3 | MYF6 | 163 | Normal Skin AS vs Lesion PsA GSE186063 |
| TRAF5 | NEUROD2 | 102 | Normal Skin AS vs Lesion PsA GSE186063 |
| INS-IGF2 | NEUROG2 | 87 | Normal Skin AS vs Lesion PsA GSE186063 |
| RGPD6 | NFIA | 144 | Normal Skin AS vs Lesion PsA GSE186063 |
| CALD1 | NFIX | 117 | Normal Skin AS vs Lesion PsA GSE186063 |
| FOSL1 | NKX2-3 | 117 | Normal Skin AS vs Lesion PsA GSE186063 |
| CAPS2 | NKX2-8 | 128 | Normal Skin AS vs Lesion PsA GSE186063 |
| APOL1 | NKX6-1 | 129 | Normal Skin AS vs Lesion PsA GSE186063 |
| JRKL | NKX6-2 | 135 | Normal Skin AS vs Lesion PsA GSE186063 |

| | | | |
|---|---|---|---|
| HCAR3 | OLIG2 | 125 | Normal Skin AS vs Lesion PsA GSE186063 |
| CD8A | PAX7 | 104 | Normal Skin AS vs Lesion PsA GSE186063 |
| ADIPOQ | POU4F2 | 99 | Normal Skin AS vs Lesion PsA GSE186063 |
| CD80 | SP4 | 85 | Normal Skin AS vs Lesion PsA GSE186063 |
| AIM2 | SPDEF | 100 | Normal Skin AS vs Lesion PsA GSE186063 |
| CD36 | SPIC | 188 | Normal Skin AS vs Lesion PsA GSE186063 |
| DUSP4 | FOXF2 | 144 | PsA vs Healthy GSE117769 |
| CLEC4D | FOXD1 | 111 | PsA vs Healthy GSE117769 |
| IL6 | IRF2 | 148 | PsA vs Healthy GSE117769 |
| OLR1 | MZF1(var.2) | 144 | PsA vs Healthy GSE117769 |

| | | | |
|---|---|---|---|
| TNFAIP3 | MAX::MYC | 190 | PsA vs Healthy GSE117769 |
| CD69 | PPARG | 131 | PsA vs Healthy GSE117769 |
| GJB6 | FOXF2 | 173 | AS vs Healthy GSE117769 |
| CCR7 | IRF2 | 206 | AS vs Healthy GSE117769 |
| DDIT3 | MZF1(var.2) | 139 | AS vs Healthy GSE117769 |
| FOS | MAX::MYC | 152 | AS vs Healthy GSE117769 |
| CXCL2 | PPARG | 182 | AS vs Healthy GSE117769 |
| CXCL8 | PBX1 | 218 | AS vs Healthy GSE117769 |
| JUN | RORA | 142 | AS vs Healthy GSE117769 |
| AREG | RREB1 | 182 | AS vs Healthy GSE117769 |

| | | | |
|---|---|---|---|
| MYC | RXRA::VDR | 153 | AS vs Healthy GSE117769 |
| PER1 | SOX9 | 144 | AS vs Healthy GSE117769 |
| RORC | SRY | 170 | AS vs Healthy GSE117769 |
| TNFAIP3 | REL | 198 | AS vs Healthy GSE117769 |
| CCR6 | FOXF2 | 100 | PsA vs AS GSE117769 |
| CLEC4D | FOXD1 | 107 | PsA vs AS GSE117769 |
| MMP9 | IRF2 | 180 | PsA vs AS GSE117769 |
| RETN | MZF1(var.2) | 111 | PsA vs AS GSE117769 |
| SLC1A2 | TBXT | 107 | Non lesion PsA vs Lesion PsA GSE205748 |
| CXCL10 | EN1 | 115 | Non lesion PsA vs Lesion PsA GSE205748 |

| | | | |
|---|---|---|---|
| NAMPT | FOXF2 | 119 | Non lesion PsA vs Lesion PsA GSE205748 |
| TRIM22 | FOXD1 | 146 | Non lesion PsA vs Lesion PsA GSE205748 |
| LEP | FOXL1 | 118 | Non lesion PsA vs Lesion PsA GSE205748 |
| ACP7 | FOXI1 | 120 | Non lesion PsA vs Lesion PsA GSE205748 |
| MMP9 | HNF1A | 194 | Non lesion PsA vs Lesion PsA GSE205748 |
| S100A8 | NHLH1 | 139 | Non lesion PsA vs Lesion PsA GSE205748 |
| GPR35 | IRF1 | 126 | Non lesion PsA vs Lesion PsA GSE205748 |
| SPON2 | IRF2 | 126 | Non lesion PsA vs Lesion PsA GSE205748 |
| CXCL13 | MZF1(var.2) | 104 | Non lesion PsA vs Lesion PsA GSE205748 |
| GJB6 | MAX::MYC | 176 | Non lesion PsA vs Lesion PsA GSE205748 |

| | | | |
|---|---|---|---|
| FAXDC2 | PPARG | 149 | Non lesion PsA vs Lesion PsA GSE205748 |
| CCL20 | PAX4 | 231 | Non lesion PsA vs Lesion PsA GSE205748 |
| PPARGC1A | PAX6 | 135 | Non lesion PsA vs Lesion PsA GSE205748 |
| IL17F | PBX1 | 147 | Non lesion PsA vs Lesion PsA GSE205748 |
| GBP5 | RORA | 133 | Non lesion PsA vs Lesion PsA GSE205748 |
| MUCL1 | RORA(var.2) | 149 | Non lesion PsA vs Lesion PsA GSE205748 |
| JDP2 | RREB1 | 122 | Non lesion PsA vs Lesion PsA GSE205748 |
| SLC51B | RXRA::VDR | 133 | Non lesion PsA vs Lesion PsA GSE205748 |
| IL37 | ELK4 | 141 | Non lesion PsA vs Lesion PsA GSE205748 |
| CCR7 | SOX9 | 195 | Non lesion PsA vs Lesion PsA GSE205748 |

| | | | |
|---|---|---|---|
| PINK1 | SRF | 161 | Non lesion PsA vs Lesion PsA GSE205748 |
| ACKR2 | SRY | 121 | Non lesion PsA vs Lesion PsA GSE205748 |
| EFCAB13 | TAL1::TCF3 | 158 | Non lesion PsA vs Lesion PsA GSE205748 |
| ICOS | YY1 | 121 | Non lesion PsA vs Lesion PsA GSE205748 |
| CSF2 | REL | 136 | Non lesion PsA vs Lesion PsA GSE205748 |
| CMTM2 | RELA | 135 | Non lesion PsA vs Lesion PsA GSE205748 |
| IL23R | NR1H2::RXRA | 132 | Non lesion PsA vs Lesion PsA GSE205748 |
| CTLA4 | NFIC::TLX1 | 137 | Non lesion PsA vs Lesion PsA GSE205748 |
| CTSK | ZNF354C | 122 | Non lesion PsA vs Lesion PsA GSE205748 |
| MMP7 | HINFP | 137 | Non lesion PsA vs Lesion PsA GSE205748 |

| | | | |
|---|---|---|---|
| TNFSF10 | PDX1 | 185 | Non lesion PsA vs Lesion PsA GSE205748 |
| IL18 | ELF5 | 145 | Non lesion PsA vs Lesion PsA GSE205748 |
| GPT | STAT1 | 121 | Non lesion PsA vs Lesion PsA GSE205748 |
| KANK4 | REST | 116 | Non lesion PsA vs Lesion PsA GSE205748 |
| CYP1A1 | CTCF | 128 | Non lesion PsA vs Lesion PsA GSE205748 |
| CXCL2 | GATA1::TAL1 | 173 | Non lesion PsA vs Lesion PsA GSE205748 |
| CD274 | STAT3 | 120 | Non lesion PsA vs Lesion PsA GSE205748 |
| TGFBR3 | TFCP2 | 161 | Non lesion PsA vs Lesion PsA GSE205748 |
| ZNF483 | EWSR1-FLI1 | 113 | Non lesion PsA vs Lesion PsA GSE205748 |
| DDIT3 | NFATC2 | 140 | Non lesion PsA vs Lesion PsA GSE205748 |

| | | | |
|---|---|---|---|
| PLIN5 | HNF1B | 156 | Non lesion PsA vs Lesion PsA GSE205748 |
| DUSP4 | INSM1 | 137 | Non lesion PsA vs Lesion PsA GSE205748 |
| LYZ | FOXO3 | 138 | Non lesion PsA vs Lesion PsA GSE205748 |
| TYMP | RARA::RXRA | 122 | Non lesion PsA vs Lesion PsA GSE205748 |
| EGF | NR4A2 | 127 | Non lesion PsA vs Lesion PsA GSE205748 |
| SERPINB1 | PLAG1 | 142 | Non lesion PsA vs Lesion PsA GSE205748 |
| PLA2G4D | ESR2 | 153 | Non lesion PsA vs Lesion PsA GSE205748 |
| ALB | ARNT::HIF1A | 129 | Non lesion PsA vs Lesion PsA GSE205748 |
| FCGR1A | DUX4 | 112 | Non lesion PsA vs Lesion PsA GSE205748 |
| KLRB1 | FLI1 | 160 | Non lesion PsA vs Lesion PsA GSE205748 |

| | | | |
|---|---|---|---|
| ABCD2 | FOS | 138 | Non lesion PsA vs Lesion PsA GSE205748 |
| DKK1 | FOSL2 | 164 | Non lesion PsA vs Lesion PsA GSE205748 |
| CRB1 | FOXH1 | 134 | Non lesion PsA vs Lesion PsA GSE205748 |
| NKG7 | HSF1 | 122 | Non lesion PsA vs Lesion PsA GSE205748 |
| FOS | JUN | 136 | Non lesion PsA vs Lesion PsA GSE205748 |
| HYAL4 | JUN(var.2) | 135 | Non lesion PsA vs Lesion PsA GSE205748 |
| SLC7A11 | JUND(var.2) | 147 | Non lesion PsA vs Lesion PsA GSE205748 |
| FRZB | MEF2C | 121 | Non lesion PsA vs Lesion PsA GSE205748 |
| FUT2 | MAF::NFE2 | 167 | Non lesion PsA vs Lesion PsA GSE205748 |
| IFI6 | NR2C2 | 169 | Non lesion PsA vs Lesion PsA GSE205748 |

| | | | |
|---|---|---|---|
| PTPN22 | NRF1 | 113 | Non lesion PsA vs Lesion PsA GSE205748 |
| GATA3 | POU2F2 | 127 | Non lesion PsA vs Lesion PsA GSE205748 |
| GBP1 | SMAD2::SMAD3::SMAD4 | 122 | Non lesion PsA vs Lesion PsA GSE205748 |
| IL36RN | STAT1::STAT2 | 136 | Non lesion PsA vs Lesion PsA GSE205748 |
| GEM | TCF7L2 | 162 | Non lesion PsA vs Lesion PsA GSE205748 |
| GJB2 | ZBTB33 | 138 | Non lesion PsA vs Lesion PsA GSE205748 |
| TBX21 | FOXP2 | 139 | Non lesion PsA vs Lesion PsA GSE205748 |
| GZMA | SREBF1 | 162 | Non lesion PsA vs Lesion PsA GSE205748 |
| GZMB | SREBF2 | 125 | Non lesion PsA vs Lesion PsA GSE205748 |
| GZMK | THAP1 | 162 | Non lesion PsA vs Lesion PsA GSE205748 |

| | | | |
|---|---|---|---|
| IFI16 | KLF5 | 123 | Non lesion PsA vs Lesion PsA GSE205748 |
| IFIT3 | DMRT3 | 115 | Non lesion PsA vs Lesion PsA GSE205748 |
| IFNG | FOXG1 | 115 | Non lesion PsA vs Lesion PsA GSE205748 |
| IGF2 | LBX1 | 113 | Non lesion PsA vs Lesion PsA GSE205748 |
| IL1A | NFATC3 | 104 | Non lesion PsA vs Lesion PsA GSE205748 |
| IL1B | POU6F1 | 120 | Non lesion PsA vs Lesion PsA GSE205748 |
| IL2RA | SHOX | 135 | Non lesion PsA vs Lesion PsA GSE205748 |
| IL4 | ALX3 | 149 | Non lesion PsA vs Lesion PsA GSE205748 |
| CXCL8 | BARHL2 | 205 | Non lesion PsA vs Lesion PsA GSE205748 |
| CXCR2 | BHLHE41 | 122 | Non lesion PsA vs Lesion PsA GSE205748 |

| | | | |
|---|---|---|---|
| IL10 | CENPB | 102 | Non lesion PsA vs Lesion PsA GSE205748 |
| IL12B | CREB3 | 151 | Non lesion PsA vs Lesion PsA GSE205748 |
| TNFRSF9 | DBP | 164 | Non lesion PsA vs Lesion PsA GSE205748 |
| IL17A | ELF4 | 133 | Non lesion PsA vs Lesion PsA GSE205748 |
| IRS1 | EN2 | 143 | Non lesion PsA vs Lesion PsA GSE205748 |
| ITGAL | ESX1 | 143 | Non lesion PsA vs Lesion PsA GSE205748 |
| JUND | ETV6 | 145 | Non lesion PsA vs Lesion PsA GSE205748 |
| MCAM | GCM1 | 151 | Non lesion PsA vs Lesion PsA GSE205748 |
| MEFV | GRHL1 | 161 | Non lesion PsA vs Lesion PsA GSE205748 |
| MMP1 | GSC | 131 | Non lesion PsA vs Lesion PsA GSE205748 |

| | | | |
|---|---|---|---|
| MMP3 | HEY2 | 100 | Non lesion PsA vs Lesion PsA GSE205748 |
| MX1 | HOXC11 | 113 | Non lesion PsA vs Lesion PsA GSE205748 |
| NOS2 | IRF8 | 131 | Non lesion PsA vs Lesion PsA GSE205748 |
| OLR1 | IRF9 | 142 | Non lesion PsA vs Lesion PsA GSE205748 |
| IL22 | ISX | 128 | Non lesion PsA vs Lesion PsA GSE205748 |
| SOST | JDP2 | 139 | Non lesion PsA vs Lesion PsA GSE205748 |
| IL23A | JDP2(var.2) | 132 | Non lesion PsA vs Lesion PsA GSE205748 |
| PER1 | KLF13 | 139 | Non lesion PsA vs Lesion PsA GSE205748 |
| PI3 | LHX6 | 173 | Non lesion PsA vs Lesion PsA GSE205748 |
| TOMM7 | MEF2B | 117 | Non lesion PsA vs Lesion PsA GSE205748 |

| | | | |
|---|---|---|---|
| PPARG | MEOX1 | 153 | Non lesion PsA vs Lesion PsA GSE205748 |
| XAF1 | MIXL1 | 134 | Non lesion PsA vs Lesion PsA GSE205748 |
| SAMD9 | MLX | 158 | Non lesion PsA vs Lesion PsA GSE205748 |
| HERC6 | MLXIPL | 117 | Non lesion PsA vs Lesion PsA GSE205748 |
| PRF1 | MSC | 164 | Non lesion PsA vs Lesion PsA GSE205748 |
| DDX60 | MSX1 | 152 | Non lesion PsA vs Lesion PsA GSE205748 |
| ZNF415 | MYF6 | 126 | Non lesion PsA vs Lesion PsA GSE205748 |
| IL21 | NEUROD2 | 134 | Non lesion PsA vs Lesion PsA GSE205748 |
| CCND1 | NEUROG2 | 183 | Non lesion PsA vs Lesion PsA GSE205748 |
| RORC | NFIA | 153 | Non lesion PsA vs Lesion PsA GSE205748 |

| | | | |
|---|---|---|---|
| RPL7 | NFIX | 161 | Non lesion PsA vs Lesion PsA GSE205748 |
| S100A9 | NKX2-3 | 106 | Non lesion PsA vs Lesion PsA GSE205748 |
| S100A12 | NKX2-8 | 115 | Non lesion PsA vs Lesion PsA GSE205748 |
| BGLAP | NKX6-1 | 139 | Non lesion PsA vs Lesion PsA GSE205748 |
| SCN1A | NKX6-2 | 133 | Non lesion PsA vs Lesion PsA GSE205748 |
| CCL2 | OLIG2 | 102 | Non lesion PsA vs Lesion PsA GSE205748 |
| NOD2 | PAX7 | 138 | Non lesion PsA vs Lesion PsA GSE205748 |
| IFIH1 | POU4F2 | 165 | Non lesion PsA vs Lesion PsA GSE205748 |
| SGK1 | SP4 | 149 | Non lesion PsA vs Lesion PsA GSE205748 |
| CSMD1 | SPDEF | 110 | Non lesion PsA vs Lesion PsA GSE205748 |

| | | | |
|---|---|---|---|
| BMP2 | SPIC | 136 | Non lesion PsA vs Lesion PsA GSE205748 |
| SOD2 | TBX2 | 139 | Non lesion PsA vs Lesion PsA GSE205748 |
| SPP1 | TBX20 | 113 | Non lesion PsA vs Lesion PsA GSE205748 |
| STAT1 | TBX21 | 108 | Non lesion PsA vs Lesion PsA GSE205748 |
| SYT1 | TFAP4 | 136 | Non lesion PsA vs Lesion PsA GSE205748 |
| INS-IGF2 | TFEB | 89 | Non lesion PsA vs Lesion PsA GSE205748 |
| CCR2 | ZBTB7B | 148 | Non lesion PsA vs Lesion PsA GSE205748 |
| RGPD6 | ZBTB7C | 142 | Non lesion PsA vs Lesion PsA GSE205748 |
| ZC3H12A | ZIC1 | 114 | Non lesion PsA vs Lesion PsA GSE205748 |
| FOSL1 | ZIC3 | 120 | Non lesion PsA vs Lesion PsA GSE205748 |

| | | | |
|---|---|---|---|
| APOL6 | ZBTB18 | 146 | Non lesion PsA vs Lesion PsA GSE205748 |
| SLC7A5 | LBX2 | 127 | Non lesion PsA vs Lesion PsA GSE205748 |
| EOMES | MEOX2 | 111 | Non lesion PsA vs Lesion PsA GSE205748 |
| EFCAB7 | MNX1 | 113 | Non lesion PsA vs Lesion PsA GSE205748 |
| IL1F10 | MSX2 | 123 | Non lesion PsA vs Lesion PsA GSE205748 |
| CAPS2 | NOTO | 125 | Non lesion PsA vs Lesion PsA GSE205748 |
| APOL1 | OTX1 | 131 | Non lesion PsA vs Lesion PsA GSE205748 |
| CCRL2 | PHOX2A | 201 | Non lesion PsA vs Lesion PsA GSE205748 |
| NMI | PITX3 | 165 | Non lesion PsA vs Lesion PsA GSE205748 |
| RSAD2 | PROP1 | 138 | Non lesion PsA vs Lesion PsA GSE205748 |

| | | | |
|---|---|---|---|
| CD3E | PRRX1 | 127 | Non lesion PsA vs Lesion PsA GSE205748 |
| CD8A | RAX2 | 104 | Non lesion PsA vs Lesion PsA GSE205748 |
| CD83 | RAX | 173 | Non lesion PsA vs Lesion PsA GSE205748 |
| ADIPOQ | RHOXF1 | 98 | Non lesion PsA vs Lesion PsA GSE205748 |
| CD28 | UNCX | 111 | Non lesion PsA vs Lesion PsA GSE205748 |
| CD80 | VAX1 | 85 | Non lesion PsA vs Lesion PsA GSE205748 |
| AIM2 | VAX2 | 101 | Non lesion PsA vs Lesion PsA GSE205748 |
| CD38 | VENTX | 122 | Non lesion PsA vs Lesion PsA GSE205748 |
| CD63 | VSX1 | 123 | Non lesion PsA vs Lesion PsA GSE205748 |
| CD69 | VSX2 | 128 | Non lesion PsA vs Lesion PsA GSE205748 |

| | | | |
|---|---|---|---|
| VIM | TBXT | 176 | Lesion PsA vs Control GSE205748 |
| MMP1 | EN1 | 131 | Lesion PsA vs Control GSE205748 |
| NAMPT | FOXF2 | 117 | Lesion PsA vs Control GSE205748 |
| TRIM22 | FOXD1 | 144 | Lesion PsA vs Control GSE205748 |
| OLR1 | FOXL1 | 139 | Lesion PsA vs Control GSE205748 |
| NOS2 | FOXI1 | 131 | Lesion PsA vs Control GSE205748 |
| PPARG | HNF1A | 153 | Lesion PsA vs Control GSE205748 |
| SELL | NHLH1 | 127 | Lesion PsA vs Control GSE205748 |
| GZMB | IRF1 | 125 | Lesion PsA vs Control GSE205748 |
| SPON2 | IRF2 | 127 | Lesion PsA vs Control GSE205748 |

| | | | |
|---|---|---|---|
| CXCL13 | MZF1(var.2) | 103 | Lesion PsA vs Control GSE205748 |
| GJB6 | MAX::MYC | 176 | Lesion PsA vs Control GSE205748 |
| PPARGC1A | PPARG | 135 | Lesion PsA vs Control GSE205748 |
| SOD2 | PAX4 | 137 | Lesion PsA vs Control GSE205748 |
| IL17F | PAX6 | 147 | Lesion PsA vs Control GSE205748 |
| GBP5 | PBX1 | 133 | Lesion PsA vs Control GSE205748 |
| MUCL1 | RORA | 148 | Lesion PsA vs Control GSE205748 |
| CCR7 | RORA(var.2) | 197 | Lesion PsA vs Control GSE205748 |
| ACKR2 | RREB1 | 120 | Lesion PsA vs Control GSE205748 |
| EFCAB13 | RXRA::VDR | 156 | Lesion PsA vs Control GSE205748 |

| | | | |
|---|---|---|---|
| ICOS | ELK4 | 121 | Lesion PsA vs Control GSE205748 |
| PPARGC1B | SOX9 | 138 | Lesion PsA vs Control GSE205748 |
| TRAF5 | SRF | 105 | Lesion PsA vs Control GSE205748 |
| CTLA4 | SRY | 137 | Lesion PsA vs Control GSE205748 |
| CTSK | TAL1::TCF3 | 123 | Lesion PsA vs Control GSE205748 |
| IFIT3 | YY1 | 115 | Lesion PsA vs Control GSE205748 |
| CYP1A1 | REL | 128 | Lesion PsA vs Control GSE205748 |
| ZNF483 | RELA | 112 | Lesion PsA vs Control GSE205748 |
| KANK4 | NR1H2::RXRA | 117 | Lesion PsA vs Control GSE205748 |
| DDIT3 | NFIC::TLX1 | 139 | Lesion PsA vs Control GSE205748 |

| | | | |
|---|---|---|---|
| TYMP | ZNF354C | 121 | Lesion PsA vs Control GSE205748 |
| TOMM7 | HINFP | 116 | Lesion PsA vs Control GSE205748 |
| CD36 | PDX1 | 181 | Lesion PsA vs Control GSE205748 |
| MEFV | ELF5 | 159 | Lesion PsA vs Control GSE205748 |
| HIF1A | STAT1 | 152 | Lesion PsA vs Control GSE205748 |
| ALB | REST | 131 | Lesion PsA vs Control GSE205748 |
| EGF | CTCF | 127 | Lesion PsA vs Control GSE205748 |
| IFI16 | GATA1::TAL1 | 122 | Lesion PsA vs Control GSE205748 |
| HK2 | STAT3 | 126 | Lesion PsA vs Control GSE205748 |
| EOMES | TFCP2 | 111 | Lesion PsA vs Control GSE205748 |

| | | | |
|---|---|---|---|
| SERPINB1 | EWSR1-FLI1 | 143 | Lesion PsA vs Control GSE205748 |
| FCGR1A | NFATC2 | 111 | Lesion PsA vs Control GSE205748 |
| XAF1 | HNF1B | 133 | Lesion PsA vs Control GSE205748 |
| FCGR3A | INSM1 | 155 | Lesion PsA vs Control GSE205748 |
| IL22 | FOXO3 | 128 | Lesion PsA vs Control GSE205748 |
| DKK1 | RARA::RXRA | 164 | Lesion PsA vs Control GSE205748 |
| CRB1 | NR4A2 | 133 | Lesion PsA vs Control GSE205748 |
| FOS | PLAG1 | 138 | Lesion PsA vs Control GSE205748 |
| TBX21 | ESR2 | 141 | Lesion PsA vs Control GSE205748 |
| HYAL4 | ARNT::HIF1A | 134 | Lesion PsA vs Control GSE205748 |

| | | | |
|---|---|---|---|
| SLC7A11 | DUX4 | 145 | Lesion PsA vs Control GSE205748 |
| MX1 | FLI1 | 113 | Lesion PsA vs Control GSE205748 |
| FRZB | FOS | 121 | Lesion PsA vs Control GSE205748 |
| FUT2 | FOSL2 | 166 | Lesion PsA vs Control GSE205748 |
| IFI6 | FOXH1 | 167 | Lesion PsA vs Control GSE205748 |
| HERC6 | HSF1 | 122 | Lesion PsA vs Control GSE205748 |
| PTPN22 | JUN | 113 | Lesion PsA vs Control GSE205748 |
| GATA3 | JUN(var.2) | 126 | Lesion PsA vs Control GSE205748 |
| GBP1 | JUND(var.2) | 121 | Lesion PsA vs Control GSE205748 |
| IL36RN | MEF2C | 135 | Lesion PsA vs Control GSE205748 |

| | | | |
|---|---|---|---|
| GEM | MAF::NFE2 | 158 | Lesion PsA vs Control GSE205748 |
| GJB2 | NR2C2 | 138 | Lesion PsA vs Control GSE205748 |
| IL37 | NRF1 | 140 | Lesion PsA vs Control GSE205748 |
| PLA2G4D | POU2F2 | 154 | Lesion PsA vs Control GSE205748 |
| GPR35 | SMAD2::SMAD3::SMAD4 | 126 | Lesion PsA vs Control GSE205748 |
| GPT | STAT1::STAT2 | 120 | Lesion PsA vs Control GSE205748 |
| CD274 | TCF7L2 | 122 | Lesion PsA vs Control GSE205748 |
| CXCL2 | ZBTB33 | 170 | Lesion PsA vs Control GSE205748 |
| IFNG | FOXP2 | 113 | Lesion PsA vs Control GSE205748 |
| IL1A | SREBF1 | 105 | Lesion PsA vs Control GSE205748 |

| | | | |
|---|---|---|---|
| IL1B | SREBF2 | 119 | Lesion PsA vs Control GSE205748 |
| IL2RA | THAP1 | 135 | Lesion PsA vs Control GSE205748 |
| IL4 | KLF5 | 149 | Lesion PsA vs Control GSE205748 |
| CXCL8 | DMRT3 | 207 | Lesion PsA vs Control GSE205748 |
| CXCR2 | FOXG1 | 122 | Lesion PsA vs Control GSE205748 |
| IL10 | LBX1 | 100 | Lesion PsA vs Control GSE205748 |
| IL12B | NFATC3 | 153 | Lesion PsA vs Control GSE205748 |
| TNFRSF9 | POU6F1 | 160 | Lesion PsA vs Control GSE205748 |
| IL17A | SHOX | 133 | Lesion PsA vs Control GSE205748 |
| IL18 | ALX3 | 144 | Lesion PsA vs Control GSE205748 |

| | | | |
|---|---|---|---|
| CXCL10 | BARHL2 | 114 | Lesion PsA vs Control GSE205748 |
| JUND | BHLHE41 | 146 | Lesion PsA vs Control GSE205748 |
| KLRB1 | CENPB | 159 | Lesion PsA vs Control GSE205748 |
| ACP7 | CREB3 | 120 | Lesion PsA vs Control GSE205748 |
| LEP | DBP | 119 | Lesion PsA vs Control GSE205748 |
| LYZ | ELF4 | 137 | Lesion PsA vs Control GSE205748 |
| MMP3 | EN2 | 97 | Lesion PsA vs Control GSE205748 |
| MMP7 | ESX1 | 135 | Lesion PsA vs Control GSE205748 |
| MMP9 | ETV6 | 194 | Lesion PsA vs Control GSE205748 |
| PER1 | GCM1 | 138 | Lesion PsA vs Control GSE205748 |

| | | | |
|---|---|---|---|
| PI3 | GRHL1 | 168 | Lesion PsA vs Control GSE205748 |
| ACP5 | GSC | 161 | Lesion PsA vs Control GSE205748 |
| SMOX | HEY2 | 128 | Lesion PsA vs Control GSE205748 |
| SAMD9 | HOXC11 | 157 | Lesion PsA vs Control GSE205748 |
| PRF1 | IRF8 | 166 | Lesion PsA vs Control GSE205748 |
| DDX60 | IRF9 | 149 | Lesion PsA vs Control GSE205748 |
| ZNF415 | ISX | 125 | Lesion PsA vs Control GSE205748 |
| ADAMTS9 | JDP2 | 158 | Lesion PsA vs Control GSE205748 |
| IL21 | JDP2(var.2) | 134 | Lesion PsA vs Control GSE205748 |
| CCND1 | KLF13 | 183 | Lesion PsA vs Control GSE205748 |

| | | | |
|---|---|---|---|
| RORC | LHX6 | 153 | Lesion PsA vs Control GSE205748 |
| RPL7 | MEF2B | 160 | Lesion PsA vs Control GSE205748 |
| RPL15 | MEOX1 | 154 | Lesion PsA vs Control GSE205748 |
| RPL41 | MIXL1 | 124 | Lesion PsA vs Control GSE205748 |
| RPS7 | MLX | 126 | Lesion PsA vs Control GSE205748 |
| RPS19 | MLXIPL | 138 | Lesion PsA vs Control GSE205748 |
| RPS21 | MSC | 130 | Lesion PsA vs Control GSE205748 |
| S100A8 | MSX1 | 135 | Lesion PsA vs Control GSE205748 |
| S100A9 | MYF6 | 106 | Lesion PsA vs Control GSE205748 |
| S100A12 | NEUROD2 | 115 | Lesion PsA vs Control GSE205748 |

| | | | |
|---|---|---|---|
| BGLAP | NEUROG2 | 139 | Lesion PsA vs Control GSE205748 |
| CCL20 | NFIA | 237 | Lesion PsA vs Control GSE205748 |
| PRDM1 | NFIX | 154 | Lesion PsA vs Control GSE205748 |
| NOD2 | NKX2-3 | 139 | Lesion PsA vs Control GSE205748 |
| IFIH1 | NKX2-8 | 169 | Lesion PsA vs Control GSE205748 |
| CSMD1 | NKX6-1 | 108 | Lesion PsA vs Control GSE205748 |
| BMP2 | NKX6-2 | 138 | Lesion PsA vs Control GSE205748 |
| SLC1A2 | OLIG2 | 105 | Lesion PsA vs Control GSE205748 |
| SPP1 | PAX7 | 111 | Lesion PsA vs Control GSE205748 |
| STAT1 | POU4F2 | 108 | Lesion PsA vs Control GSE205748 |

| | | | |
|---|---|---|---|
| STAT3 | SP4 | 136 | Lesion PsA vs Control GSE205748 |
| SYT1 | SPDEF | 134 | Lesion PsA vs Control GSE205748 |
| TGFBR3 | SPIC | 160 | Lesion PsA vs Control GSE205748 |
| ZC3H12A | TBX2 | 113 | Lesion PsA vs Control GSE205748 |
| FOSL1 | TBX20 | 120 | Lesion PsA vs Control GSE205748 |
| APOL6 | TBX21 | 145 | Lesion PsA vs Control GSE205748 |
| SLC7A5 | TFAP4 | 126 | Lesion PsA vs Control GSE205748 |
| EFCAB7 | TFEB | 114 | Lesion PsA vs Control GSE205748 |
| IL1F10 | ZBTB7B | 126 | Lesion PsA vs Control GSE205748 |
| CAPS2 | ZBTB7C | 125 | Lesion PsA vs Control GSE205748 |

| | | | |
|---|---|---|---|
| APOL1 | ZIC1 | 130 | Lesion PsA vs Control GSE205748 |
| RSAD2 | ZIC3 | 137 | Lesion PsA vs Control GSE205748 |
| CD3E | ZBTB18 | 128 | Lesion PsA vs Control GSE205748 |
| OSMR | LBX2 | 141 | Lesion PsA vs Control GSE205748 |
| CD8A | MEOX2 | 103 | Lesion PsA vs Control GSE205748 |
| ADIPOQ | MNX1 | 97 | Lesion PsA vs Control GSE205748 |
| CD28 | MSX2 | 111 | Lesion PsA vs Control GSE205748 |
| CD80 | NOTO | 85 | Lesion PsA vs Control GSE205748 |
| AIM2 | OTX1 | 99 | Lesion PsA vs Control GSE205748 |
| CD38 | PHOX2A | 121 | Lesion PsA vs Control GSE205748 |

| | | | |
|---|---|---|---|
| CD63 | PITX3 | 124 | Lesion PsA vs Control GSE205748 |
| SPP1 | FOXF2 | 99 | Non lesion PsA vs Control GSE205748 |
| DYSF | FOXF2 | 153 | EA vs Control GSE221786 |

*This table shows the genes and their associated transcription factors, identified from gene expression analyses in different datasets. The "Occurrences" column indicates the frequency with which each gene-transcription factor association was observed or its significance in the studies evaluated. The "Datasets" detail the specific comparisons made, such as "Non-lesion PsA vs. Lesion PsA" (Psoriatic arthritis without lesion vs. with lesion) and "Normal Skin AS vs. Lesion PsA" (Normal skin of Ankylosing Spondylitis vs. PsA with lesion), as well as comparisons of PsA and AS with healthy individuals, indicating the relevance of these interactions in different disease states.*
